# Motor cortical inactivation reduces the gain of kinematic primitives in mice performing a hold-still center-out reach task

**DOI:** 10.1101/304907

**Authors:** Tejapratap Bollu, Samuel C. Whitehead, Nikil Prasad, Jackson Walker, Nitin Shyamkumar, Raghav Subramaniam, Brian Kardon, Itai Cohen, Jesse Heymann Goldberg

## Abstract

Motor sequences are constructed from primitives, hypothesized building blocks of movement, but mechanisms of primitive generation remain unclear. Using automated homecage training and a novel forelimb sensor, we trained freely-moving mice to initiate forelimb sequences with clearly resolved submillimeter-scale micromovements followed by millimeter-scale reaches to learned spatial targets. Hundreds of thousands of trajectories were decomposed into millions of kinematic primitives, while closed-loop photoinhibition was used to test roles of motor cortical areas. Inactivation of contralateral motor cortex reduced primitive peak speed but, surprisingly, did not substantially affect primitive direction, initiation, termination, or complexity, resulting in isomorphic, spatially contracted trajectories that undershot targets. Our findings demonstrate separable loss of a single kinematic parameter, speed, and identify conditions where loss of cortical drive reduces the gain of motor primitives but does not affect their generation, timing or direction. The combination of high precision forelimb sensing with automated training and neural manipulation provides a system for studying how motor sequences are constructed from elemental building blocks.

## INTRODUCTION

An infamous problem in motor control is ‘the curse of dimensionality:’ a hand in motion sweeps through a near infinite continuum of possible trajectories, making motor control seem intractably high-dimensional (Bernstein, 1967; Shadmehr and Wise, 2005; Woodworth, 1899). One solution is to construct movement from a discrete set of elementary building blocks, or motion primitives (Flash and Hochner, 2005; Mussa-Ivaldi et al., 1994). For example, when you draw the letter “N” you carve a complex path through space, but “N” can be decomposed into three distinct strokes, or kinematic primitives, each with only a few parameters such as direction, speed and duration (Flash and Hogan, 1985; Milner, 1992; Viviani and Terzuolo, 1982)(Figure 1A-C). Motor primitives are abnormally generated and sequenced in disorders such as stroke, dystonia and Parkinson’s (Desmurget et al., 2004; Inzelberg et al., 1995; Majsak et al., 1998; Rohrer et al., 2004), yet precise circuits that generate primitives, determine their kinematics, and sequence them into a trajectory are poorly understood (Giszter, 2015).

**Figure 1.**
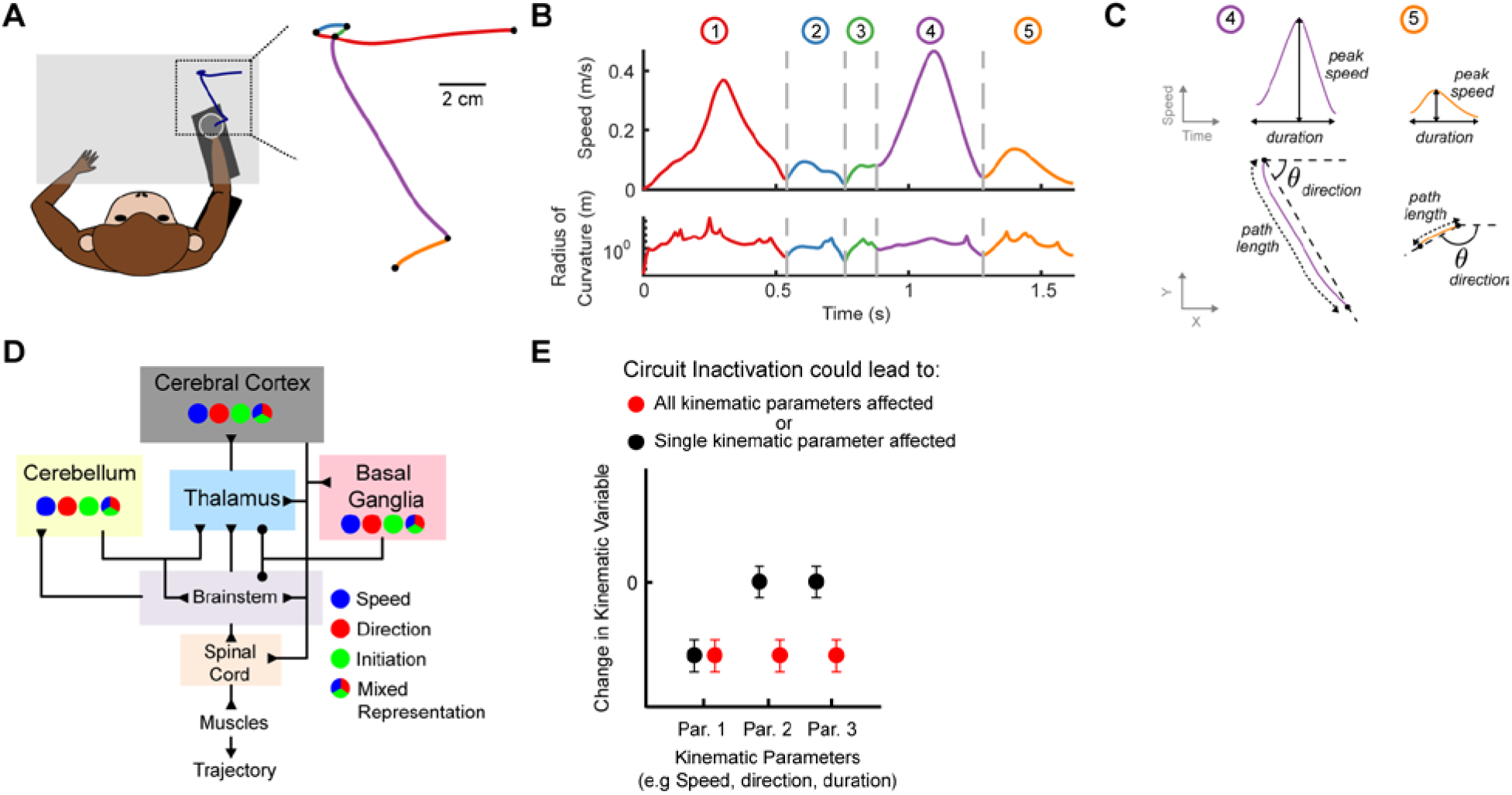
Primate hand trajectories can be decomposed into kinematic primitives. (A) Schematic of a hand trajectory during a sequential reach task (left) with primate kinematic data from a previous study (Gowda et al, 2015). Black dots denote boundaries separating discrete segments. (B). Speed (top) and radius of curvature (bottom) are plotted as a function of time in the trajectory from A. Segment boundaries (gray dashed lines) are detectable as temporally coincident minima. (C) The speed (top) and path (bottom) of the last two segments from the trajectory in A-B. Individual segments are described by kinematic parameters such as peak speed, duration, direction and pathlength. (D) Schematic of mammalian motor system, highlighting the existence of distributed kinematic representations. (E) Possible outcomes of inactivation experiments on primitive kinematics. Inactivation of a brain region could affect one (black) or multiple (red) kinematic variables.

One challenge is that kinematic representations and initiation signals are not regionally localized but are instead distributed throughout cortical, cerebellar, and basal ganglia circuits (Fortier et al., 1989; Fu et al., 1997; Schwartz, 2007; Scott, 2003; Shenoy et al., 2013; Turner and Anderson, 1997; Wong et al., 2014), complicating the identification of structure-function relationships and motivating causal experiments to provide at least some constraints on theories of motor control (Omrani et al., 2017; Pruszynski et al., 2011; Wolff and Ölveczky, 2018). For example, two extreme and opposite views of primitive generation are both compatible with distributed kinematic representations. Distinct kinematic parameters could differentially depend on distinct and separable neural circuits (Favilla et al., 1989; Flanders and Soechting, 1990), in which case a regional brain inactivation could affect one specific parameter (e.g. speed) and not others (e.g. direction, duration). Alternatively, kinematic parameters could be encoded interdependently (Shenoy et al., 2013), in which case a regional inactivation could affect multiple kinematic parameters at once (Figure 1D-E).

Genetic tools in mice enable temporally precise circuit manipulations (Guo et al., 2014), but forelimb kinematics have not been analyzed to extract primitives as in Figure 1, in part because trajectory segmentation requires sensitivity to resolve the rapid, tiny details of motion that occur at sharp turns. To address this issue, we designed ultra-low torque touch-sensing joysticks that resolve mouse forelimb kinematics with micron-millisecond spatiotemporal resolution and built an automated homecage system to train mice in a hold-still-center-out reach task. To complete the task, mice learned to first actively maintain the joystick in a small center position and then to produce an outward reach to learned spatial targets. The resultant trajectories carved complex paths in space. Algorithms previously used in primates were effective in decomposing forelimb trajectories into kinematic primitives, enabling us to test hypotheses of primitive generation. Inactivation of contralateral motor cortex reduced the peak speeds of primitives of all magnitudes, but did not affect their direction, initiation, duration, or complexity. As a result, forelimb trajectories exhibited isomorphic hypometria, i.e. they retained their basic shapes but were spatially contracted, or ‘shrunk,’ as if your letter “N” was a miniaturized “N”.

Methodologically, we demonstrate the utility of an automated system for high-throughput dissection of neural circuits that control basic building blocks of forelimb movement in mice. Conceptually, our findings identify conditions where the loss of cortical drive reduces the gain of motor primitives.

## RESULTS

### A novel sensor quantifies mouse forelimb kinematics with micron-millisecond spatiotemporal precision

Joysticks can be used in rodents to resolve forelimb kinematics during reach tasks (Francis and Chapin, 2004; Kimura et al., 2012; Mathis et al., 2017; Miri et al., 2017; Morandell and Huber, 2017; Panigrahi et al., 2015; Sanders and Kepecs, 2012; Slutzky et al., 2010; Wagner et al., 2017). To obtain raw trajectory data suitable for decomposition into primitives, we implemented a novel joystick design to increase spatial precision, reduce displacement force and ensure an isometric force profile. We designed a capacitive touch-sensing joystick that used contactless magnetic field sensing to detect motion, endowing it with micron-scale resolution (Figure 2 A-D, average spatial resolution: 320±35 nm, n=5 joysticks). We also replaced the standard two axis spring re-centering mechanism with a single pair of magnets, resulting in a stable, uniform force-displacement relationship and a 10-100 fold reduction in displacement force (Figure 2E, stiffness: 8.11 mN/mm (or 0.82 gf/mm), r2=0.99, see Methods). These modifications were necessary to resolve tiny details of mouse forelimb motion for trajectory decomposition, as outlined in more detail below.

**Figure 2.**
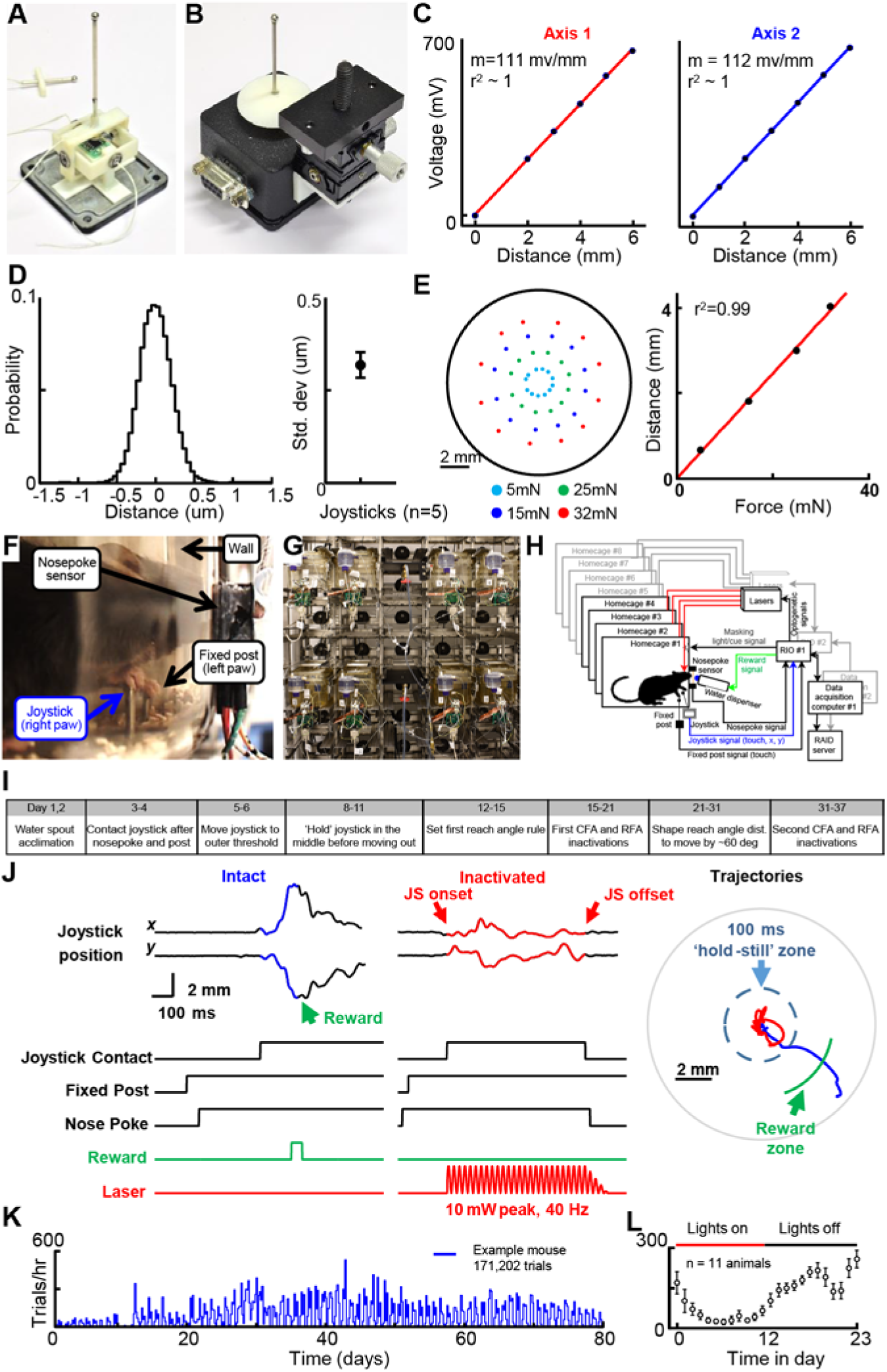
Automated homecage training of mice in a hold-still-center-out reach task. (A-E) Characterization of the joystick. (A-B) The joystick was a manipulandum capped with a capacitive touch sensor and mounted on a modified Gimbal assembly equipped with a 2-axis Hall sensor. (C-E) Hall sensor voltage measured as a function of distance across two non-orthogonal radial axes. (D) Spatial resolution characterization: measured probability distribution of a single joystick displacement for a single joystick (left) and mean±SEM standard deviation of the distribution across 5 joysticks (right). (E) Example of force-displacement characterization for a single joystick. Each dot indicates the measured displacement by a fixed force, color coded at bottom (left). Average joystick displacement as a function of force. (F) Still frame of mouse engaged in the task. (G) Photograph of homecage training room in mouse facility. (H) Schematic of signal processing pipeline for each homecage. (I) Timeline of behavioral shaping and photoinhibition experiments (J) Example trajectories and sensor data from a control (left) and CFAcl inactivated (middle) trial. (K) Trials per hour exhibited by an example animal for 80 days. (L) Trials per hour as a function of time in day (mean±SEM, n=11 animals).

To stabilize body posture, the joystick was integrated into a narrow ‘reward port’ consisting of five parts: (1) The joystick, which detects right paw contact and x-y movements; (2) A touch-sensing fixed post positioned for the left paw; (3) two side-walls that constrain the animal’s body position and orientation; (4) an IR-sensing nosepoke; and (5) a solenoid-controlled water dispensation spout within tongue’s reach of the nosepoke sensor. The requirement that animals engage the joystick, fixed post, and nosepoke contacts constrained the animal’s posture and ensured joystick manipulation with the right forepaw (Figure 2F, Movie S1).

### Automated homecage training of mice in a hold-still-center-out reach task

Automated training facilitates high-throughput experimentation on rodents in sophisticated learning tasks (Erlich et al., 2015; Murphy et al., 2016; Poddar et al., 2013; Woodard et al., 2017). We incorporated the joystick-reward port into rack-mountable, fully automated mouse homecages (Figure 2G-L). Mice enjoyed continuous, ad lib access to the joystick for their daily water, resulting in thousands of trajectories per animal per day (2792±362 trials per day, mean±sem, n=11 animals). We built a three-stage signal processing system pipeline to automate training and data acquisition: (1) an FPGA implemented millisecond timescale real-time analysis of all sensors, including joystick position, for closed-loop control of reward dispensation and of laser pulses for optogenetic experiments; (2) a real-time processor implemented second-timescale analysis for selective acquisition of trajectory and sensor data associated with eligible trials; (3) a host PC implemented day-timescale analysis of recent joystick manipulation patterns for automated contingency updates underlying training (Figure 2H, see Methods, Supplemental Methods).

A sequence of fully automated reward contingency updates shaped right forelimb trajectories in a direction-specific center-out reach task (Figure 2I, see Methods). First, mice were trained to contact the joystick after the nosepoke and fixed-post to ensure that joystick movement was attributable to right paw (Figure S1A,B). The timing of joystick contact became stereotyped with experience (Figure S1C,D; joystick contact onset: 184±14.5ms after nosepoke; entropy of JS contact time: 7.32±0.13 (day 1) vs 7.05±0.15 (criterion day), p<0.001, paired t-test). Next mice were trained to hold the joystick within an inner radius of 2 mm for 100 milliseconds prior to reaching past an outer radius of 4 mm (Figure 2J). This ‘hold period,’ defined as the latency from joystick contact to the moment of inner-radius transection, was implemented to study neural basis of maintaining stability and also to impose a delay in outward reaching that would allow for cortical photoinhibition to take effect before reach onset.

### Mice learn to hold still before reaching out

Due to the joystick’s low stiffness, the hold-in-center requirement approximates an inverted pendulum problem, in which a control policy is required to produce corrective micromovements to prevent rapid deviations from center position (Anderson, 1989; Bhounsule et al., 2015; Cabrera and Milton, 2002). Consistent with this, whereas early in training trajectories exhibited rapid displacements from the center position, later in training mice produced a clearly resolved sequence of ‘micromovements’ that maintained the joystick within the inner radius for longer periods of time (Figure 3 and Figure S2) (Mean hold time: 57±6 ms (first 100 trials) vs. 93 ms±14ms (at criterion), p<0.01, n=11 animals).

**Figure 3.**
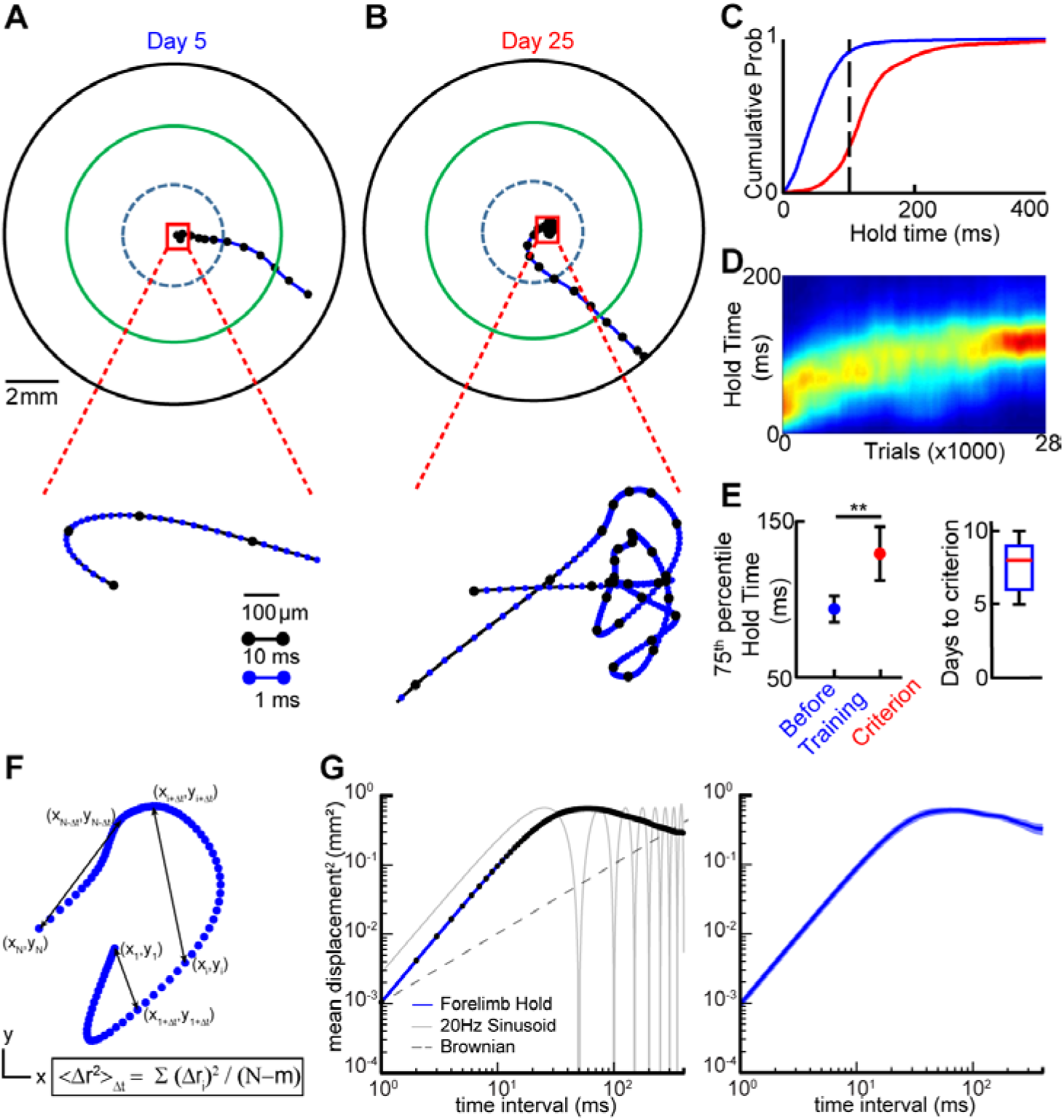
Mice learn hold-in-center requirement by producing sequential micromovements. (A) Example trajectory produced at day 5 since homecage introduction. Blue dashed circle indicates inner ‘hold’ region; green circle indicates reward zone; black circle indicates joystick boundaries. Black dots denote 10 millisecond intervals in the trajectory. Note distinct displacement scale bar for the expanded view of movement inside the hold region (bottom). (B) Data plotted as in A for a trajectory produced by the same mouse 20 days later. Expanded view at bottom plots details of micromovements produced to maintain hold requirement. (C) Cumulative probability of the hold time of trajectories before (blue) and after (red) training for the example mouse. (D) Heatmap showing the probability density plot of hold times as a function of trial number for the mouse show in in A-C. (E) 75^th^ percentile hold-time before training and at criterion for all mice (left), number of days to criterion (right) (data are mean±SEM across 11 animals). (F) Schematic describing calculation of the stabilogram diffusion function (SDF) of a short trajectory segment. (G) Example SDF from a single animal (left, blue line) and across all animals (right; n=7, line and shading represent mean+/-SEM of SDF). Example SDF’s of a purely Brownian motion (left, dashed gray) and a pure 20Hz sinusoid (left, solid gray) are shown for comparison.

To test if micromovements produced during the ‘hold’ reflected an active control policy for maintaining position, we computed stabilogram diffusion functions (SDFs). SDFs plot mean square displacement for all pairs of points in a trajectory as a function of time interval Δ*t* (Collins and De Luca, 1993; Peterka, 2000) (Methods, Figure 3H). The analysis effectively distinguishes between different types of motion. For example, the SDF of a classic random walk exhibits a slope of 1. Persistent motion biased towards increased time-dependent displacement exhibits a slope greater than 1. Finally, anti-persistent motion biased towards stabilizing position exhibits a slope less than 1 (Figure 3G). SDFs of trajectories acquired during the hold-period exhibited at least two regimes. At short latencies, the time-dependent displacement adhered to a power law (mean slope of log-log: 1.957±0.0028), reflecting a Levy flight process of rapid deviations consistent with the absence of a corrective process at those latencies. Yet a second mode appeared after a brief delay (mean inflection point: 27.59±1.56 ms), reflecting the onset of a distinct process that reversed time-dependent displacement (Figure 3G). SDFs acquired during human standing exhibit an identical shape, with a critical point reflecting the onset of an active control process to maintain balance (Peterka, 2002). These findings suggest that an active control policy, potentially similar to one used during the distinct inverted pendulum problem of maintaining of upright balance, was implemented to achieve the hold-still component of the task.

### Mice learn to reach in different rewarded directions

Once the contact, hold-still, and reach sequence was learned, all outward reach directions were rewarded, enabling each animal’s natural reach direction and variability to be quantified (Figure S3). Mice learned to execute the ‘contact’, ‘hold-still’ and reach sequence approximately one week after being placed into a homecage (Figure 3C-E, 8.5±1.05 days, n=11 animals, see Methods). Reach directions, defined as the angle at which the outer radius was transected, were next rotated clockwise (CW) or counterclockwise (CCW) by contingency updates that rewarded the 15th (for CW) or 85th (for CCW) percentile of their reach direction distribution (Figure 4). All mice learned to change their reach directions commensurate with rewarded contingency changes and changed their reach direction at a rate of 6.8 ± 2.3 (CW) and 4.6±1.46 (CCW) degrees/day (mean±sem) (Figure 4C-D).

**Figure 4.**
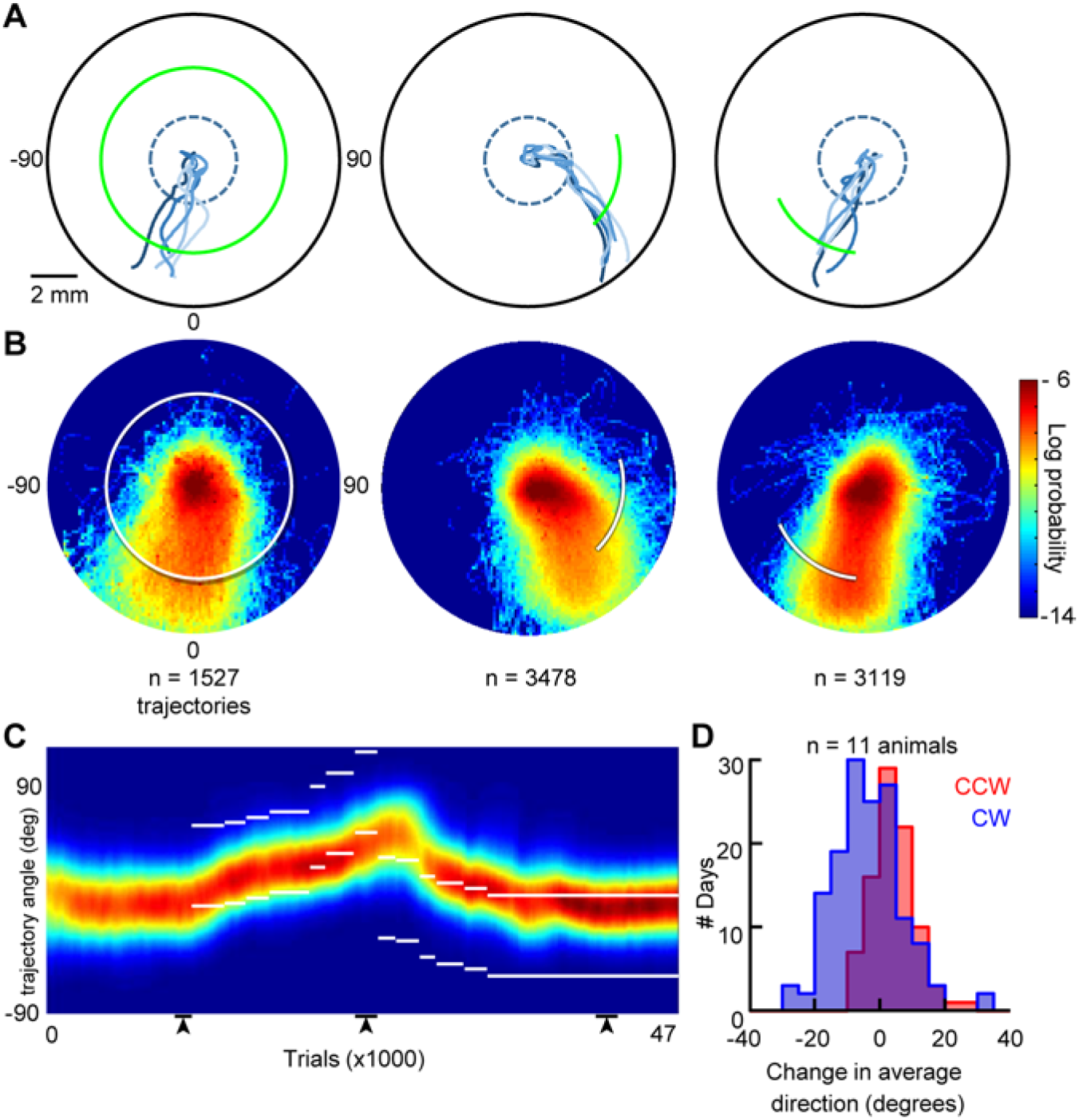
Mice learn to reach to spatial targets. (A) Example trajectories from three non-consecutive days (5 trajectories plotted per day). Green line indicates the reward zone. (B) 2D-probability distributions of all trajectories for the mouse on days from which trajectories were sampled from in (A), white bar indicates the reward zone. Number of trajectories denoted at bottom. (C) Probability distribution of reach direction as a function of trial number for an example mouse, white bars indicate the rewarded zone boundaries. Black bars at bottom indicate data from the three non-consecutive days plotted in A-B (left to right order preserved). (D) Histogram showing change in reach direction per day as reward zones were rotated clockwise (negative angle change, blue) or counter-clockwise (positive angle change, red).

### Roles of different motor cortical areas in holding still and reaching to target locations

By design, our task structure resolved many aspects of movement: from posture-maintaining micromovements for holding still in center to larger amplitude reaches aimed at learned spatial targets. We wondered how different motor cortical areas contribute to these processes. Suppressing an area required to stabilize posture or keep the brakes on movement will lead to an inability to hold still and a premature outward reach (Ebbesen and Brecht, 2017; Shadmehr, 2017; Velliste et al., 2014). Suppressing activity in a cortical area that promotes movement will reduce the probability of reaching out (Guo et al., 2015; Miri et al., 2017; Morandell and Huber, 2017; Peters et al., 2014). Suppressing activity in an area that controls reach direction could lead to inaccurate or highly variable movements (Mason et al., 1998). Finally, inactivating an area that is unrelated to forelimb movement should not affect performance.

To distinguish these outcomes, we used joystick contact-triggered photoinhibition on randomly interleaved trials, using methods in VGAT-hChr2 mice described previously (see Methods, Movie S2) (Guo et al., 2014). We targeted four motor cortical regions previously implicated in forelimb control (ipsi- and contralateral rostral and caudal forelimb areas of mouse motor cortex: CFAcl, CFAil, RFAcl and RFAil, see Methods)(Brown and Teskey, 2014; Harrison et al., 2012; Rouiller et al., 1993; Wang et al., 2017). In no case did a cortical inactivation impair the ability to execute the learned hold; in fact inactivation CFAcl significantly increased the probability of satisfying the hold criterion (Figure S2 and Table 1, p(hold=success) 0.38±0.07 (control) vs 0.49±0.09 (CFAcl Inactivated), p<0.05). During the hold, CFAcl inactivation significantly altered the slope (intact: 1.957+/-0.0028 vs inactivated: 1.966+/-0.0027, p<0.01 paired t-test), the inflection point (intact: 27.59+/-1.57ms vs inactivated: 32.45+/-1.77ms, p<0.001 paired t-test) and the offset (intact: 8.12e-04 mm^2^/s vs inactivated: 5.98e-04 mm^2^/s, p<0.01 paired t-test) of the stabilogram diffusion function (Figure 5).

**Figure 5.**
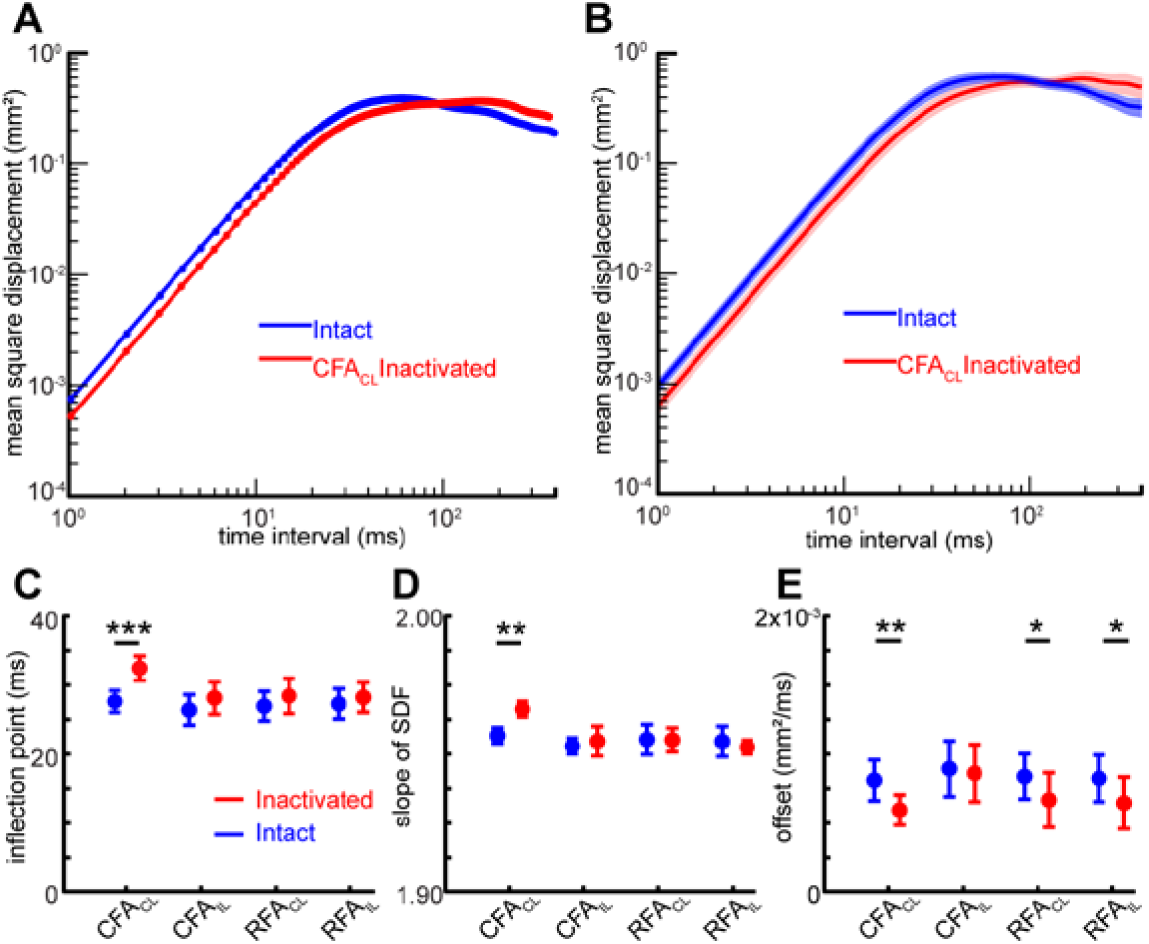
Effect of cortical inactivations on the stabilogram diffusion function (SDF) during the hold. (A) SDF of the hold component of the trajectories with CFAcl intact (blue) and inactivated from an example animal. (B) SDFs of trajectories with CFAcl intact and inactivated across animals (n=7) (C) mean±SEM of Inflection points, (D) mean±SEM of slopes, and (E) mean±SEM of offsets of SDF’s between intact and inactivated trials across various motor cortical areas. ***denotes p<0.001, **denotes p<0.01 and *denotes p<0.5, for a paired t-test (see Tables 2–4 for stats).

**Table 1:** Probability of a successful hold (LME)

| Table 1: Probability of a successful hold (LME) |  |  |  |  |
| --- | --- | --- | --- | --- |
| *Probability of successful hold~ 1 + Laser + (1 \| Mouse) + (1\|Mouse:Day)* |  |  |  |  |
|  | **Term** | **Estimate** | **Standard Error** | **pValue of ttest** |
| CFAcl | Intercept | 0.3815 | 0.0723 | 1.62E-05 |
|  | Laser | 0.1099 | 0.0457 | 0.0237 |
| CFAil | Intercept | 0.3826 | 0.0925 | 3.26E-04 |
|  | Laser | 0.0657 | 0.041 | 0.1213 |
| RFAcl | Intercept | 0.3742 | 0.0894 | 2.89E-04 |
|  | Laser | 0.0097 | 0.0466 | 0.8369 |
| RFAil | Intercept | 0.3764 | 0.0887 | 2.48E-04 |
|  | Laser | 0.0472 | 0.0475 | 0.3292 |

**Table 2:**
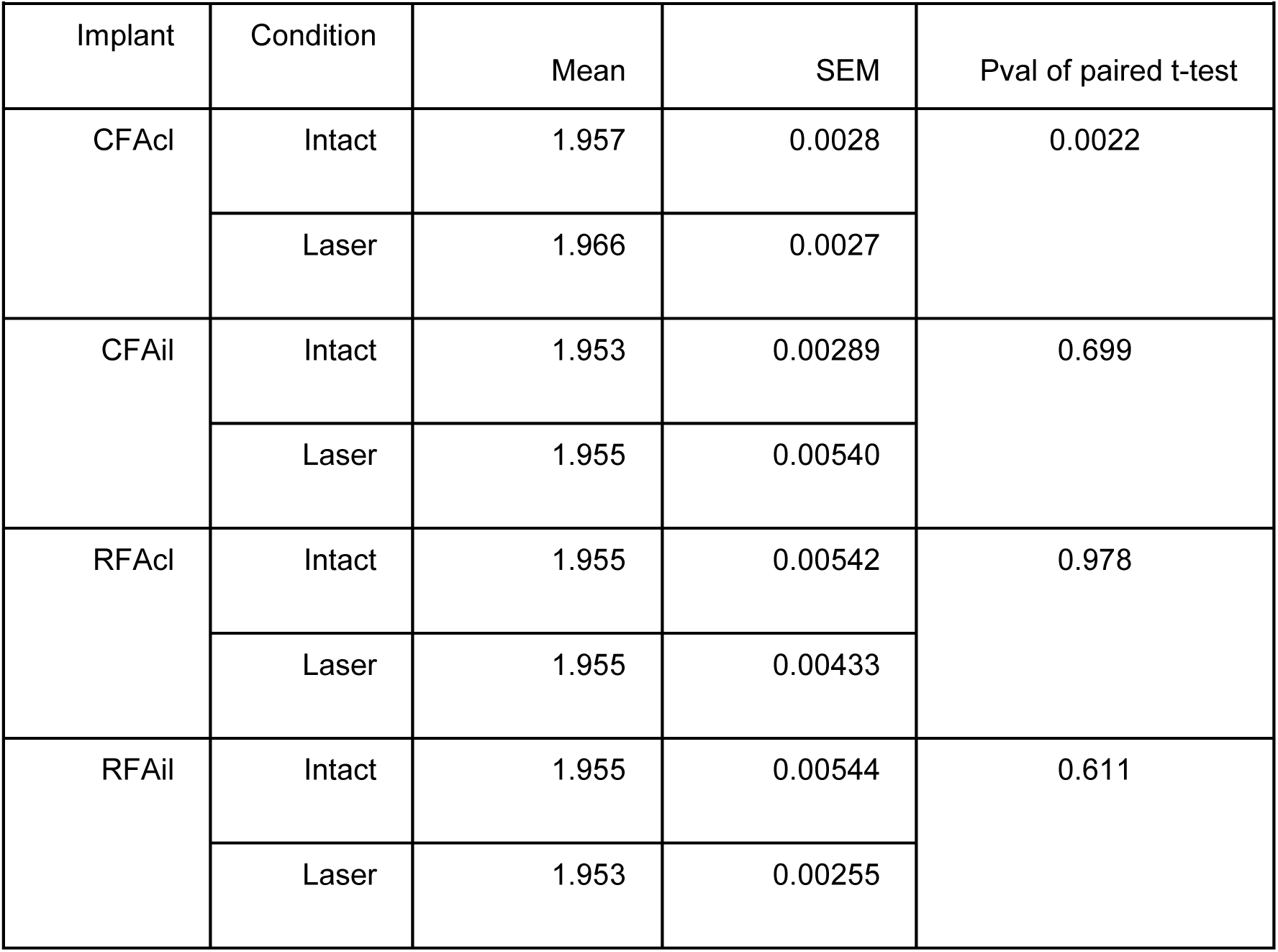
Slope of SDF function

**Table 3:** Inflection point of SDF function

| Table 3: Inflection point of SDF function |  |  |  |  |
| --- | --- | --- | --- | --- |
| Implant | Condition | Mean | SEM | Pval of paired t-test |
| CFAcl | Intact | 27.59786 | 1.568056 | 9.92E-05 |
|  | Laser | 32.4554 | 1.773085 |  |
| CFAil | Intact | 26.38059 | 2.217872 | 0.098023 |
|  | Laser | 28.12336 | 2.356636 |  |
| RFAcl | Intact | 26.92599 | 2.139888 | 0.146999 |
|  | Laser | 28.41068 | 2.538786 |  |
| RFAil | Intact | 27.26027 | 2.191394 | 0.322409 |
|  | Laser | 28.21111 | 2.175476 |  |

**Table 4:** Offsets of SDF function (log)

| Table 4: Offsets of SDF function (log) |  |  |  |  |
| --- | --- | --- | --- | --- |
| Implant | Condition | Mean | SEM | Pval of paired t-test |
| CFAcl | Intact | 8.12e-04 | 1.47e-04 | 0.0012 |
|  | Laser | 5.98e-04 | 1.09e-04 |  |
| CFAil | Intact | 8.96e-04 | 2.01e-04 | 0.340 |
|  | Laser | 8.63e-04 | 2.07e-04 |  |
| RFAcl | Intact | 8.4e-04 | 1.64e-04 | 0.021 |
|  | Laser | 6.7e-04 | 1.98e-04 |  |
| RFAil | Intact | 8.26e-04 | 1.69e-04 | 0.023 |
|  | Laser | 6.48e-04 | 1.85e-04 |  |

CFAcl inactivation also reduced peak trajectory speed and impaired outward reaching (Figure 6, Tables 5-6) (Peak trajectory speed: 115.98±7.82 mm/s (control) vs 94.92±8.03 (CFAcl), p<0.001; Probability that a trajectory would transect outer radius: Control: 67.42±25.4% vs. CFAcl: 48.02±3.43%, p<0.001, linear mixed effects (LME) models, n=7 animals, see Methods). In contrast, inactivation of CFAil or either RFA only modestly affected peak trajectory speed and did not affect the likelihood of reaching out (Figure 6, Tables 5-6.

**Figure 6.**
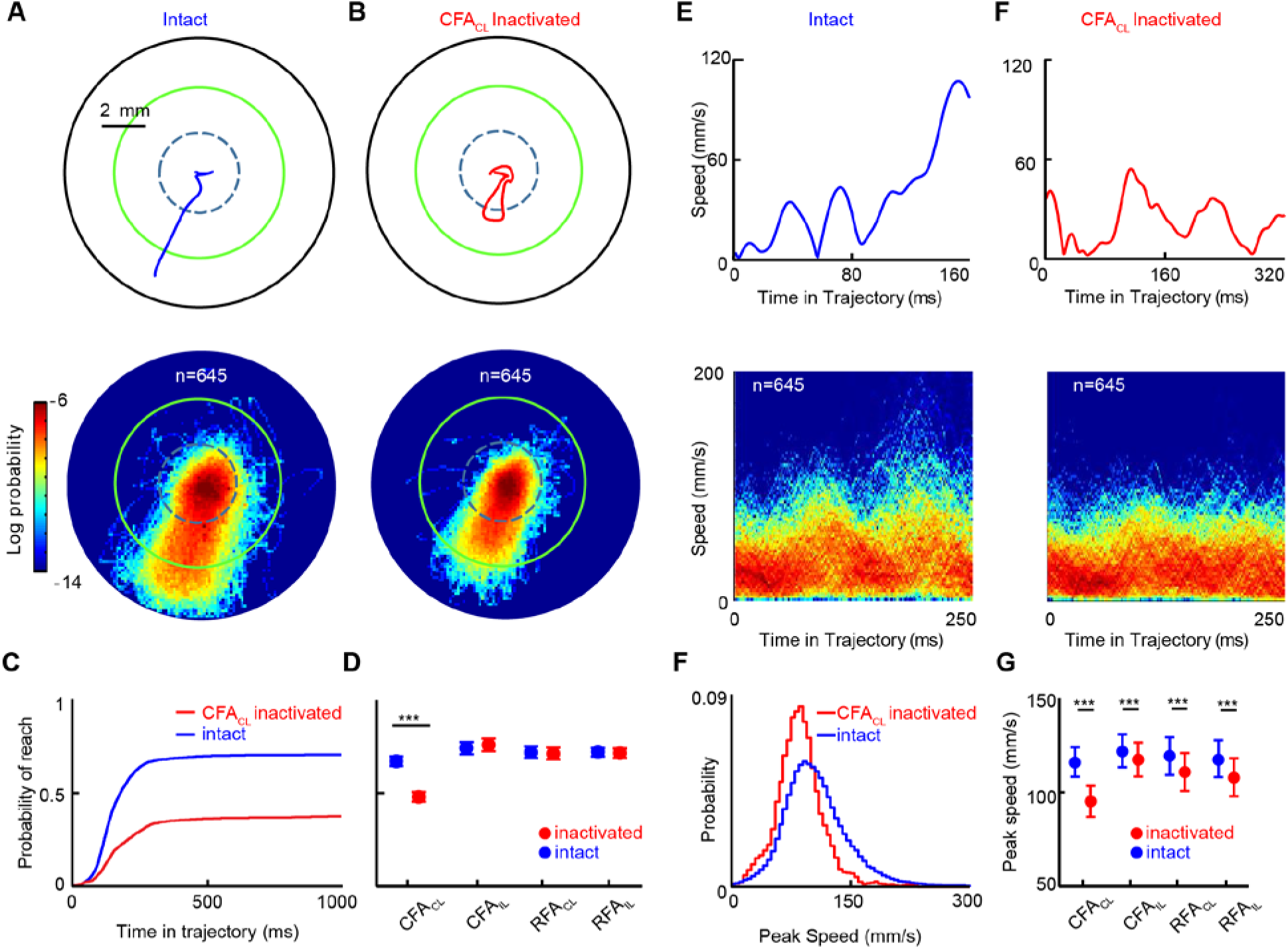
Effect of cortical inactivations on trajectory kinematics. (A-B) Example trajectories (top) and 2D spatial probability distributions (bottom) of trajectories during intact trials (A) and during randomly interleaved CFAcl-inactivated trials (B). Data in A-B are from same mouse and same day. (C) Cumulative probability that a trajectory would transect the outer radius as a function of time during control (blue) and CFAcl inactivated (red) trials. (D) Mean±SEM of probability of transecting the outer radius with or without inactivation of various motor cortical areas (n=7 mice). (E-F) Examples (top) and 2D distributions (bottom) of the speed profiles of the trajectories from (A-B). (G) Distributions of trajectory peak speed during the intact (blue) and CFAcl-inactivated (red) trials shown in A-B. (H) Mean±SEM of the peak speeds with or without inactivation of various motor cortical areas (n=7 mice). *** denotes p<0.001 LME (see Tables 5 and 6).

**Table 5:** Probability of transecting outer radius (4mm), LME model

| Table 5: Probability of transecting outer radius (4mm), LME model |  |  |  |  |
| --- | --- | --- | --- | --- |
| *Probability of transecting outer radius ~ 1 + Laser\* + (1 \| Mouse) + (1\|Mouse:Day)* |  |  |  |  |
|  | Term | Estimate | Standard Error | pValue of ttest |
| CFaCl | Intercept | 0.6706 | 0.0255 | 6.16E-40 |
|  | Laser | -0.1908 | 0.0239 | 1.17E-11 |
| CFaIl | Intercept | 0.7409 | 0.033 | 1.91E-26 |
|  | Laser | 0.018 | 0.0104 | 0.0914 |
| RFaCl | Intercept | 0.7179 | 0.0311 | 1.99E-34 |
|  | Laser | -0.0053 | 0.0073 | 0.4683 |
| RFaIl | Intercept | 0.71831499 | 0.022468451 | 4.60E-33 |
|  | Laser | -0.0039307 | 0.013062648 | 0.765 |

**Table 6:** Peak Speed of trajectories (mm/s), LME model

| Table 6: Peak Speed of trajectories (mm/s), LME model |  |  |  |  |
| --- | --- | --- | --- | --- |
| *Peak Speed ~ 1 + Laser + (1 \| Mouse) + (1\|Mouse:Day)* |  |  |  |  |
|  | Term | Estimate | Standard Error | pValue of ttest |
| CFAcl | Intercept | 115.9873 | 7.82 | 3.85E-24 |
|  | Laser | -21.0618 | 1.8242 | 2.17E-18 |
| CFAil | Intercept | 121.8371 | 8.798 | 5.06E-18 |
|  | Laser | -4.5188 | 1.5344 | 0.0051 |
| RFAcl | Intercept | 119.198 | 9.956 | 1.32E-18 |
|  | Laser | -8.4271 | 1.423 | 1.21E-07 |
| RFAil | Intercept | 117.8757 | 9.8914 | 1.16E-15 |
|  | Laser | -9.9938 | 1.574 | 8.66E-08 |

To test how cortical inactivations influenced reach direction, we inactivated each cortical site as mice reached to at least two different targets. Surprisingly, cortical inactivations did not significantly affect reach direction (Figure 7A-F, Table 7, p>0.05 all conditions, LME, see Methods). Inactivations also did not have a significant effect on the trial-to-trial variability of the direction of outward reaches (Figure 7G, Table 8, all conditions, p>0.05).

**Figure 7.**
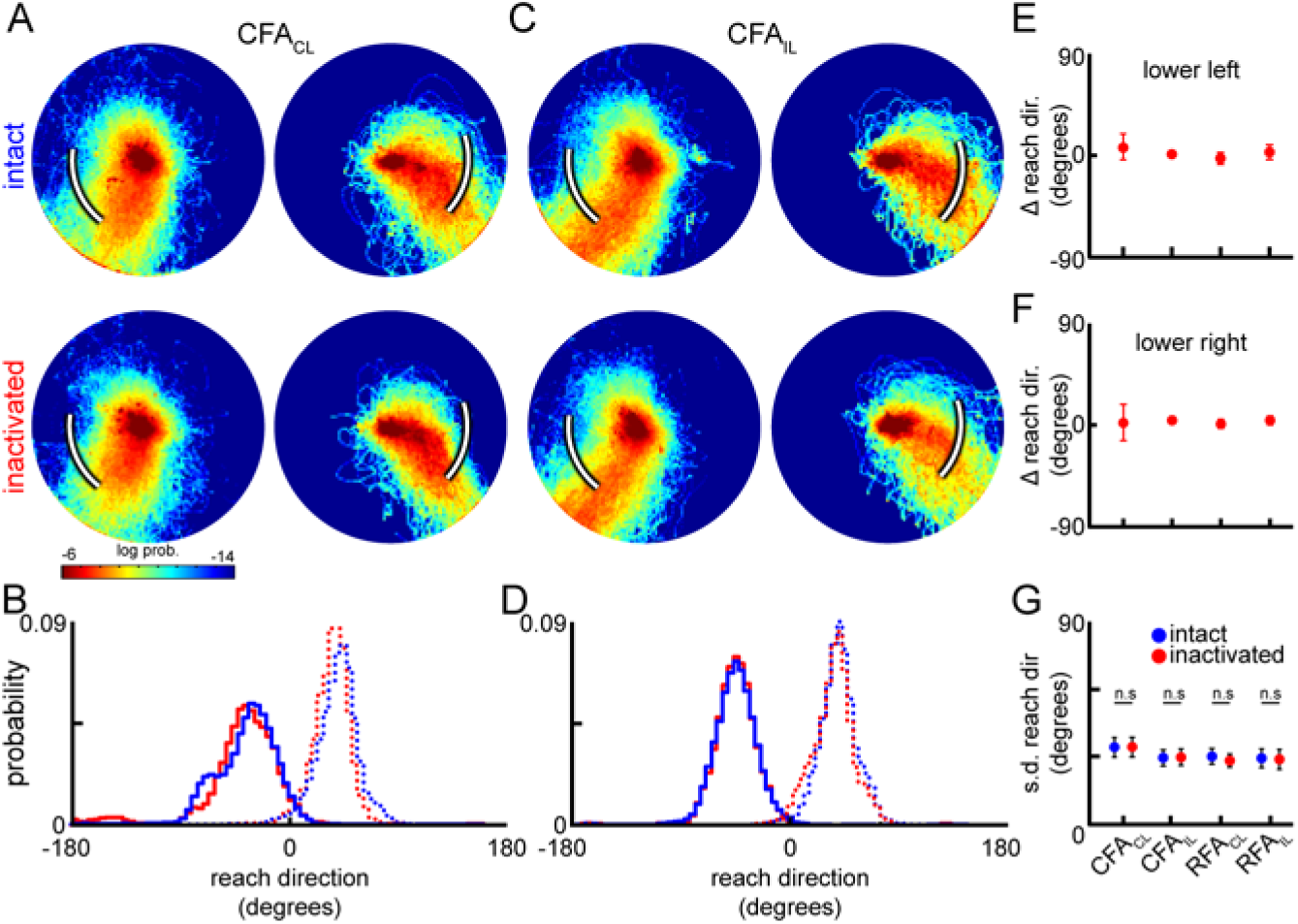
Effect of cortical inactivations on trajectory direction. (A) 2D distributions of trajectories from intact (top) and CFAcl inactivated (bottom) trials when an example mouse was reaching to different rewarded locations, indicated by white bars. (B) Reach direction distributions for intact (blue) and CFAcl inactivated (red) trajectories when the reward zone was lower left (solid lines) or lower right (dotted lines). (C-D) Data plotted as in A-B from a different day when CFAil was inactivated. (E-F) Mean±SEM of difference in reach angle between intact and inactivated trajectories across various motor cortical areas when the mice are reaching to lower left (E) or lower right (F). (G) mean±SEM in standard deviation of reach angle between intact and inactivated trials across various motor cortical areas (n=7 mice).

**Table 7:**
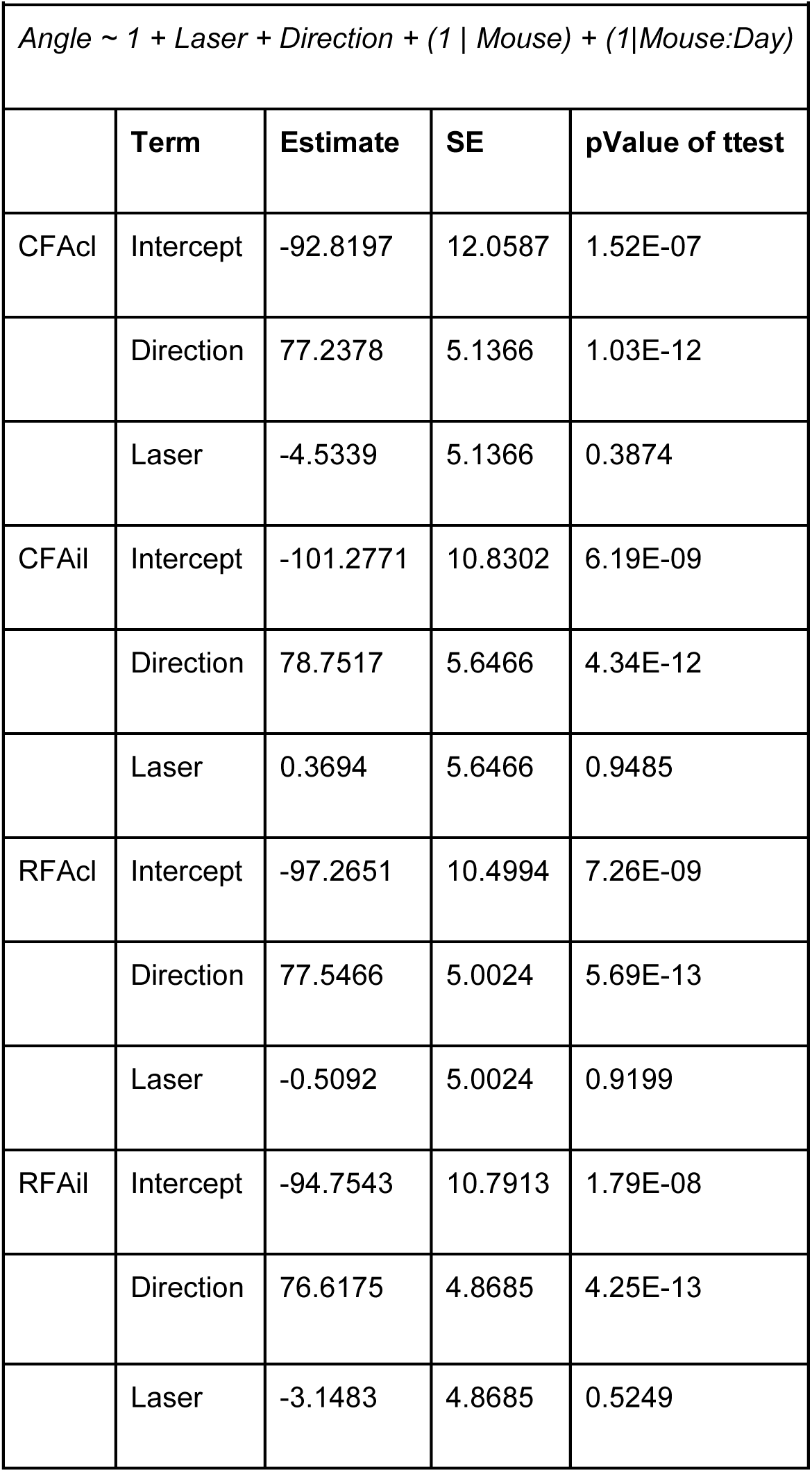
Angle of trajectory (degrees), LME model

**Table 8:** StdDev of Reach Angle for trajectory (degree)

| Table 8: StdDev of Reach Angle for trajectory (degree) |  |  |  |  |
| --- | --- | --- | --- | --- |
|  | Laser | Mean Std Dev of Angle | SEM | pValue of paired ttest |
| CFAcl | off | 33.97343034 | 4.37539 | p = 0.9945 |
|  | on | 33.96230309 | 4.35355 |  |
| CFAil | off | 29.93535776 | 3.52972 | p = 0.5823 |
|  | on | 27.98774658 | 2.91409 |  |
| RFAcl | off | 29.19536116 | 3.68252 | p = 0.8809 |
|  | on | 29.25396581 | 3.66252 |  |
| RFAil | off | 29.0139461 | 4.27275 | p = 0.4444 |
|  | on | 28.48927885 | 4.50755 |  |

### The roles of different cortical areas in controlling primitive kinematics

The joystick’s high spatiotemporal precision allowed us to analyze kinematic patterns of movement beyond the relatively coarse trajectory-level metrics described above. Trajectories exhibited complex, multi-peaked speed profiles, suggesting that they were composed of a sequence of discrete segments (Figures 1A-C and 7A-C). We tested the utility of two distinct decomposition algorithms commonly used in primate studies. One algorithm assumes that trajectories are derived from minimum-jerk basis functions with only three parameters: peak speed, duration, and the time in the sequence at which it is generated (see methods) (Gowda et al., 2015; Rohrer and Hogan, 2003; Viviani and Flash, 1995). This method was effective in decomposing mouse forelimb trajectories, in part because trajectories were composed of primate-like superpositions of bell-shaped velocity curves (Figure S4A-B), previously observed in mice (Azim et al., 2014; Mathis et al., 2017; Panigrahi et al., 2015). Following convention in primates we termed this class of kinematic primitives ‘submovements’. We also used a second decomposition algorithm that imposes segment boundaries at movement discontinuities revealed by temporally coincident minima in the radius of curvature and speed (Figure 1, Figure 7A-B) (Viviani and Terzuolo, 1982). This method essentially functions as a ‘sharp turn detector’ and identifies segments bounded by moments in a trajectory when a new force was imposed on the forelimb and, in a task that lacks external perturbations, also defines moments when an efferent neural command signal must have been generated by the central nervous system (Milner, 1992). We term this class of kinematic primitives ‘segments’.

Decomposition enabled each trajectory to be analyzed as a sequence of discrete primitives – each of which differed in kinematic parameters such as duration, complexity, speed, pathlength and direction (Figure 1C). Cortical inactivations could affect any or all of these parameters, and potentially in an amplitude-dependent way, i.e. differentially affecting hold-still versus reach components of the task. As expected, the durations, peak speeds, and pathlengths of primitives produced during hold-still periods were significantly smaller than during reaches (p<0.001, LME, Tables 9-11). Notably, the difference in segment durations between hold and reach was subtle compared to differences in segment speed and pathlength (ratio of reach versus hold value for duration: 1.86 ± 0.04; peak speed: 2.54 ± 0.19; path length: 4.19 ± 0.35, LME, Fig 7J-L). In fact, segment path length was more strongly predicted by its peak speed (R^2^=0.77 ± 0.04) than by its duration (0.58 ± 0.13, n=7 animals, Figure S5), suggesting an adherence to the isochrony principle, previously observed in primate reach and human handwriting tasks, in which a segment’s pathlength is more strongly predicted by its peak speed than by its duration (Viviani and Flash, 1995; Viviani and McCollum, 1983).

**Table 9:** Duration of segments (milliseconds), LME model

| Table 9: Duration of segments (milliseconds), LME model |  |  |  |  |
| --- | --- | --- | --- | --- |
| *Duration ~ 1 + Laser\*Condition + (1 \| Mouse) + (1\|Mouse:Day)* |  |  |  |  |
|  | **Model Term** | **Estimate** | **SE** | **pValue of ttest** |
| CFACl | Intercept | 34.057 | 2.8549 | 1.43E-23 |
|  | Laser (hold) | -1.5385 | 1.7015 | 0.36733 |
|  | Laser (reach) | -0.16667 | 1.7015 | 0.9221 |
|  | Condition (reach vs hold) | 28.41 | 1.7015 | 3.30E-36 |
|  | Laser:Condition (interaction) | 1.3718 | 2.4063 | 0.56946 |
| CFAil | Intercept | 35.563 | 3.6848 | 8.35E-16 |
|  | Laser (hold) | 0.14 | 2.622 | 0.95753 |
|  | Laser (reach) | 0.96 | 2.622 | 0.71507 |
|  | Condition (reach vs hold) | 29.56 | 2.622 | 2.82E-19 |
|  | Laser:Condition (interaction) | 0.82 | 3.7081 | 0.82545 |
| RFAcl | Intercept | 33.834 | 3.0082 | 1.86E-21 |
|  | Laser (hold) | -1.4595 | 1.7568 | 0.4075 |
|  | Laser (reach) | -0.25676 | 1.7568 | 0.88401 |
|  | Condition (reach vs hold) | 29.77 | 1.7568 | 4.15E-36 |
|  | Laser:Condition (interaction) | 1.2027 | 2.4846 | 0.62907 |
| RFAil | Intercept | 34.061 | 3.4471 | 6.31E-16 |
|  | Laser (hold) | 0.34783 | 2.7918 | 0.90113 |
|  | Laser (reach) | 2.5 | 2.7918 | 0.37297 |
|  | Condition (reach vs hold) | 31.065 | 2.7918 | 1.81E-18 |
|  | Laser:Condition (interaction) | 2.1522 | 3.9482 | 0.58706 |

**Table 10:**
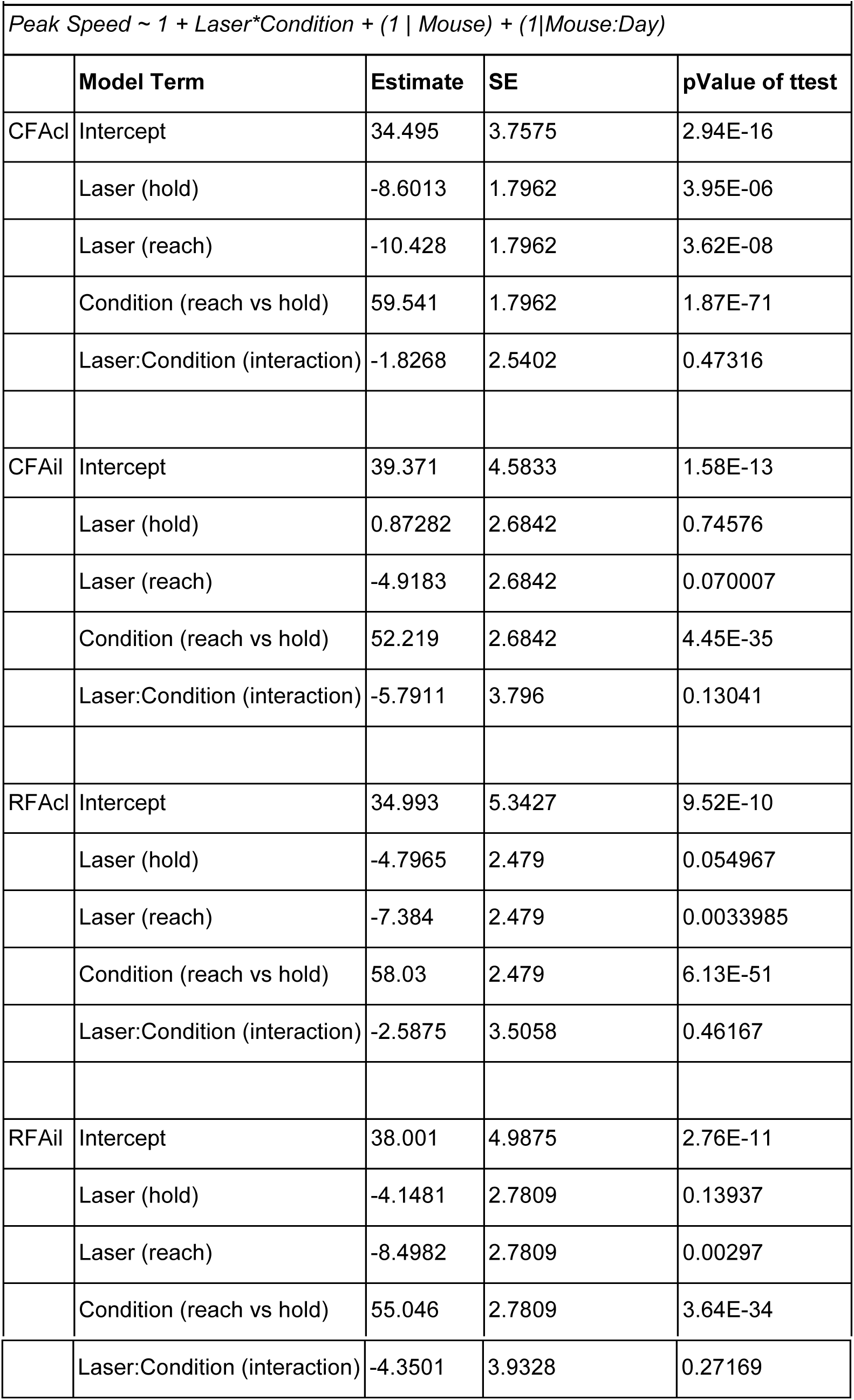
Peak speed of segments (mm/s), LME model

**Table 11:** Path length of segments (mm), LME model

| Table 11: Path length of segments (mm), LME model |  |  |  |  |
| --- | --- | --- | --- | --- |
| *Path Length ~ 1 + Laser\*Condition + (1 \| Mouse) + (1\|Mouse:Day)* |  |  |  |  |
|  | **Model Term** | **Estimate** | **SE** | **pValue of ttest** |
| CFACl | Intercept | 0.8393 | 0.15934 | 4.65E-07 |
|  | Laser (hold) | -0.22265 | 0.10844 | 0.041768 |
|  | Laser (reach) | -0.49128 | 0.10844 | 0.000011833 |
|  | Condition (reach vs hold) | 2.9877 | 0.10844 | 5.52E-61 |
|  | Laser:Condition (interaction) | -0.26863 | 0.15336 | 0.081844 |
| CFAil | Intercept | 1.0139 | 0.16868 | 3.32E-08 |
|  | Laser (hold) | 0.0081177 | 0.14906 | 0.95668 |
|  | Laser (reach) | -0.076229 | 0.14906 | 0.61026 |
|  | Condition (reach vs hold) | 2.7779 | 0.14906 | 1.17E-33 |
|  | Laser:Condition (interaction) | -0.084347 | 0.21081 | 0.68996 |
| RFAcl | Intercept | 0.8777 | 0.19372 | 0.000012245 |
|  | Laser (hold) | -0.1234 | 0.11648 | 0.2912 |
|  | Laser (reach) | -0.32152 | 0.11648 | 0.0065281 |
|  | Condition (reach vs hold) | 2.9168 | 0.11648 | 2.51E-54 |
|  | Laser:Condition (interaction) | -0.19812 | 0.16473 | 0.23108 |
| RFAil | Intercept | 0.92915 | 0.16365 | 1.73E-07 |
|  | Laser (hold) | -0.071143 | 0.12195 | 0.56112 |
|  | Laser (reach) | -0.29834 | 0.12195 | 0.016415 |
|  | Condition (reach vs hold) | 2.9244 | 0.12195 | 2.32E-40 |
|  | Laser:Condition (interaction) | -0.22719 | 0.17246 | 0.19113 |

We leveraged our large dataset to obtain highly resolved distributions of kinematic parameters under cortex-intact and inactivated conditions (4,473,441 primitives (19.3% inactivated) from 131,092 trajectories (21.7% inactivated) from 7 mice). We first wondered if cortex might contribute to primitive complexity. Importantly, making primitive-level comparisons between cortex-intact and inactivated trials requires that segments retain their shape and complexity regardless of condition. For example segments could be either simple point-to-point reaches or more complex, tortuous curves (Figure 8A-B). Cortical inactivations did not affect three independent measures of primitive complexity: segment tortuosity (p > 0.05 across all brain areas, LME, Table 13), the number of acceleration peaks per segment (p > 0.05 across all brain areas, LME, Table 14), or the probability that decomposed, minimum jerk submovements would overlap (p > 0.05 across all brain areas, Wilcoxon rank sum test on fraction of overlapping submovements, Table 15), (Figure 8C-H) (Novak et al., 2002; Rohrer et al., 2004). This lack of effect of cortical inactivations on primitive complexity was striking and further justified direct comparisons of primitive kinematics across conditions.

**Figure 8.**
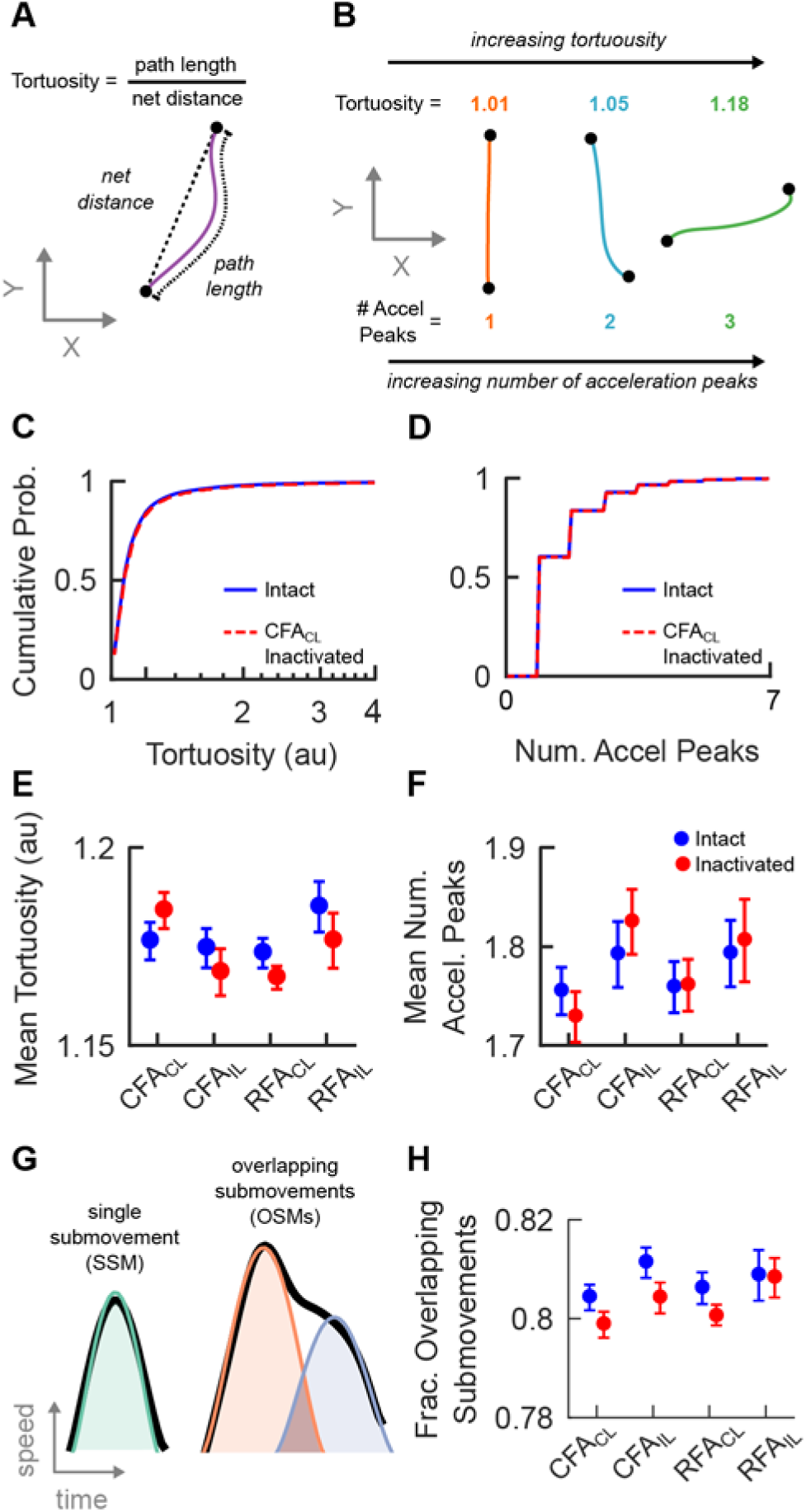
Motor cortical inactivations do not affect complexity of kinematic primitives. (A) Schematic defining the tortuosity of a segment. (B) Examples of segments with increasing complexity, as measured by increasing tortuosity (top) and number of acceleration peaks (bottom). (C) Cumulative probability distribution of the tortuosity values of segments in intact (blue) and CFAcl inactivated (red dashed line) trials. (D) Cumulative distribution of the number of acceleration peaks of segments in intact and CFAcl inactivated trials. (E) Mean±SEM of segment tortuosity with (red) and without (blue) inactivation of various motor cortical areas. (F) Mean±SEM of number of acceleration peaks in a segment with (red) and without (blue) inactivation of various motor cortical areas. (G) A complementary complexity analysis for submovement sequencing. Schematic showing a single submovement produced in isolation (left) and two submovements that overlapped (right). (H) Mean±SEM of the fraction of overlapping submovements with (red) and without (blue) inactivation of various motor cortical areas

**Table 12:** Segment direction, Watson U^2^ statistic

| Table 12: Segment direction, Watson U<sup>2</sup> statistic |  |  |  |
| --- | --- | --- | --- |
|  | **Reward Direction** | **Condition (reach v hold)** | **p value** |
| CFAcI | Lower Right | hold | 1.15E-04 |
|  | Lower Left | hold | 0.0057 |
|  | Lower Right | reach | 0.0608 |
|  | Lower Left | reach | 0.515 |
| CFaIl | Lower Right | hold | 0.4872 |
|  | Lower Left | hold | 0.8818 |
|  | Lower Right | reach | 0.4761 |
|  | Lower Left | reach | 0.3515 |
| RFaCl | Lower Right | hold | 0.1768 |
|  | Lower Left | hold | 0.3543 |
|  | Lower Right | reach | 0.0908 |
|  | Lower Left | reach | 0.2921 |
| RFaIl | Lower Right | hold | 0.2266 |
|  | Lower Left | hold | 0.2252 |
|  | Lower Right | reach | 0.7528 |
|  | Lower Left | reach | 0.6787 |

**Table 13:** Segment tortuosity, LME model

| Table 13: Segment tortuosity, LME model |  |  |  |  |
| --- | --- | --- | --- | --- |
| *Tortuosity ~ 1 + Laser + (1 \| Mouse)* |  |  |  |  |
|  | Model Term | Estimate | SE | p value of ttest |
| CFAcl | Intercept | 1.1766 | 0.0061316 | 1.04E-102 |
|  | Laser | 0.0081169 | 0.005917 | 0.17422 |
| CFAil | Intercept | 1.1753 | 0.007764 | 5.54E-66 |
|  | Laser | -0.0060958 | 0.0061618 | 0.32748 |
| RFAcl | Intercept | 1.1724 | 0.0050784 | 4.51E-105 |
|  | Laser | -0.0062216 | 0.0041455 | 0.13777 |
| RFAil | Intercept | 1.1856 | 0.0082281 | 1.71E-60 |
|  | Laser | -0.0085329 | 0.0082743 | 0.30806 |

**Table 14:**
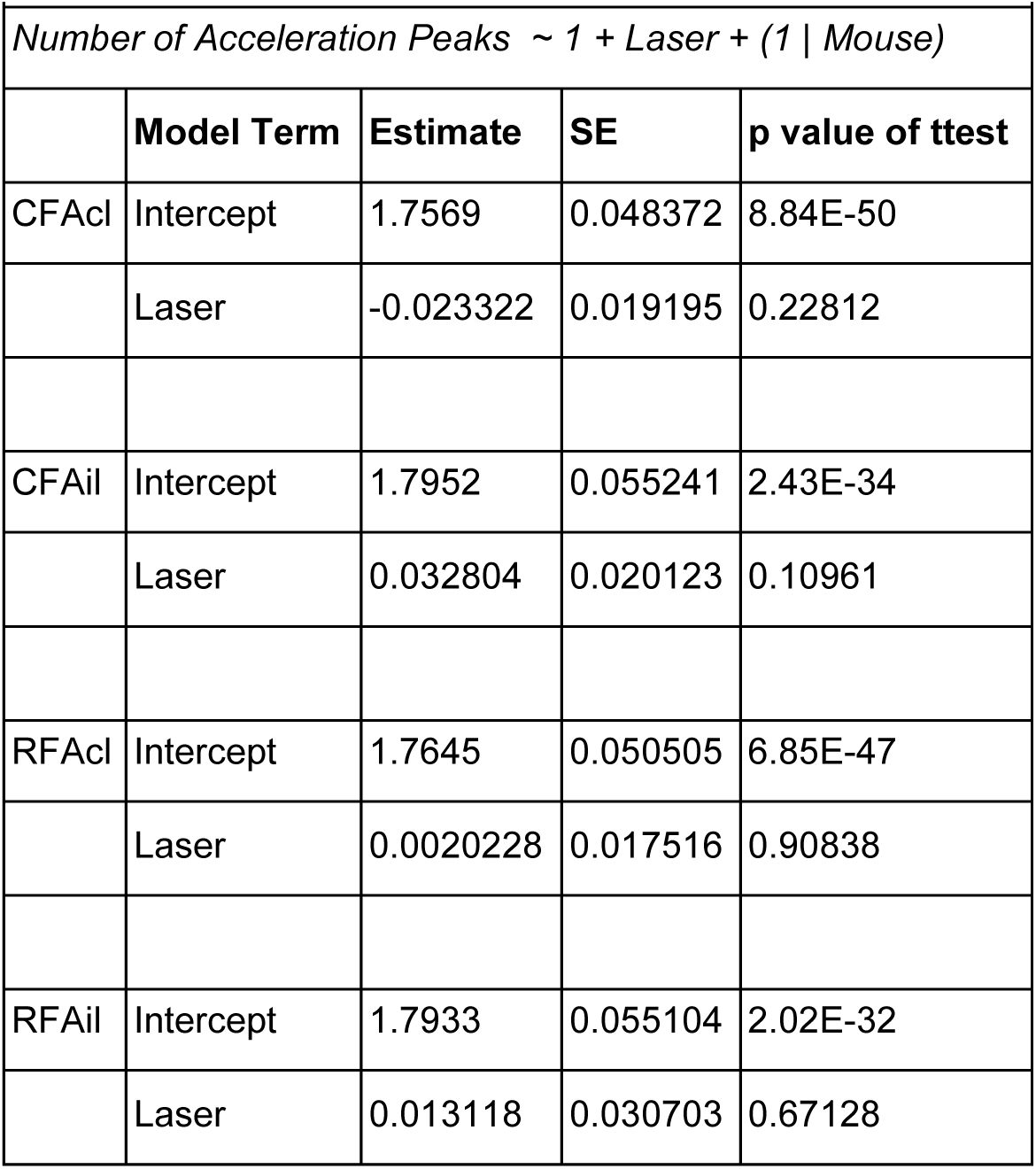
Segment number of acceleration peaks, LME model

**Table 15:**
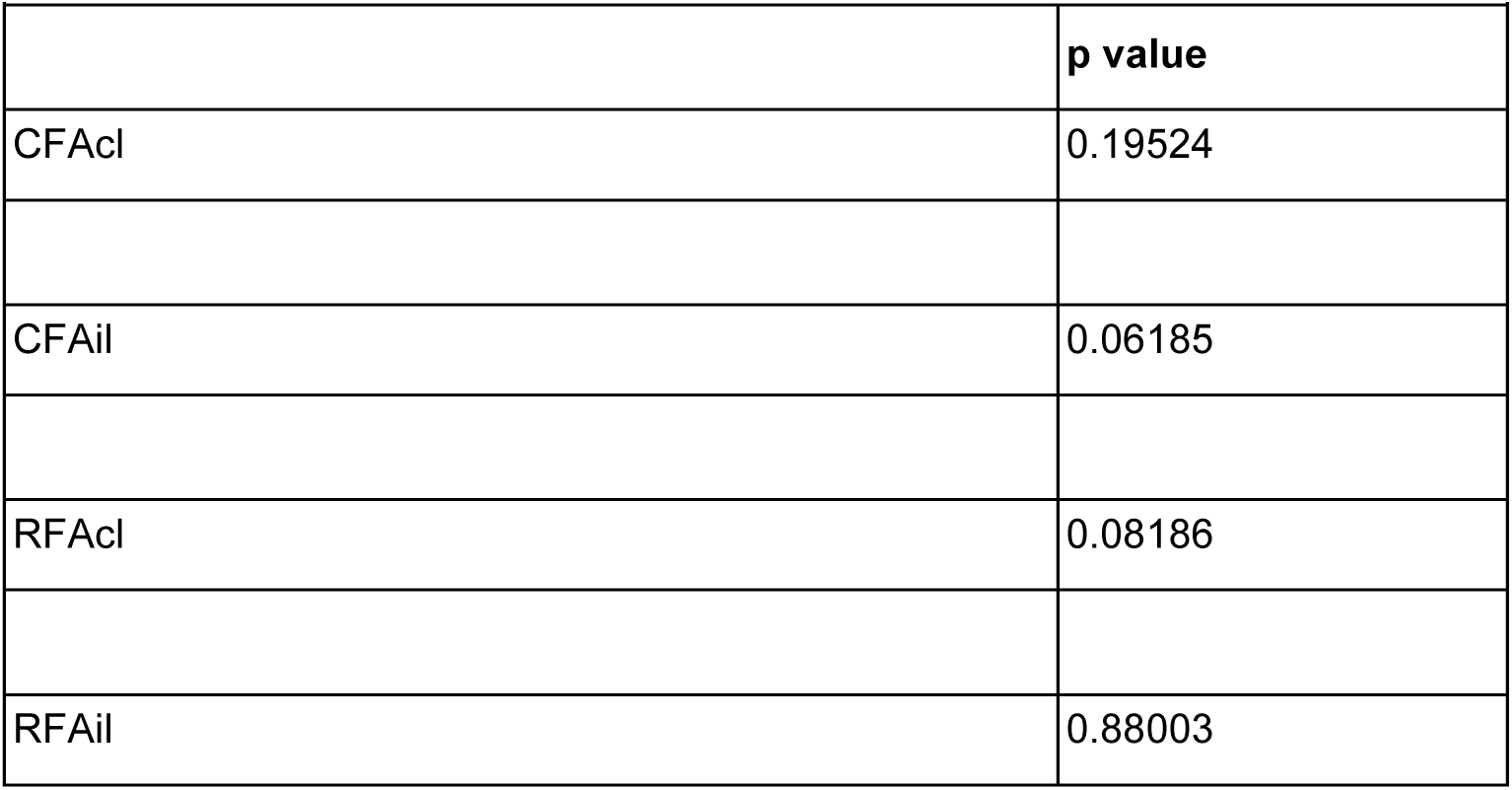
Fraction of overlapping submovements (OSMs), Wilcoxon rank sum test

|  |  |
| --- | --- |
|  | **p value** |
| CFAcl | 0.19524 |
| CFAil | 0.06185 |
| RFAcl | 0.08186 |
| RFAil | 0.88003 |

If cortical activity contributes to the neural command to generate a primitive, for example by redirecting the limb at a sharp turn, then silencing that area should alter primitive duration distributions (or equivalently, inter-segment onset intervals). Surprisingly, cortical inactivations did not change segment or submovement duration distributions for holds or for reaches (segment duration: *p* > 0.05 across all conditions, LME, Table 9; submovement duration: *p* > 0.05 across all conditions, Wilcoxon rank sum test, Table 15) (Figure 9D,G, Figure S4D,G), ruling against a primary role of motor cortex in generating or timing primitives in our task.

**Figure 9.**
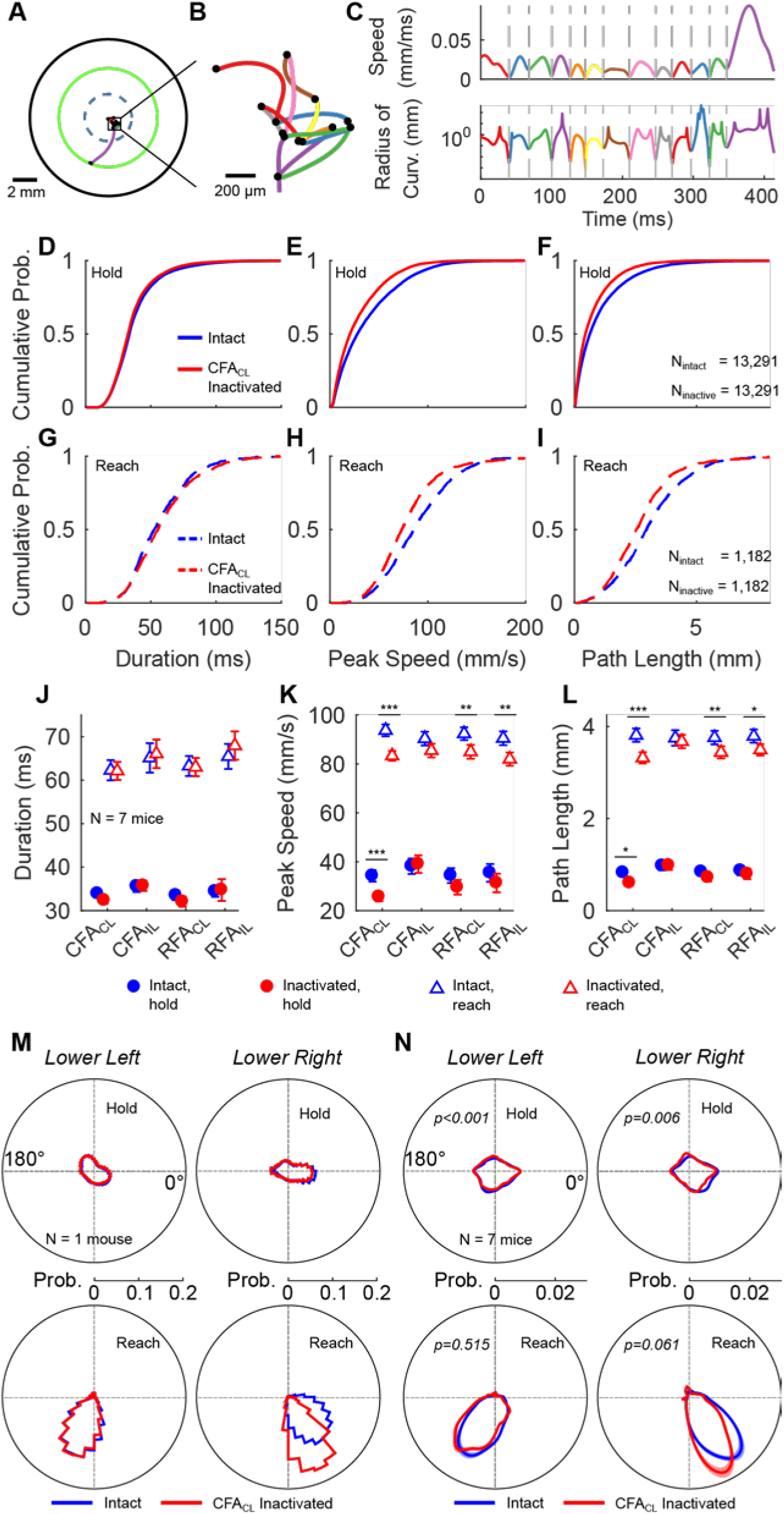
Effect of cortical inactivations on primitive kinematics. (A-C) Data plotted as in Figure 1A-B for a mouse forelimb trajectory. (A) Example mouse forelimb trajectory with black dots denoting boundaries separating decomposed segments. (B) Expanded view zooming in on micromovements produced during the hold period of the trajectory; different segments are color-coded. (C) Speed (top) and radius of curvature (bottom) are plotted as a function of time in the trajectory from A-B. Segment boundaries (gray dashed lines) are detectable as temporally coincident minima in speed and radius of curvature (see methods). (D-F) Cumulative probability distributions of durations (D), peak speeds (E) and pathlengths (F) of segments produced within the hold zone. Blue and red traces indicate data intact CFAcl inactivated trials, respectively. (J-I) Distributions plotted as in D-F for the single segment in a trajectory that transected the outer radius (i.e. the reach segment). (J-L) Mean±SEM of duration (J), peak speed (K) and pathlength (L) with (red) and without (blue) inactivation of various motor cortical areas for hold (circle) and reach (triangle) segments. (M) Direction distributions of hold (top) and reach (bottom) segments for an example mouse in intact (blue) and CFAcl inactivated (red) trials. (N) Data plotted as in M with average direction distributions across animals (n=7). See Figure S4 for similar analyses on the submovement class of kinematic primitives decomposed using a minimum jerk model.

CFAcl inactivation caused a significant reduction in segment speed and, because primitive durations were not affected, caused an associated reduction in segment pathlength (Figure 9D-L). This result was similarly observed for segments produced during both holds and reaches (peak speed during hold: 34.50 ± 3.76 mm/s (control) vs 25.89 ± 4.16 mm/s (CFAcl), *p* < 0.001; peak speed during reach: 94.04 ± 4.16 mm/s (control) vs 81.78 ± 5.20 mm/s (CFAcl), *p* < 0.001; path length during hold: 0.84 ± 0.15 mm (control) vs 0.62 ± 0.19 mm (CFAcl), *p* < 0.05; path length during reach: 3.83 ± 0.19 mm (control) vs 3.07 ± 0.27 mm (CFAcl) *p* < 0.001, LME, Tables 10–11). In contrast, inactivation of other motor cortical areas (RFAcl and RFAil) affected primitives executed during reaches but not holds (peak speed during reach: 93.02 ± 5.89 mm/s (control) vs 83.05 ± 7.29 mm/s (RFAcl), *p* < 0.01; path length during reach: 3.79 ± 0.23 mm (control) vs 3.27 ± 0.30 mm (RFAcl), *p* < 0.01; peak speed during reach: 93.05 ± 5.71 mm/s (control) vs 80.20 ± 7.47 mm/s (RFAil), *p* < 0.01; path length during reach: 3.85 ± 0.20 mm (control) vs 3.33 ± 0.29 mm (RFAil), *p* < 0.05; all other brain area/condition combinations for peak speed and path length: *p* > 0.05, LME, Tables 10–11).

**Figure 10.**
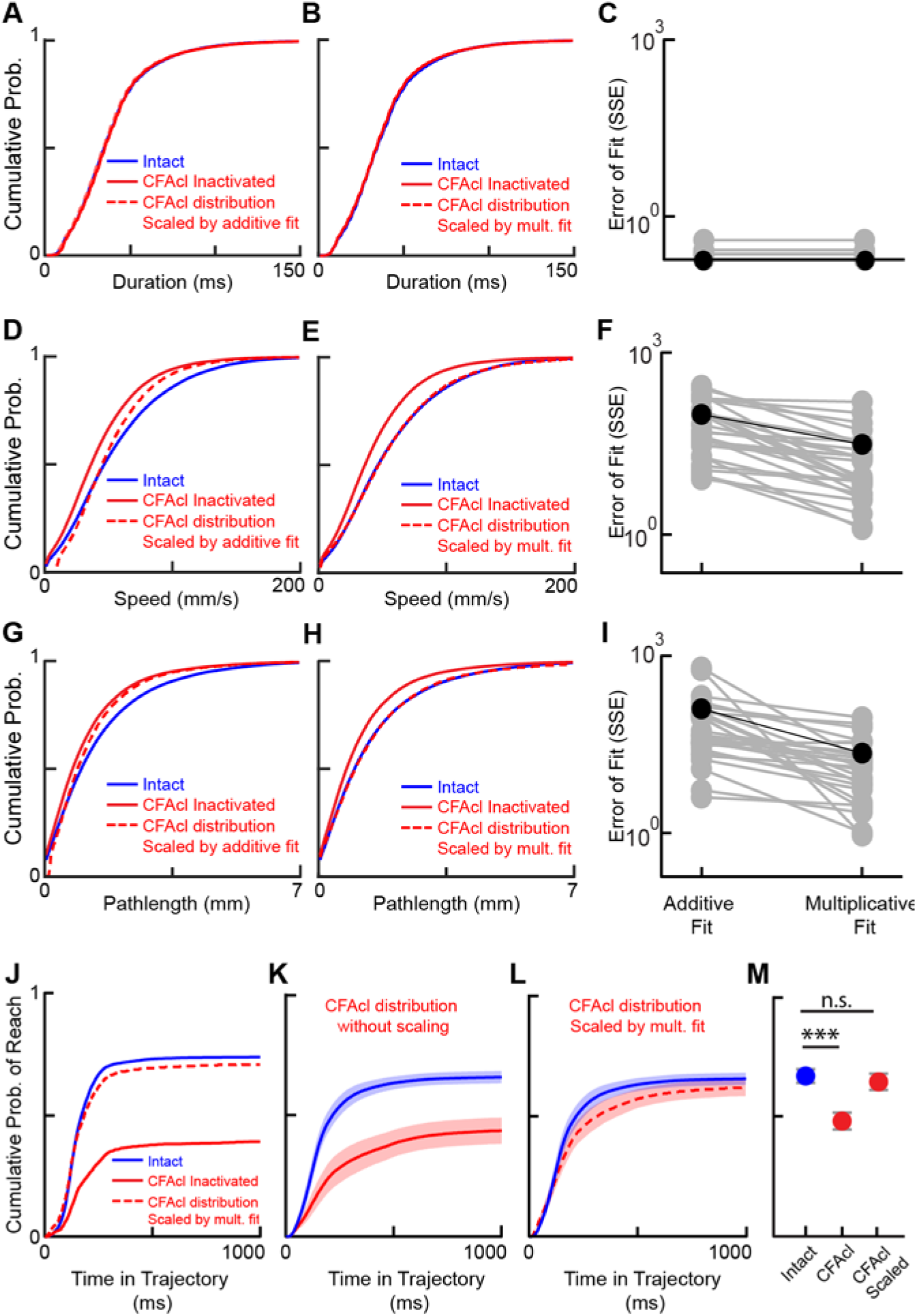
Primitive kinematic distributions from CFAcl inactivated dataset can be transformed into intact distributions by a single multiplicative gain factor. (A) Cumulative distribution of segment durations (intact, blue; CFAcl inactivated: solid red; CFAcl data rescaled by the best additive fit to match the intact distribution: dashed red). (B) Same as (A) but with CFAcl rescaled by the best multiplicative fit (see Methods). (C) Summed squared error for the additive and multiplicative fits for all animals. (n=7). (D-I) Data plotted as in A-C for peak segment speed (D-F) and pathlength (G-I) distributions. Note the extent to which the multiplicative (but not the additive) fits match the intact data, suggesting that CFAcl inactivation is associated with the loss of an amplitude-dependent (and not fixed) premotor drive. (J) Cumulative distribution from a single mouse of the probability of reaching outer radius as a function of time in intact trials (blue), CFAcl inactivated trials (solid red) and in trials with trajectories reconstituted by multiplying all primitives by the multiplicative factor determined from fitting pathlengths (dashed red). (K) Mean±SEM of probability of reaching outer threshold as a function of time in intact (blue) and CFAcl inactivated (trials) across mice. (L) Same as (K) but with CFAcl inactivated trials rescaled by a multiplicative fit determined from fitting pathlengths. (M) Mean ± SEM of probability that a trajectory would reach the outer radius, n=7 animals.

Finally, we analyzed how kinematic primitives were directed in space. Rather than assigning each trajectory a single direction value based on where it transected the outer radius (as in Figure 4), here each primitive in a trajectory was assigned a single direction value based on its peak velocity vector (see Methods). Inactivation of CFAcl had a statistically significant but extremely subtle effect on primitive direction distributions, in line with the absence of an effect at the trajectory level (hold, lower right reward zone: *p* = 1.15e-4; hold, lower left reward zone: *p* = 0.0057; reach, both reward zones: *p* > 0.05, Watson U^2^ test, Table 12) (Figure 9). Other cortical inactivations had no effect (p > 0.05 for all other brain areas/conditions, Watson U^2^ test, Table 12) (Figure S6).

All results on primitive duration, speed, pathlength and direction were independently replicated when we instead decomposed trajectories into submovements using a minimum jerk model (Figure S4 D-N, Tables 16–19).

**Table 16:** Submovement duration, Wilcoxon rank sum test

| Table 16: Submovement duration, Wilcoxon rank sum test |  |  |
| --- | --- | --- |
|  | Condition (reach v hold) | p value |
| CFAcl | hold | 0.49137 |
|  | reach | 0.31859 |
| CFAil | hold | 0.28021 |
|  | reach | 0.57837 |
| RFAcl | hold | 0.20225 |
|  | reach | 0.41309 |
| RFAil | hold | 0.29862 |
|  | reach | 0.42025 |

**Table 17:** Submovement velocity, Wilcoxon rank sum test

|  | **Condition (reach v hold)** | **p value** |
| --- | --- | --- |
| CFAcl | hold | 0.039819 |
|  | reach | 0.0016283 |
| CFAil | hold | 0.43024 |
|  | reach | 0.29142 |
| RFAcl | hold | 0.12417 |
|  | reach | 0.011623 |
| RFAil | hold | 0.15715 |
|  | reach | 0.045952 |

**Table 18:** Submovement path length, Wilcoxon rank sum test

|  | **Condition (reach v hold)** | **p value** |
| --- | --- | --- |
| CFAcl | hold | 0.039819 |
|  | reach | 0.00036394 |
| CFAil | hold | 0.43889 |
|  | reach | 0.18386 |
| RFAcl | hold | 0.098607 |
|  | reach | 0.019922 |
| RFAil | hold | 0.21775 |
|  | reach | 0.022086 |

**Table 19:** Submovement direction, Watson U2 statistic

|  | **Reward Direction** | **Condition (reach v hold)** | **p value** |
| --- | --- | --- | --- |
| CFACl | Lower Right | hold | 0.0536 |
|  | Lower Left | hold | 0.0728 |
|  | Lower Right | reach | 0.000987 |
|  | Lower Left | reach | 0.344 |
| CFAil | Lower Right | hold | 0.6753 |
|  | Lower Left | hold | 0.7345 |
|  | Lower Right | reach | 0.7981 |
|  | Lower Left | reach | 0.2914 |
| RFAcl | Lower Right | hold | 0.5678 |
|  | Lower Left | hold | 0.482 |
|  | Lower Right | reach | 0.1347 |
|  | Lower Left | reach | 0.453 |
| RFAil | Lower Right | hold | 0.4111 |
|  | Lower Left | hold | 0.1132 |
|  | Lower Right | reach | 0.6452 |
|  | Lower Left | reach | 0.4869 |

### Cortical inactivation reduced the gain of kinematic primitives

The observation that CFAcl inactivations similarly affected ~0.1 mm scale primitives executed during the hold and millimeter-scale ones executed during reaches suggests that the impact of CFAcl inactivations may follow a general rule. For example one possibility is that inactivated and intact kinematics differed by a constant value, which would be consistent with the loss of a fixed, amplitude-independent drive from the inactivated area. Alternatively, inactivated and intact kinematics could differ by a multiplicative value, which would be consistent with a decrease in gain, i.e. the loss of an amplitude-dependent drive from the inactivated area. To distinguish these possibilities, we combined hold and reach kinematic data into single distributions (Figure 10), and tested additive and multiplicative transformations of the inactivated data to “recover” cortex intact primitive distributions. We performed least-squares fits to determine the transformation that minimized the distance between the empirical distribution functions of the intact and the transformed-inactivated data. Intact peak speed and pathlength distributions were strikingly well fit by multiplying data from the inactivated dataset by a single ‘gain’ factor, while an additive model provided poor fits (sum of squared residuals (SSR) for duration: 0.058 + 1.07 (additive model) vs 0.058 + 1.07 (multiplicative model); peak speed: 388.1 ± 1.2 (additive model) vs 81.8 + 2.0 (multiplicative model); path length: 647.3 ± 2.0 (additive model) vs 64.5 +1.5 (multiplicative model)) (Figure 10 A-I). The high quality of the multiplicative fits suggests that cortical inactivation simply reduced the gain of primitives. Consistent with this, multiplying the pathlength of all segments produced under CFAcl inactivation by a single gain factor (see Methods), resulted in a probability of reaching out that was indistinguishable from intact trajectories (p(transecting outer radius) intact: 66.74±2.83% vs FAcl inactivated and rescaled: 64.31±3.4%, p>0.05, paired t test). Thus CFAcl inactivation-associated reduction in primitive peak speed and path length was sufficient to explain the undershooting observed at the trajectory level (Figure 10 J-M).

## DISCUSSION

We developed a touch-sensing, low torque joystick that resolves mouse forelimb kinematics with micron-millisecond spatiotemporal precision. We built joysticks into a computerized homecage system that automatically trains mice to produce complex, directed center-out forelimb trajectories while implementing closed-loop optogenetics. We then tested several hypotheses about how different motor cortical areas contribute to maintaining limb position and reaching to targets. We also used trajectory decomposition to test hypothesis about how kinematic parameters of primitives are controlled. We find that inactivation of contralateral CFA reduces the peak speed of kinematic primitives, but preserves their timing, complexity and direction. As a result, trajectories exhibited isomorphic hypometria, i.e. they retained their shapes and remained appropriately directed to targets but were spatially contracted. Our results identify conditions where motor cortical inactivation simply reduces the gain of motor output.

### Forelimb motor control in rodents

While it is clear that rodent motor cortex is required for learning (Kawai et al., 2015; Peters et al., 2017), the roles of cortex in executing movements is complicated. First, permanent lesion, pharmacological inactivation, and photoinhibition can yield dissimilar results even during seemingly similar tasks (Otchy et al., 2015). Cortex lesions preserve ethologically relevant, potentially innate behaviors such as grooming, eating, swimming, fighting, playing and walking over obstacles, as well as producing previously learned behaviors such as grasping for pellets and producing learned lever tapping patterns (Grill and Norgren, 1978; Kawai et al., 2015; Sorenson and Ellison, 1970; Whishaw and Kolb, 1983; Whishaw et al., 1981). Yet acute inactivations impair prehension and cue-guided action selection under head-restraint (Galinanes et al., 2018; Guo et al., 2015; Miri et al., 2017; Morandell and Huber, 2017; Peters et al., 2014). Distinct task requirements in different behavioral paradigms result in different dependence on motor cortex, but core principles that dictate when a behavior will or will not require cortex remain frustratingly elusive.

To reduce task complexities and begin to dissect minimum circuits for the production and sequencing of primitives, we developed a task uncomplicated by head restraint, cues, visual guidance, or externally-induced sensory prediction errors yet rich enough to produce complex trajectories that are directed to spatial targets. Under these conditions loss of motor cortex caused a remarkably clear and specific deficit: reduced gain of movements of all amplitudes.

First, mice gradually learned to hold their right forelimb still in a small 2 mm ‘hold zone.’ Static maintenance of limb posture during appetitive tasks may require specific patterns of activity to actively suppress movement and/or actively maintain current limb position (Ebbesen and Brecht, 2017; Kaufman et al., 2014; Shadmehr, 2017). Given the joystick’s compliance, holding still while applying downward force (i.e. during gripping) created an inverted pendulum, or stick balancing, problem, in which corrective micromovements are required to maintain position (Anderson, 1989; Bhounsule et al., 2015; Cabrera and Milton, 2002). Stabilogram diffusion analysis suggested that mice used an active control policy to maintain center-position that, interestingly, strongly resembled one used for maintenance of upright posture in humans (Peterka, 2002).

The generation, direction and timing of submillimeter-scale primitives produced during the hold were surprisingly intact during cortical inactivations, suggesting that subcortical pathways are sufficient to produce corrective movements required for postural control during holding-still (Azim et al., 2014; Murray et al., 2018; Walter et al., 2006). However, the change in the offset, slope, and inflection point of the SDF observed during motor cortical inactivation suggests that, although the cortex is not required for the performance of the hold, it could modulate corrective processes underlying postural stability.

Aftger executing the hold, mice reached out in different directions to spatial targets. CFAcl inactivation reduced both peak trajectory speed and impaired the ability to reach out. Previously, CFAcl photoinhibition resulted in halted or impaired prehension (Galinanes et al., 2018; Guo et al., 2015; Wang et al., 2017), and failure to push or pull a joystick in response to a cue (Miri et al., 2017; Morandell and Huber, 2017; Peters et al., 2014). Our novel findings emerged directly from our ability to resolve the tiny details of motion that persisted following CFAcl inactivation. First, at the trajectory level, neither reach direction nor variability were affected by cortical inactivation. Second, at the primitive level, direction, tortuosity, acceleration patterns and probability of submovement overlap were also not significantly affected. Even primitive duration distributions, which provide proxies for the rate at which the CNS sends efferent movement initiation commands to the periphery, were not affected. These data provide convergent evidence for subcortical control of primitive generation and patterning in our task. Notably, a learned sequential tapping task in rats also implicated subcortical circuits in these functions (Kawai et al., 2015).

### Motor cortex inactivation decreased gain of motor primitives

Behavioral deficits following photoinhibition do not necessarily reveal the function of the inactivated area (Otchy et al., 2015; Wolff and Ölveczky, 2018), cautioning against interpreting ‘positive’ results in our experiments as revealing the function motor cortex. Specifically, we regard it as unlikely that the sole function of motor cortex, with its complex recurrent circuitry and detailed kinematic representations (Brecht et al., 2013), is to simply to control the gain of movement. Our main discovery is not the positive result on speed in isolation; it is moreso the negative results showing that the cortical inactivation has minimum effect on primitive initiation, direction and complexity and, relatedly, on trajectory shape.

Another caveat is that mice are clearly not primates, and our task differed in several respects from canonical primate center-out reach tasks. First, whereas our task gradually shaped reach direction over day timescales, most primate reach tasks are visually guided and explicitly cue target locations on a trial-by-trial basis (Georgopoulos et al., 1988). Second, our animals were freely moving and only engaged the task of their own volition in their homecage, whereas studies in primates typically involve water/food deprivation and head restraint. Third, primates have a highly specialized forelimb motor cortex that includes multiple corticospinal regions and direct projections to motor neurons (Dum and Strick, 2002; Lemon, 2008). Finally, whereas cortex-lesioned rodents exhibit remarkably intact behavioral repertoire (Sorenson and Ellison, 1970; Whishaw and Kolb, 1983; Whishaw et al., 1981), large cortex lesions in primates are associated with more severe effects including paresis that can be refractory to recovery, especially without subsequent training (Darling et al., 2011; Nudo, 2013; Zeiler and Krakauer, 2013). All of these considerations likely underlie important differences in cortical control of movement across species.

Caveats aside, our main result - that cortical inactivations reduce the gain of primitives, resulting in hypometric but otherwise intact trajectories - bears an uncanny resemblance to several studies in humans and non-human primates, even replicating some very specific details. Cortical stroke, GPi inactivation, Parkinsonism and even some types of cerebellar damage can result in weakness and bradykinesia but with essentially intact initiation and direction (Darling et al., 2011; Lawrence and Kuypers, 1968; Manto et al., 1998; Passingham et al., 1983). Motor cortex lesion can result ‘undershooting’ of appropriately-timed and directed wrist movements (Hoffman and Strick, 1995), or mastication patterns with normal timing and patterning yet with contracted trajectories (Larson et al., 1980). Motor cortex lesion can also diminish the amplitude of essential tremor, showing a specific case where loss of cortex decreases the gain of subcortically generated (albeit pathologic) movements (Dupuis et al., 2010; Kim et al., 2006). Inactivation of basal ganglia output and Parkinsonism also result in bradykinetic, hypometric reaches that are otherwise appropriately timed and aimed (Desmurget and Turner, 2008; Panigrahi et al., 2015), again suggesting an oddly similar phenotype. The essence of these deficits are evident in analysis of handwriting in patients with Parkinson’s: micrographia, or shrunk text, is due to normally timed and directed strokes that simply have reduced peak speed and, therefore, pathlength. Notably, the magnitude of bradykinesia and a hypometria is usually proportional to movement amplitude suggesting the loss of an amplitude-dependent drive (Broderick et al., 2009; Van Gemmert et al., 2003), exactly as we observed (Figure S8). Finally, hypometric reaching can also be observed following inactivation of cerebellar outputs (Cooper et al., 2000; Martin et al., 2000). Thus our results capture essential features of conventionally distinct pathologies: motor cortex lesion, inactivation of basal ganglia or cerebellar output, and Parkinsonism.

Why are bradykinesia and hypometria such common failure modes of the motor system? The answer may not lie in the idea that disparate parts of the motor system are ‘dedicated’ to speed or gain control (Golub et al., 2014), and may instead in control theoretic accounts of movement (Scott, 2004; Shadmehr and Krakauer, 2008; Todorov and Jordan, 2002). The essence of this framework is that the motor system can be described as one or more feedback controllers generating movement commands while minimizing effort and maximizing target accuracy. Consistent with this idea, disrupting feedback during a goal directed reach causes oscillations around the target (Azim et al., 2014; Flament et al., 1984), similar to what occurs when feedback is disrupted in canonical feedback systems, such as PID controllers used in servo motors (Aström and Murray, 2010). However, instead of inactivating the feedback, what would happen if you inactivate the *output* of the control loops? In cases where the system is described by a single control loop, loss of output means, trivially, loss of behavior. However, in a system with multiple control loops operating in parallel, the specific behavioral deficit associated with loss of a single loop can be complicated because each controller’s contribution to behavior is task-dependent (Wolpert and Kawato, 1998). Each controller is likely to also play unique roles in learning (Doya, 1999).

But for well-learned behaviors detailed kinematics of upcoming movements are represented in cortex, basal ganglia and cerebellum (Fortier et al., 1989; Fu et al., 1997; Schwartz, 2007; Scott, 2003; Shenoy et al., 2013; Turner and Anderson, 1997; Wong et al., 2014). These representations suggest that when the system is well trained multiple controllers have acquired useful models that contribute to motor output, such as initiating a primitive and directing its path to a target. It follows that transiently losing a single controller could reduce the drive of the system towards the target, but the behavior would remain essentially intact due to the ongoing commands of the other, non-inactivated controllers. This hypothesis attempts to account for the isomorphic hypometria observed in our study (in which mice were highly trained) as well as the observation that diverse insults to the adult human motor system result in conceptually similar micrographia (Inzelberg et al., 2016). An interesting prediction of this hypothesis is that, in our well-trained center-out task, inactivating basal ganglia output, cerebellar output, or inducing Parkinsonism could all lead to similar deficits, an intriguing prediction supported by previous work (Cooper et al., 2000; Desmurget and Turner, 2008; Martin et al., 2000; Panigrahi et al., 2015) that is now easily testable with our open-sourced automated system.

## ACKNOWLEDGMENTS

We thank Joe Fetcho, Melissa Warden, Lena Ting, and Goldberg lab members for helpful discussions and providing comments on the manuscript. JHG was supported by NIH (DP2HD087952), the Pew Charitable Trusts, and the Esther A. & and Joseph Klingenstein Fund. T.B. and S.C.W. were supported by the Cornell Mong Neurotech Fellowship.

## AUTHOR CONTRIBUTIONS

T.B. and J.H.G designed and conceived the project. T.B performed the experiments. T.B and S.C.W analyzed and interpreted the data with input from J.H.G and I.C. T.B. designed and constructed the hardware and software components with input and assistance from J.S.W, N.P, R.S., N.S. and B.K. J.H.G, T.B., S.C.W and B.K wrote the manuscript.

## DECLARATION OF INTERESTS

The authors declare no competing interests

## METHODS

### CONTACT FOR REAGENT AND RESOURCE SHARING

Further information and requests for reagents should be directed to, and will be fulfilled by the Lead Contact Jesse H. Goldberg.

### EXPERIMENTAL MODEL AND SUBJECT DETAILS

All experiments and procedures were performed according to NIH guidelines and approved by the Institutional Animal Care and Use Committee of Cornell University.

#### Experimental Animals

A total of 11 adult male VGAT-ChR2-EYFP line 8 mice (B6.Cg-Tg(Slc32a1-COP4*H134R/EYFP) 8Gfng/J; Jackson Laboratories stock #014548) were used. The mice were between 18-40 weeks old for the duration of the experiment. All animals were individually housed under a 12hr light/dark cycle for the duration of the study and had continuous access to the task. The animal’s daily water intake was monitored using an automated system. If the water intake was less than 1ml, the system automatically dispensed water to make up the difference.

### METHOD DETAILS

#### Surgery

Under isoflurane induced anaesthesia, we implanted 7 of the mice with 400um 0.43 NA fiber optic cannulas bilaterally over Caudal Forelimb Area (CFA) (0.5mm A/P, ±1.5mm M/L) and Rostral Forelimb Area (RFA) (2.2mm A/P, ±1.0mm M/L).

#### Principle of Operation and Signal Processing Pipeline

##### The joystick

*The need for a custom joystick to monitor forelimb kinematics in mice*. We considered commercially available joysticks such as: HF46S10 from APEM, the 67A from grayhill and the inductive joysticks from CTI. There were five main shortcomings of these products that made them unsuitable for the study of kinematics in mice. (1) Isometric force profile: All the industrial joysticks have a recentering mechanism that relies on two compression springs set along orthogonal directions. This always leads to moving along the diagonal being up to 40% harder than along the axes of the springs. We solved that problem by using one set of centering magnets right below the joystick. (2) Low displacement force: The industrial joysticks have a displacement force of about 100-250 gf (gram force), ~2.5 Newtons of force, this can be reduced by having a long lever arm. For example, you can lengthen the lever arm by 10 times to get into a 0.1-0.2N range, however, this still puts a lot of load on mouse forelimb kinematics. The centering mechanism we use gives us 0.0081 N/mm, or about 32 mN to move the full range of the joystick, this allows us to get get the fine details of motion even when the animal is holding still (10-100 um scale) and finer resolution when the mouse is making large (~ 1 mm scale) movements. (3) Future Force Controllability: Our design supports the dynamical control of displacement force on the joystick. By switching out the centering magnet with an electromagnet, we can dynamically control force in real time, and coupled with our real-time microsecond FPGA, we can make that force dependent on a behavioral variable like speed. (4) Built-in touch sensor: Off the shelf joysticks do not provide a readout of when the animal is actually holding the joystick, complicating attribution of displacement by potentially confusing active (animal-induced) versus passive motion. (5) Dead Space: Some commercial joysticks are intentionally designed (in hardware) to have dead-space. i.e. there will be no reading if the joystick is moved in within a certain distance. Presumably to protect against accidental movement of the joystick.

*A custom joystick that solves these problems*. The core of our joystick system was a two Axis Hall Sensor (Sentron, 2SA-10G). Moving the joystick changed the angle of the incident magnetic field generated by the magnets on the hall sensor. The change in incident magnetic field provided X and Y voltage (0-5V) as a linear readout of displacement. The top of the joystick was a conductive ball which was connected to an active capacitance touch sensor (AT42QT1011) that provided instantaneous read on the joystick being contacted. To calibrate the joystick, i.e. to determine the voltage vs distance relationship, we set up the joystick in a custom, precision machined setup that allowed the manipulandum to move along a narrow channel along an outward directed radial axis. Then, using an electronic screw gauge, the joystick was displaced from the center position along the radial axis in increments of 500um. At each displacement we recorded the voltage from the joystick’s hall sensor. We then repeated these measurements along a radial axis in a direction non-orthogonal to the first. To quantify spatial resolution we first measured the x, y voltages from the device when it was at rest, used the calibration to convert the signal from volts into units of distance (mm). We then calculated the standard deviation of the distance distribution over 2000 samples (2sec) for each device. To quantify stiffness of the joystick, we rotated the joystick by 90 degrees, such that the joystick manipulandum was horizontal to the ground, and applied varying weights to the end of the joystick while measuring the displaced distance. We repeated these measurements by rotating the joystick around the axis of the manipulandum in ~30 degree increments. See supplemental methods for construction and design details.

##### Fixed Post

A “Fixed Post” was placed in the home-cage and it also had a conductive ball connected to an active capacitance touch sensor (AT42QT1011).

##### Nose Poke

The Nosepoke Sensor was a modified IR diode-phototransistor pair (LTH-301-32) from Lite-on devices. The IR LED and the phototransistor were separated and placed across the nose poke port. When the mouse put its snout through the port, it broke the IR beam and drove the signal high.

##### Water Delivery

The mice received water through a lick spout (H24-01-TB-01, Coulbourn Instruments) connected to a precision solenoid valve (Lee Company LHDA2433215H). The valve received water from a reservoir whose level was constantly maintained by closed loop water recirculation, providing stable microliter precision in water delivery over week timescales.

##### Signal Conditioning Circuitry

We built signal conditioning circuitry at each homecage to calibrate the sensors and provide noise immunity. Specifically, we (1) used unity gain buffer amplifiers (LM358, Texas Instruments) to protect the joystick signals from noise as they were routed to the computational system; and (2) calibrated the nose poke sensor by biasing the phototransistor at the edge of the linear zone to reduce detection hysteresis, and used a comparator (TL331, Texas Instruments) to convert it to a TTL digital output.

##### Breakout Board and Voltage Protection System

The digital and analog sensor signals from all the behavioral boxes reached the computational unit (CU) through the breakout board. On the breakout board, for each line we used a Thermistor (PRF18BB471QB5RB) to protect the CU from high current surges and Schottky diodes (TBAT54S, Toshiba Electronics) to protect the CU from high voltages. After the voltage/current protection circuitry, analog and digital lines from each behavior box were separated and routed to the right connector.

#### Computational System

##### Hardware

The core of the computational system was a Single Board Real-time Input/Output system (sbRIO-9636, National Instruments). The sbRIO had both a Field Programmable Gate Array (FPGA) and a Real-Time Processor (RTP) on the same board. It had 16 Analog inputs, 4 analog outputs and 28 Digital I/O ports. The FPGA is programmed through LabVIEW. See supplemental methods for construction and design details.

##### Overview of the Software Infrastructure

There were four major components of the software architecture: (i) Every millisecond the FPGA code processed both the digital inputs (nose poke sensors, touch sensors) and analog inputs (Joystick X,Y) to determine if the trial was “live” and if the “hold” and reach-out contingency was met. The FPGA code also directly interacts with the digital outputs. It controlled the solenoid valve for the water reward, the masking light used during photoinhibition and patterns the optogenetic output. The FPGA took ~37 microseconds to process the contingency requirements. At the end of each millisecond, the FPGA wrote all the input sensor data and the generated outputs to a FIFO register on the FPGA. Using an onboard FIFO prevented sampling errors and discontinuities. (ii) Every second, the code on the RT processor read from the FIFO register using a Direct Memory Access method (DMA) and packaged it into a single array element. The sbRIO had limited onboard memory and could not hold the vast behavioral data being generated, it required a secondary storage solution. Importantly, the data-transfer needed to be fast, in real time and have temporal continuity. We used Network Stream Objects to set up communications between the sbRIO and the server grade acquisition computer. We developed a simple data protocol containing the data, the timestamp and a frame ID. The data from the RT wass packaged using this protocol and sent over the Network to the acquisition computer. In addition to packaging and transferring the data, the code on the RT also had the programmatic access to all the parameters of the task, i.e. hold time, reach angle, water dispense time, optogenetic power, optogenetic stimulation time and had the ability to deliver rewards to mouse boxes with a tunable poisson distribution. (iii) The code on acquisition computer (“PC code”) was set up to interface with the appropriate sbRIO and continuously monitored the network stream for any new data packets. If a data packet was detected, the PC code unpacked the data, split it into separate channels, identified the sequence of the data, created filenames for each data file, and then wrote the behavioral data onto the RAID server with generated filenames. Each filename was coded to provide information on the mouse it came from and the settings of the behavioral paradigm. (iv) During training, the behavioral data was automatically analyzed at the end of the day (~11 PM) to determine the new training parameters. The code combined and extracted all valid trials joystick trials from the day, the number of rewards, distributions of hold-times and reach directions. If the animal hadn’t had enough successful trials to meet its daily water requirements, the system communicated with the RT code to automatically dispense water to make up the difference and the contingency of the task remained unchanged. If the animal met its daily requirement of water intake, the contingency wass updated to reward only 15% of the trials based on the current distributions of the hold time and reach direction

##### Optogenetics light source and control

We used Laser LED light sources (LDFLS_450-450, Doric Life Sciences Ltd), attached to an optical rotary joint (FRJ_1×2i_FC-2FC_0.22) and delivered light to the implanted cannulas using 400um, 0.43NA lightly armored metal jacket patch cords. The light sources were set to analog input mode and driven with a sinusoidal pulse (40 Hz, 15mW peak) generated by the sbRIO.

#### Behavioral Shaping and Photoinhibition

We used a sequence of reward contingencies to train mice to perform ‘hold-still’ + direction-specific center-out reaches. i) On day 1 water restricted mice (~6hrs) were placed into joystick endowed home cages. Water wass automatically dispensed with a poisson distribution (mean 150s). If the mice simultaneously did a nosepoke (np) and contacted both joystick and the fixed post (fp) for more than 50ms they were rewarded with water. ii) On Day2-Day3 after mice started interacting with the joystick, we changed the reward contingency such that the mice only get rewarded if they contact the Joystick after both the NosePoke and FP. (iii) After the mice learned to contact the joystick last in sequence, we changed the reward contingency such that mice had to move the joystick out to at least 4mm from the center in order to get rewarded. (iv) Once the mice started reaching out to 4mm, we gradually shaped the “hold-time” of the trajectories. Hold time was defined at the amount of time the trajectory spent within 2mm radius of the center. We shaped the trajectories by setting the required hold-time for reward each day as greater than the 75th percentile of hold time distribution of the previous day. This was repeated till the 75th percentile of the hold time was greater than 100ms. At which point, the required hold-time was set to 100ms. (v) We then characterized the reach-angle distributions of the mice. The reach-angle was defined as the angle at which the trajectories crossed the 4mm reward threshold. (vi) After characterizing the reach angle, we performed the first set of CFA and RFA inactivations, we iterated through CFAcl, CFAil, RFAcl, and RFAil by moving the fiber optic cable onto the relevant implanted cannula and turning on the laser in ~15% randomly interleaved trials (vii) The reach-angle was gradually shaped clockwise (CW) or counter clockwise (CCW) by rewarding either the 15th percentile (CW) or the 85th percentile (CCW) of the reward distribution till the mean of the new distribution was different by at least ~60 degrees. (viii) We then stopped the angle shaping and performed the second set of CFA and RFA inactivations.

### QUANTIFICATION AND STATISTICAL ANALYSIS

#### Trajectory level analysis

Trajectories were acquired at 1kHz and low pass filtered at 50Hz with an 8-pole Butterworth filter in software. Hold-time was defined as the amount of time the trajectory was within the inner radius (2mm) starting from joystick contact. The reach direction was defined as the angle at which the outer radius was transected by the trajectory. Velocity was calculated as a one sample difference of the position vector. To make sure that we were specifically analyzing trajectories that were attributable to right forelimb movement with the animal in a consistent posture, only the trajectories that were contacted after the nose poke and fixed-post contact were considered valid trials eligible for reward. If the mouse exited the nose poke, lost contact with the fixed post or the joystick, the trial was immediately failed. The mouse also failed the trial if it exited the inner-radius earlier than its hold-still requirement or if it reached in the wrong direction. There were no cues for failed or successful rewards (except the water delivery apparatus). However, a blue LED masking light roughly at eye level to the mouse was turned on at joystick contact. The mouse had to re-contact the joystick in order to start a new trial after both successful and failed trials.

#### Minimum jerk decomposition and trajectory segmentation

In order to decompose complex trajectories into primitives, we employed two methods: the first was to fit trajectory kinematics to a linear combination of minimum jerk basis functions (called submovements) (Gowda et al., 2015; Rohrer and Hogan, 2003; Viviani and Flash, 1995), and the second was to break up trajectory kinematics into segments, where segment boundaries were defined by temporally coincident minima of velocity and radius of curvature (Milner, 1992; Viviani and Terzuolo, 1982). To implement the former, we followed the method from (Gowda et al., 2015), using the MATLAB package described therein. In short, the code performed a least-squares fit of the six-dimensional description of joystick kinematics (position, velocity, and acceleration for both the x and y coordinates) to a series of minimum-jerk basis, whose velocity profile took the functional form:

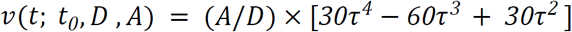

Where *τ* gives the normalized time, defined as *τ = (t − t*_0_*)/D, t*_0_ is the initiation time of the submovement, *D* is the duration of the submovement, and *A* is the amplitude. Importantly, the function is defined to be zero outside the bounds *[t*_0_*, t*_0_*+ D]*, The fit procedure did not require strong assumptions about the number of submovements prior to running the fit, but rather iteratively added submovements to improve the overall fit until a cost function threshold was reached.

The second decomposition algorithm, which used coincident velocity and radius of curvature minima and is the primary decomposition method discussed in the main text, did not rely on any optimization or assumptions regarding the parameterization of primitives. To determine these segment boundaries, we identified the velocity and radius of curvature minima for each trajectory, and a assigned a segment boundary whenever these minima were temporally separated by no more than 1 ms. This method acted as a sharp turn detector, and identified temporal boundaries within a trajectory that must have arisen from a new force applied to/via the forelimb.

#### Segment Analysis

We performed analyses on primitives, obtained using the methods described in the previous section, in order to gain a finer-scale description of joystick trajectories. Because we are primarily interested in task-related movements, we excluded from our analyses the first primitive produced during each trial. This ensured that our data set did not include motion related to grasping the joystick, but instead only included controlled motion. Furthermore, we excluded from our analyses any primitive with duration <10 ms. From the primitives we obtained through decomposition/segmentation, we extracted various kinematic parameters to describe said primitive. In the case of minimum jerk basis functions, each submovement was uniquely defined by four parameters: the start time, duration, amplitude (path length), and direction of the submovement. Each of these was calculated directly from the code in (Gowda et al., 2015). Peak speed for submovements was obtained by taking the ratio of amplitude to duration, multiplied by the constant 1.875 (reflecting the characteristic bell-shape of the submovement basis function). Finally, because multiple, distinct submovements can be occuring at the same time within a trajectory, we measured the fraction of submovements that have temporal overlap (calculated as number of overlapping submovements divided by total number of submovements) as a proxy for motion complexity--when multiple submovements overlap, they give rise to velocity profiles that have richer structure than a train of non-overlapping bell-shaped velocity peaks. In the case of segments defined by coincident minima of velocity and radius of curvature, there was no set number of descriptors that uniquely defines a segment. Segments varied from simple point-to-point reaches to motions with a more complex spatial and velocity profiles. To quantify the degree of complexity for each segment, we used two complementary metrics: the tortuosity of the segment (defined as the path length divided by the distance between start and end points) and the number of acceleration peaks within a trajectory (found using standard peak identification techniques in MATLAB). We also extracted from segments various other kinematic parameters, including the duration, path length, peak speed, and direction. Here we defined segment direction as the angle between the initial position of the segment and the position of the segment at its maximum speed. Finally, because we were interested in the differential effects of cortical inactivations on different types of movement, for many analyses we separately analyzed the primitives involved in the “hold still” and “reach” portions of the task. The “hold” primitives for each trial were defined as the primitives that occur prior to the first inner threshold crossing of the trajectory--this corresponded directly to the definition of the hold period in the task structure. The “reach” primitive for each trajectory was defined as the first primitive to cross the outer threshold, regardless of whether the hold period was completed successfully. Thus each trajectory contained at most one reach segment, but potentially multiple overlapping reach submovements. We used these categories to analyze the movement-type-dependent effects of cortical inactivations, as in Figure 7.

#### Stabilogram Diffusion Analysis of the ‘Hold-Still’

The stabilogram diffusion function (SDF) gives the mean squared displacement over a specified time window. The SDF was calculated on ‘hold-still’ part of the trajectories by only analyzing the trajectory up to the onset of the segment that exited the inner radius. The mean squared displacement <Δr^2^> for a time interval Δt is given by:

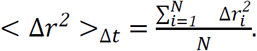

Where N is the total number of points for the time interval across all the trajectories for that condition, and Δr^2^ is the squared displacement for each of those of points. The slope and the offset of the log-log plot of the SDF is estimated by linear fit to the first 10ms of the SDF. The inflection point is defined as the intersection of the line estimated by the linear fit of the first 10ms with the tangent drawn at the peak of the SDF in the first 100ms.

#### Statistical Analyses

To determine the effect of cortical inactivation on segment and trajectory kinematic distributions, we fit linear mixed-effects (LME) models to our data set, with random intercept terms to account for variation both i) between mice and ii) across days for a given mouse. We added fixed effects corresponding to i) whether the segment or trajectory was performed during a trial with optogenetic manipulation, ii) in case of segments, whether the mouse was executing the ‘hold’ or ‘reach’ portion of the task (called ‘Condition’), and iii) the interaction between laser and task portion. We fit individual models for each pair of kinematic variable and brain region, where kinematic variable is peak speed, path length, or duration and brain region is CFAcl, CFAil, RFAcl, RFAil. The generic form of this model is thus: *(Kinematic Variable) ~ 1 + Laser*Condition + (1|Mouse) + (1|Mouse:Day)*. From these models, we determined the effect of inactivation of different motor cortical areas on segment kinematics; a summary of these model fits along with their specific formulation is given in Tables 1–8.

We use a similar model structure to analyze the statistics of segment complexity. In this case, we do not add a fixed effect for hold vs reach segments, so the model is of the form: *(Measure of Segment Complexity) ~ 1 + Laser + (1 | Mouse)*. We use tortuosity and number of acceleration peaks as measures of segment complexity (discussed above), and thus fit individual models for each combination of brain area and the two complexity measures. A summary of these results is given in Tables 10–11.

For the analysis of submovement kinematics, we found that a linear mixed-effects did a poor job of capturing the structure of our data; as an alternative, we performed a Wilcoxon rank sum test on measures of submovement kinematics and complexity, separating data by both brain area and movement type (hold vs reach), and then aggregating data at the day level. The results obtained from these rank sum tests are given in Tables 12–14 (submovement kinematics) and Table 16 (fraction of overlapping submovements). Finally, to analyze the statistics of both segment and submovement direction, we applied a Watson U^2^ test to cortex intact vs cortex inactivated data. The Watson U^2^ test is used to test the null hypothesis that two samples of data with periodic boundary conditions (i.e. angle data) are drawn from the same distribution--in the case of a significant *p* value, we can reject the null hypothesis and conclude that the two samples are likely drawn from different distributions. Importantly, this test only assesses the similarity of the sample distributions and does not provide information about the magnitude or type of difference between the two samples. In the case of this direction analysis, we separate primitive data for testing based not only on the basis of 1) hold vs reach and 2) intact vs cortex inactivated, but also by the direction that the animal has learned to reach during that particular day. At a coarse-grained level, we thus separated primitive direction data by whether the animal was being asked to reach to the lower left or lower right. We performed the Watson U^2^ test on direction data for both segments and submovements for each of the 16 possible combinations of data types ((4 brain areas) × (2 movement types) × (2 rewarded reach directions)). These results are summarized in Tables 9 (segment) and 15 (submovement).

## DATA AND SOFTWARE AVAILABILITY

All the software (trajectory analysis, real time control software, automation analysis) and the hardware design schematics, including part files and building instructions are available at: www.github.com/GoldbergLab/RodentJoystick

## SUPPLEMENTAL INFORMATION TITLES AND LEGENDS

**Figure S1.**
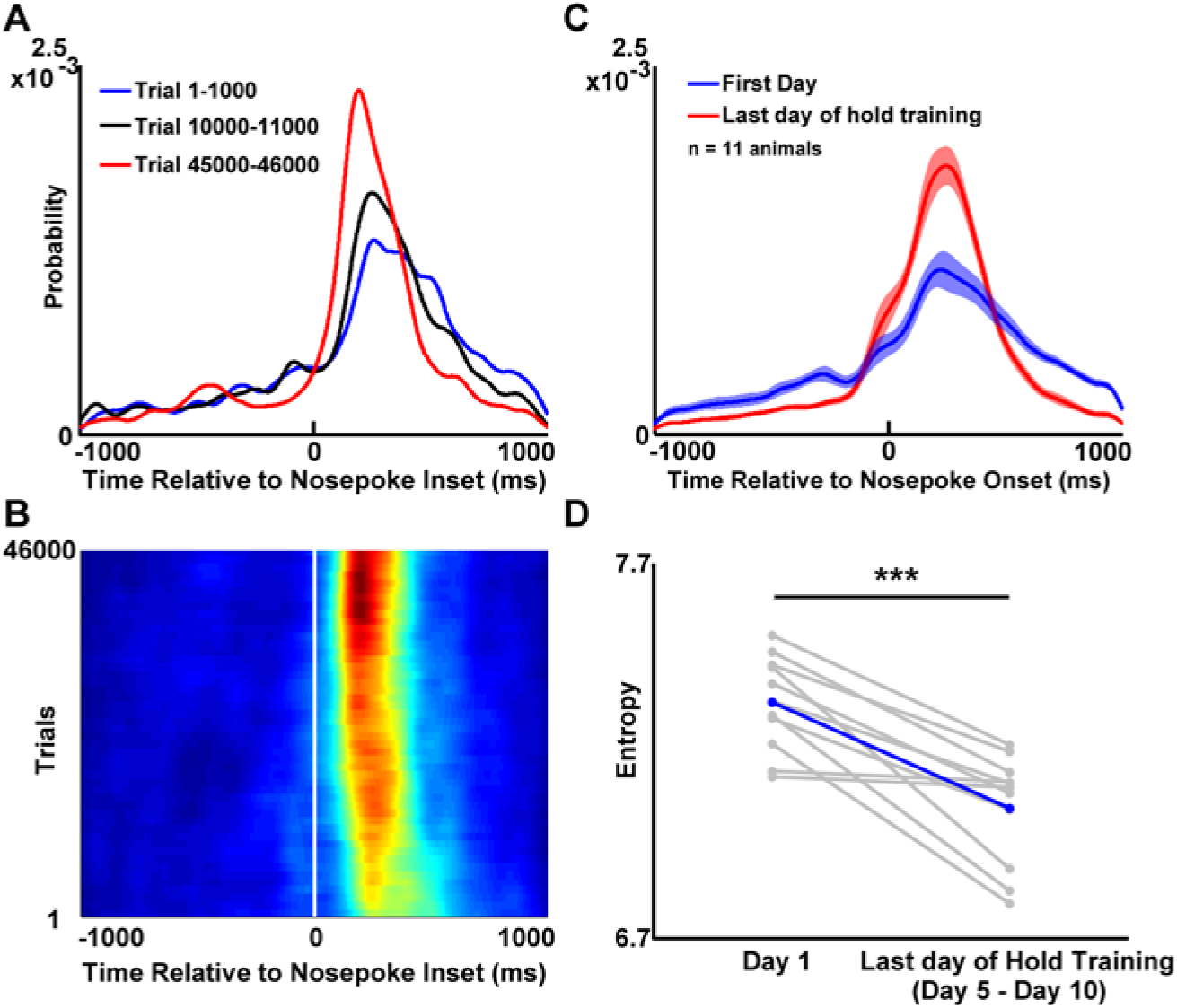
Mice learn to contact the joystick after the nosepoke. Related to Figure 2. (A) Probability distribution of time of joystick contact relative to nosepoke onset in an example mouse at early (trials 1-1000, red), intermediate (trials 10000-11000, black) and late stages of training (trials 45000-46000, blue). (B) Evolution of stereotypy in joystick contact time. Distribution of joystick contact times relative to nosepoke onset as a function of trial number. (C) Time of joystick contact relative nosepoke across animals (n=11 animals, mean±SEM). (D) Entropy of time of the joystick contact relative to nosepoke on Day 1 in the homecage and the last day of hold training (range: 5-10 days later across mice).

**Figure S2.**
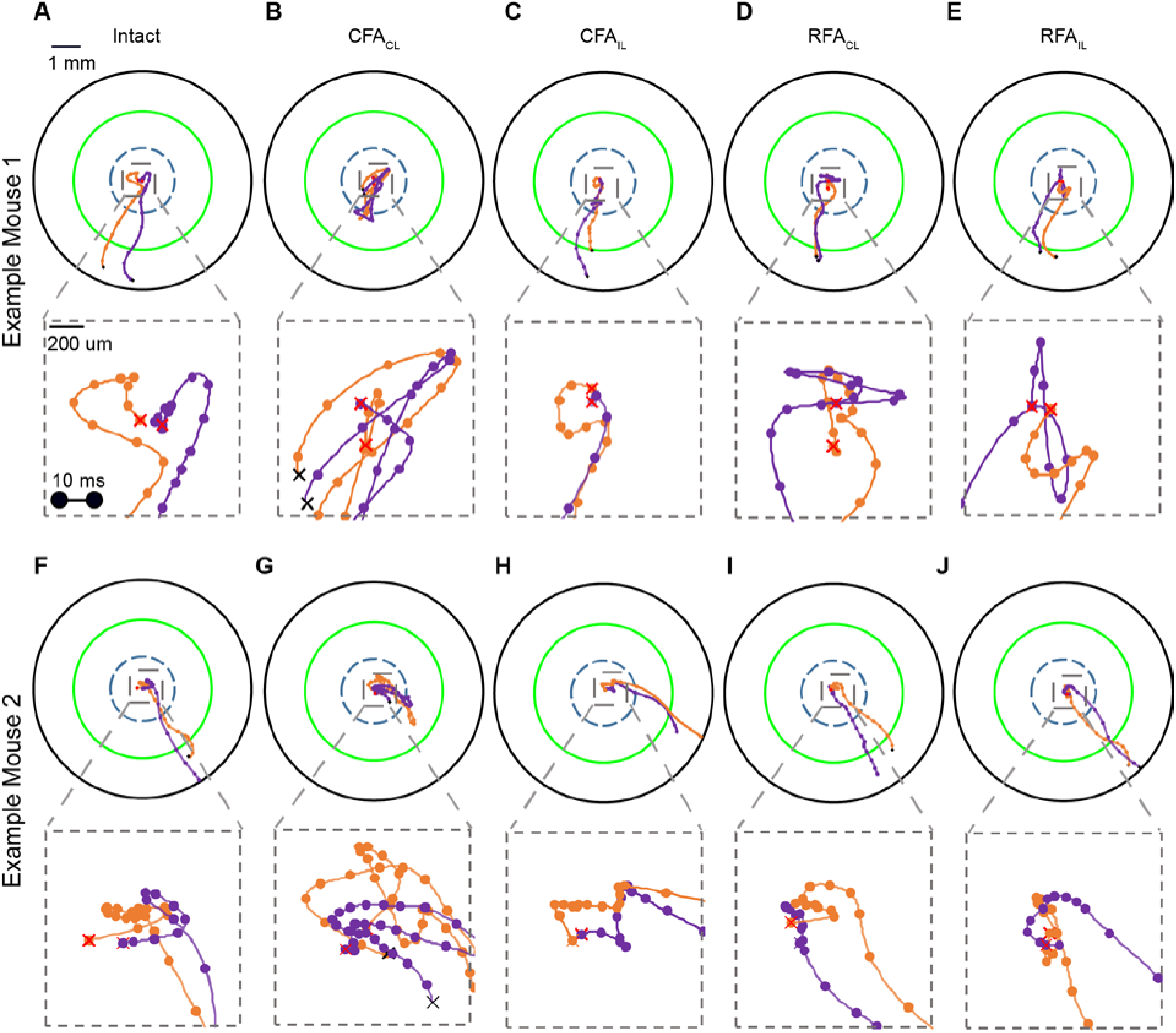
Example trajectories forelimb trajectories. Related to Figure 3. (A) Top: Two trajectories (orange and purple) from an example mouse from control trials. Bottom inset: expanded view of micromovements executed during the hold period. (B-E) Data plotted as in A showing example trajectories from the same mouse with CFAcl (B), CFAil (C), RFAcl (D) and RFAil (E). (F-J). Data plotted as in A-E for a different mouse.

**Figure S3.**
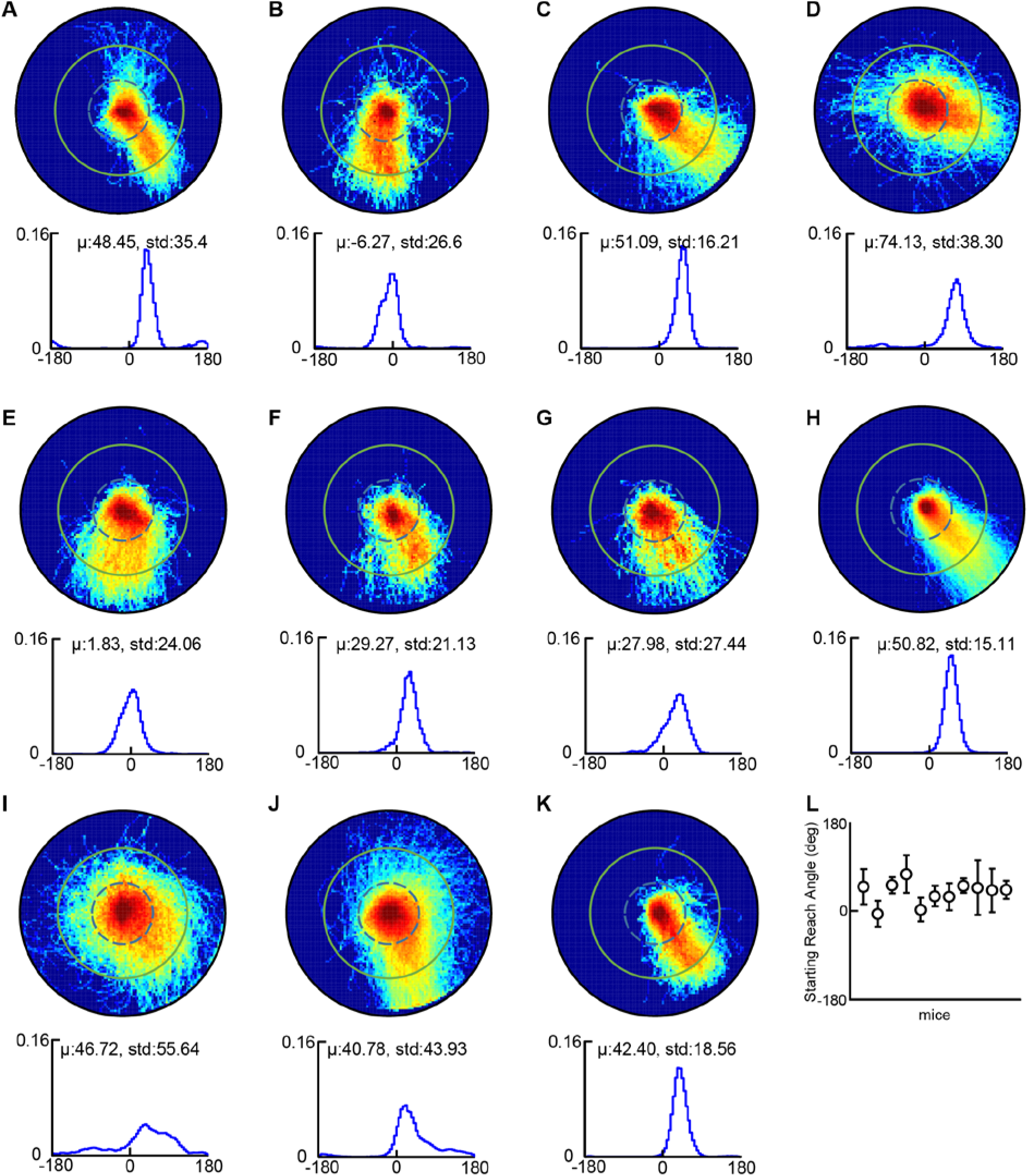
Trajectory patterns exhibited by mice during all-directions-rewarded stage of training. Related to Figure 4. (A) Top: Probability density function of all trajectories produced by a single mouse during a single day, immediately after hold training, when all directions were rewarded (green circle indicates reward zone). Bottom: corresponding reach direction distribution (down corresponds to 0 degrees) (bottom). (B-K) Data plotted as in A for all mice in the study. (L) Reach angles (mean± S.D.) produced by each mouse in the study at all-directions-rewarded stage shown in A-K.

**Figure S4.**
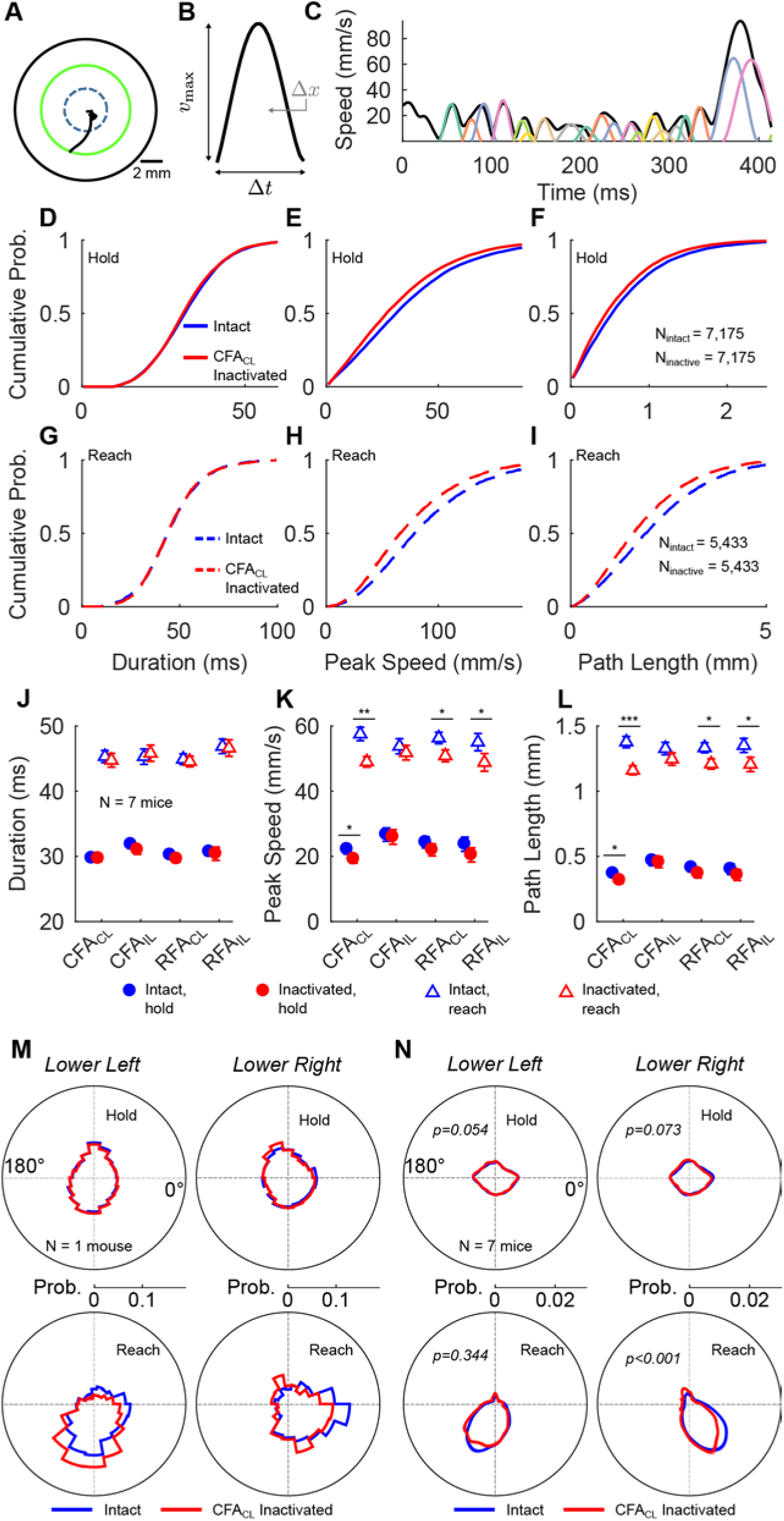
Effect of motor cortical inactivations on the ‘submovement’ class of kinematic primitives. Related to Figure 7. (A) Example mouse forelimb trajectory. (B) Example of a minimum-jerk basis function. (C) Example decomposition of the trajectory in A into minimum jerk basis functions, or submovements, using algorithms previously used in primate studies (Rohrer and Hogan, 2003; Gowda et al, 2016) (see methods).(D-F) Cumulative probability distributions of durations (D), peak speeds (E) and pathlengths (F) of submovements produced within the hold zone. Blue and red traces indicate data intact CFAcl inactivated trials, respectively. (G-I) Distributions plotted as in D-F for submovements that transected the outer radius. (J-L) Mean±SEM of duration (J), peak speed (K) and pathlength (L) with (red) and without (blue) inactivation of various motor cortical areas for hold (circle) and reach (triangle) segments. (M) Direction distributions of hold (top) and reach (bottom) submovements for an example mouse in intact (blue) and CFAcl inactivated (red) trials. (N) Data plotted as in M with average direction distributions across animals (n=7).

**Figure S5.**
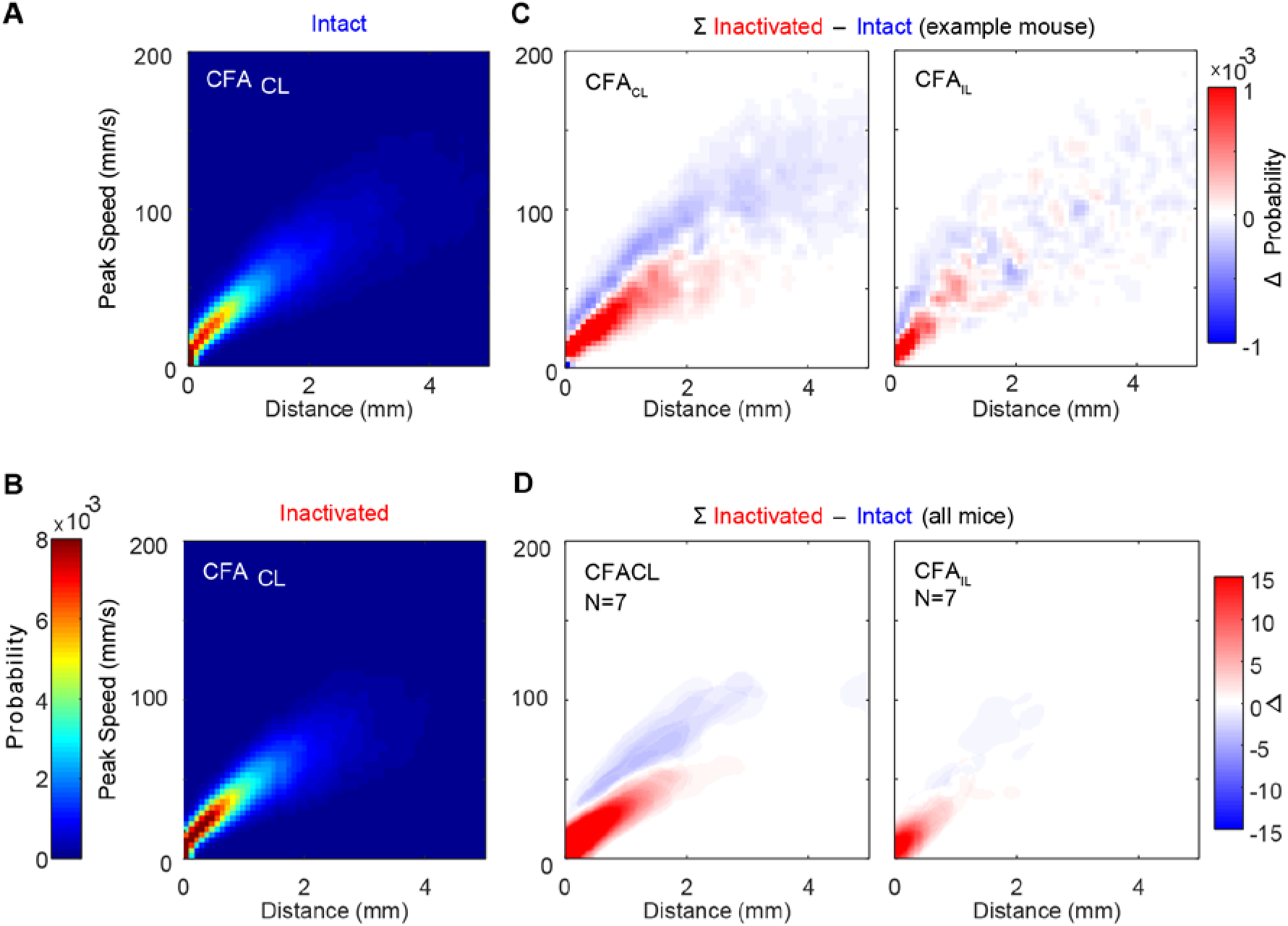
Segment pathlength is predicted by its peak speed. Related to Figure 7. (A-B) Joint probability distributions of segment pathlength and peak speed in intact (A) and CFAcl inactivated (B) trials for an example mouse. (C) Difference between joint distributions of intact and inactivated trials for an example mouse (CFAcl left, CFAil right). (D) Data plotted as in C, overlaying difference histograms across all mice (n=7).

**Figure S6.**
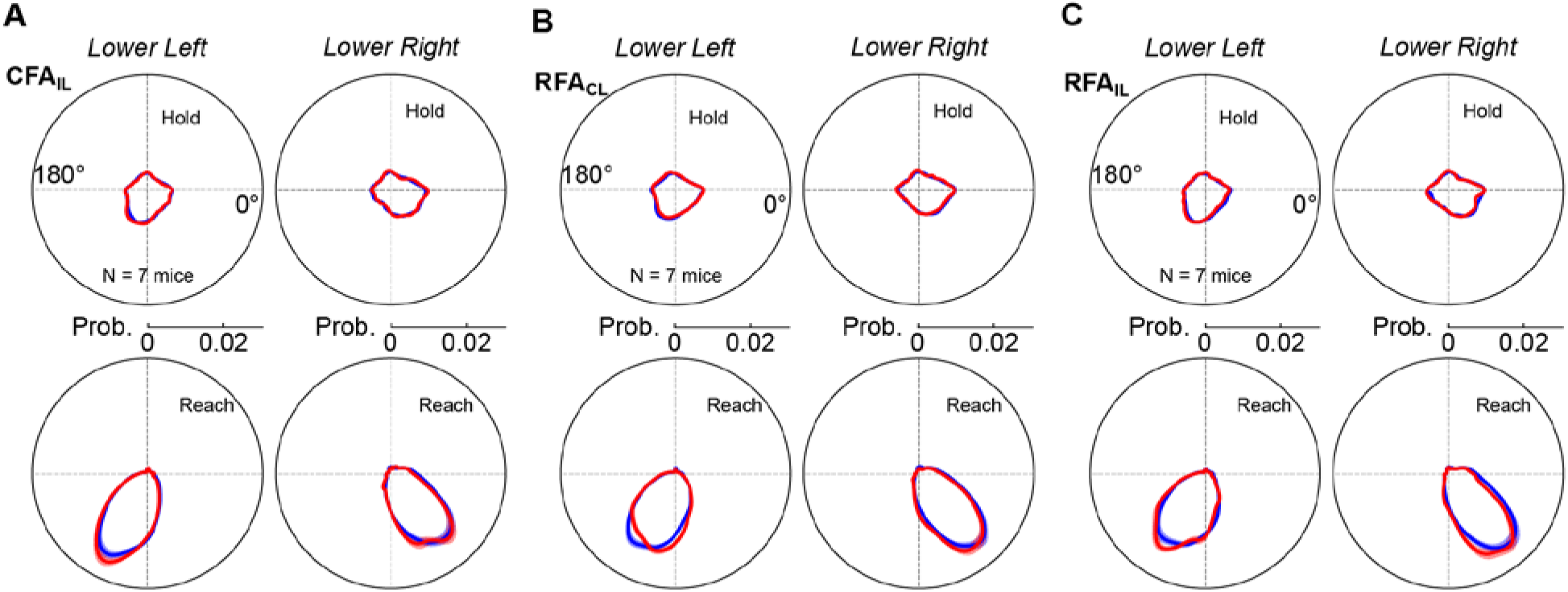
Impact of cortical inactivations on segment directions. Related to Figure 7. (A) Direction distributions of hold (top) and reach (bottom) segments for all mice in intact (blue) and CFAil inactivated (red) trials. (B-C) Data plotted as in A for RFAcl (B) and RFAil (C) datasets.

## Supplemental Experimental Methods

### Instructions for constructing a rodent joystick and a real time homecage behavioral system

#### Introduction

This guide describes in detail the process for building a rack-mountable homecage for running rodent behavioral assays. It includes a water dispensing system, a system for optogenetic (in)activation, a variety of behavioral sensors, including a rodent joystick, and a signal conditioning and data acquisition system. Most of the components are commercially available; we have listed the manufacturer, model, and a link (if available). Some have been custom machined or 3D printed – the specs for these components are made available at the following URL: https://github.com/GoldbergLab/RodentJoystick/blob/master/RodentJoystick-HardwareDesign/

This guide assumes a basic knowledge of soldering (including surface-mount soldering) and handling of electrical tools.

##### Motivation for a Custom Joystick

We considered commercially available joysticks such as: HF46S10 from APEM, the 67A from grayhill and the inductive joysticks from CTI. There were five main shortcomings of these products that made them unsuitable for the study of kinematics in mice.

1. Isometric force profile: All the industrial joysticks have a recentering mechanism that relies on two compression springs set along orthogonal directions. This always leads to moving along the diagonal being up to 40% harder than along the axes of the springs. We solved that problem by using one set of centering magnets right below the joystick.
2. Low displacement force: The industrial joysticks have a displacement force of about 100-250 gf(gram force), ~2.5 Newtons of force, this can be reduced by having a long lever arm. For example, you can lengthen the lever arm by 10 times to get into a 0.1-0.2N range, however, this still puts a lot of load on mouse forelimb kinematics. The centering mechanism we use gives us 0.0081 N/mm, or about 32 mN to move the full range of the joystick, this allows us to get get the fine details of motion even when the animal is holding still (deca micron scale movements) and finer resolution when the mouse is making large (~ 1 mm scale) movements.
3. Future Force Controllability: Our design supports the dynamical control of displacement force on the joystick. By switching out the centering magnet with an electromagnet, we can dynamically control force in real time, and coupled with our real-time microsecond FPGA, we can make that force dependent on a behavioral variable like speed.
4. Built-in touch sensor: Off the shelf joysticks do not provide a readout of when the animal is actually holding the joystick, complicating attribution of displacement by potentially confusing active (animal-induced) versus passive motion.
5. Dead Space: Some commercial joysticks are intentionally designed (in hardware) to have dead-space. i.e. there will be no reading if the joystick is moved in within a certain distance. Presumably to protect against accidental movement of the joystick.

#### Part I: Constructing a joystick

##### Full list of Materials

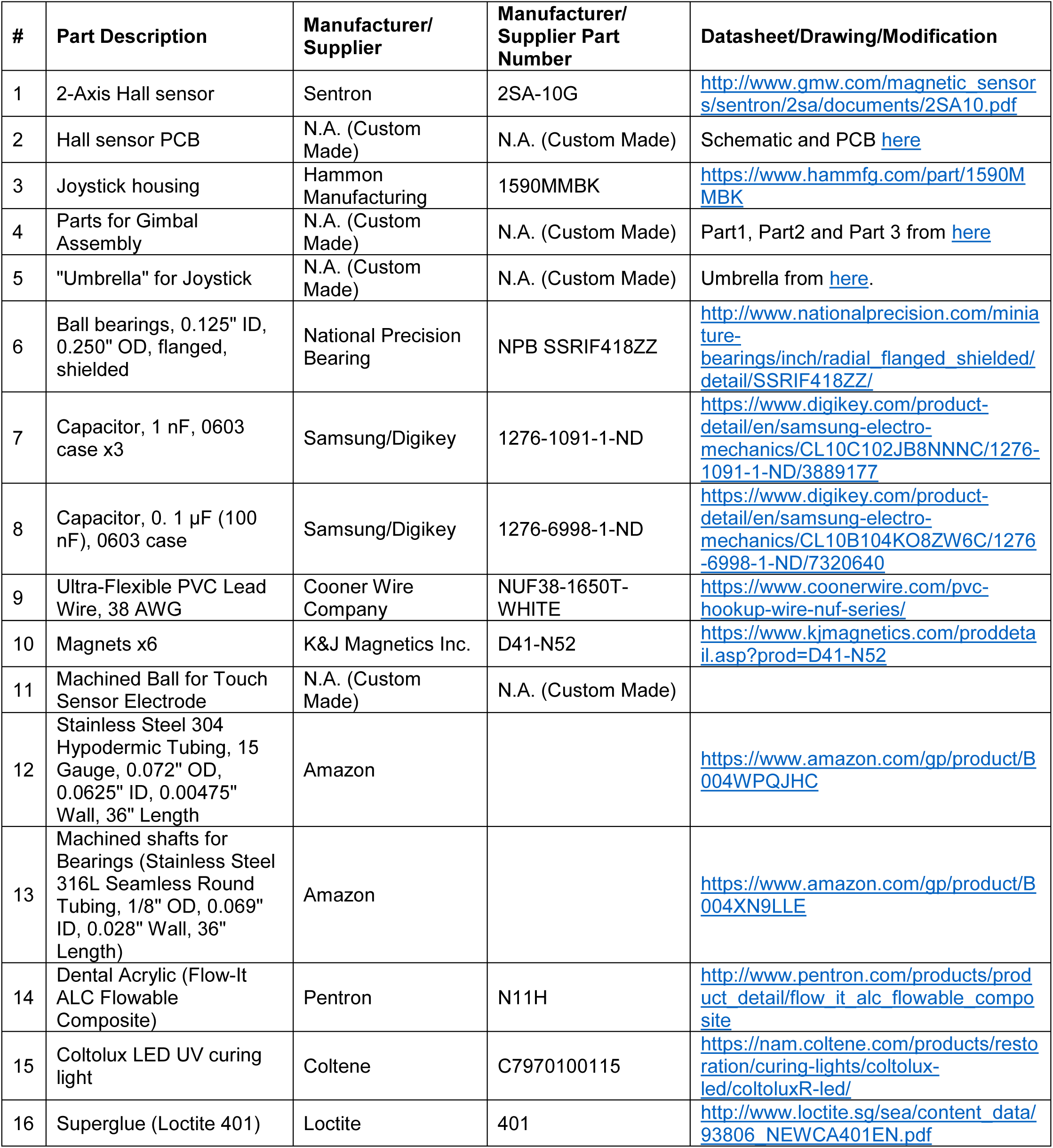
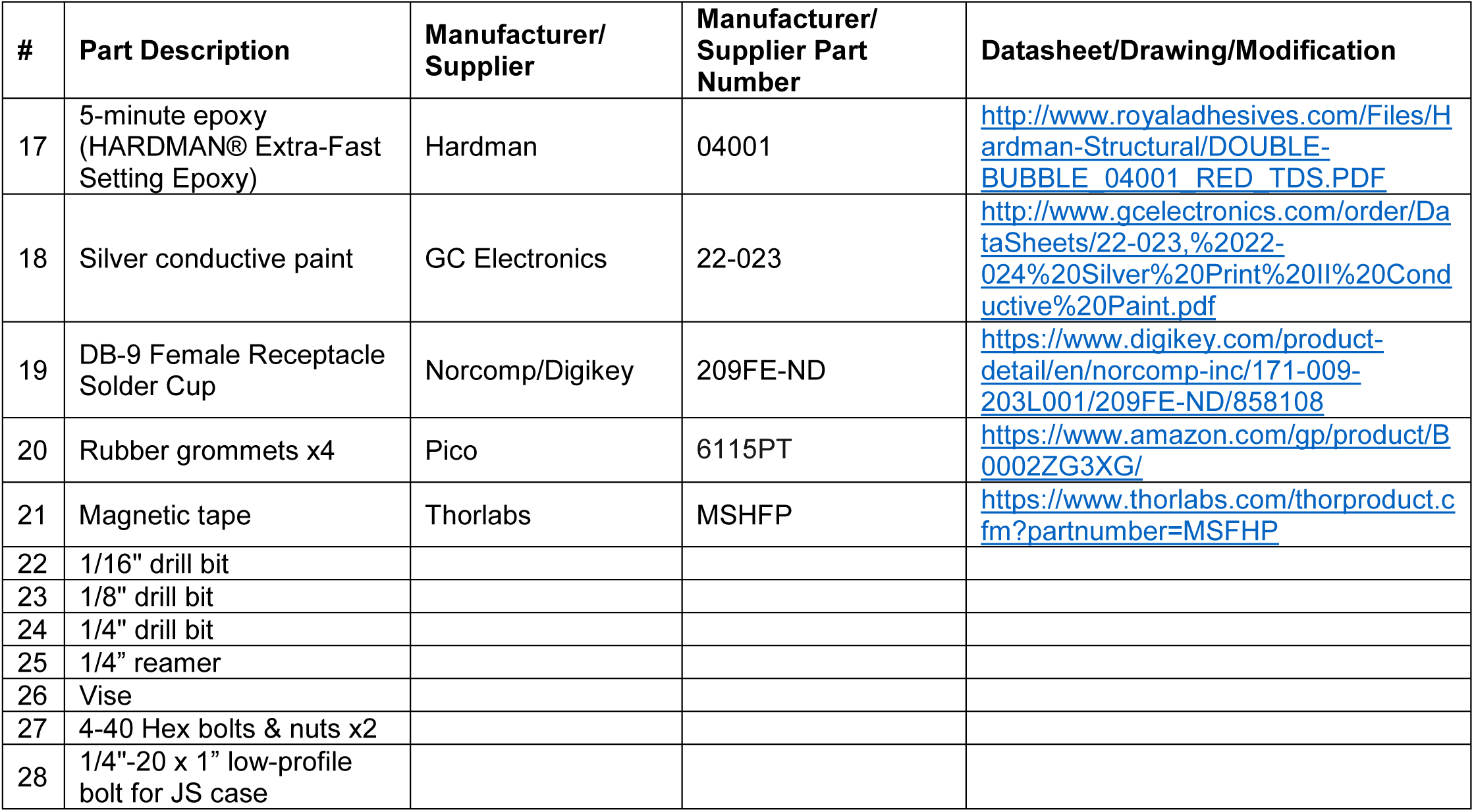

##### Building Instructions

###### General notes

Assembling a usable joystick requires a very high degree of attention to detail and care. As noted below, several of the parts need to be very carefully aligned at every step in the process, or the joystick will not produce a linear, symmetrical voltage vs. position relationship for the full range of motion, and may not provide a smooth force vs. displacement relationship for the animal. We recommend making several joysticks in parallel, and more than you need, since it is likely that at least some of the delicate steps and parts will fall victim to small errors. After the initial learning curve, experienced builders find that around 4 out of 5 joystick builds result in a high enough quality joystick to deploy.

###### 1. Assembling hall effect sensor

The first step in making the joystick is populating a Hall sensor base PCB with capacitors and the Hall sensor IC. The schematic and the board layout are linked in the Materials section.

###### Materials needed for this section

– Hall sensor base PCB
– 0.1 uF (100 nF) capacitor, 0603 case
– 1 nF capacitor, 0603 x3
– Hall Effect sensor chip (2SA-10G)
– 38 AWG Cooner wires x4 (~3” long, stripped and tinned at both ends)
– Dental Acrylic

**Figure 1.**
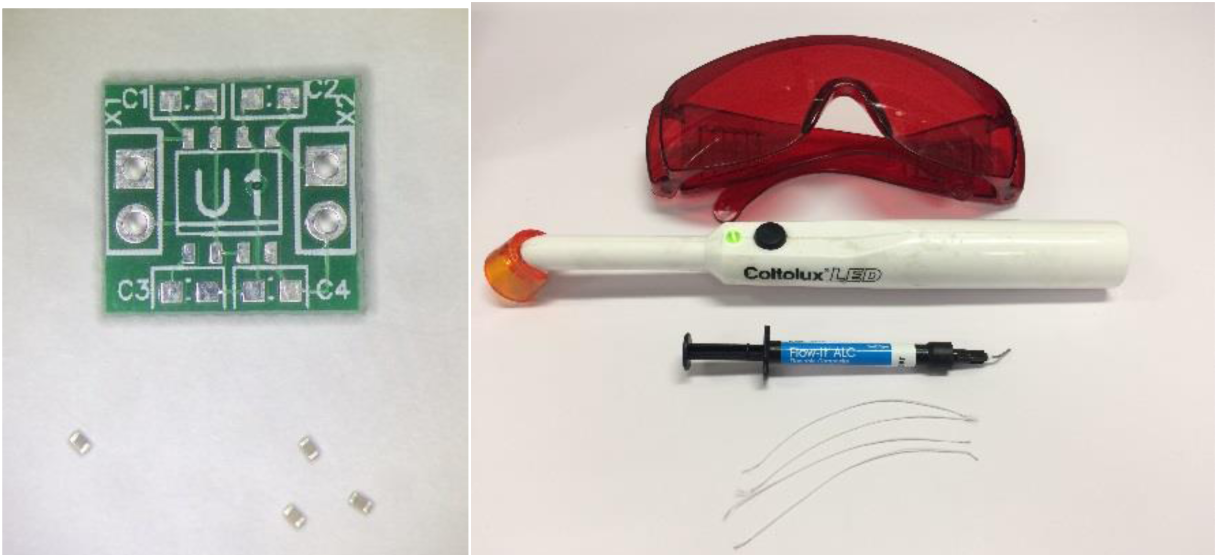
Hall sensor materials. Clockwise from top left: Hall sensor base PCB, dental acrylic glasses, UV lamp, and dispenser, Cooner wires, 1 nF capacitors, 0.1 uF capacitor

###### Instructions

Solder capacitors and Hall sensor chip onto Hall sensor board
– C1 = 0.1 uF
– C2 = C3 = C4 = 1 nF
– **Ensure that hall chip is oriented so chip edges are very close to parallel with the PCB edges (**Figure 2)
– **Ensure that hall chip is very close to horizontal against the PCB (**Figure 3)

Add leads
– Fill the four through-holes on the board with solder
– Re-melt the solder and insert the bare end of a wire up to the insulation in each hole. Orient the wire so it is pointing laterally away from the PCB as shown.
– Cover entire solder joint & a little of the wire insulation with dental acrylic
– Cure it with the UV light

**Figure 2.**
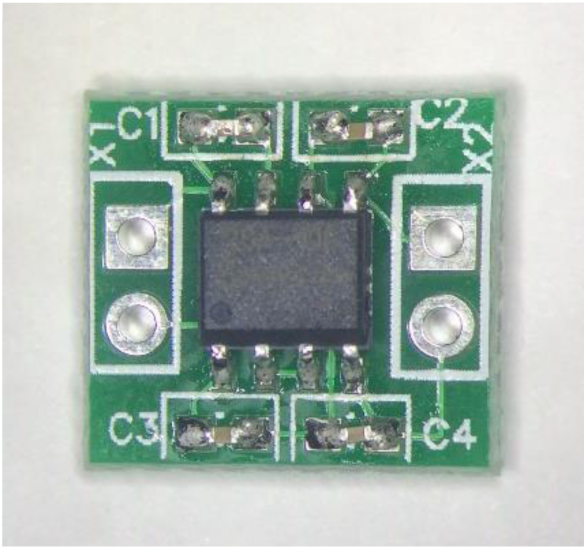
Hall sensor PCB with components soldered. Note the careful alignment of the Hall sensor IC

Trim excess wire/solder from bottom of board
– Board must lay flush on gimbal platform
– If board doesn't sit flat, run the bottom gently over a file a few times
– CAREFUL not to damage the board traces on the back
– See Figure 4 for the finished Hall sensor board

**Figure 3.**
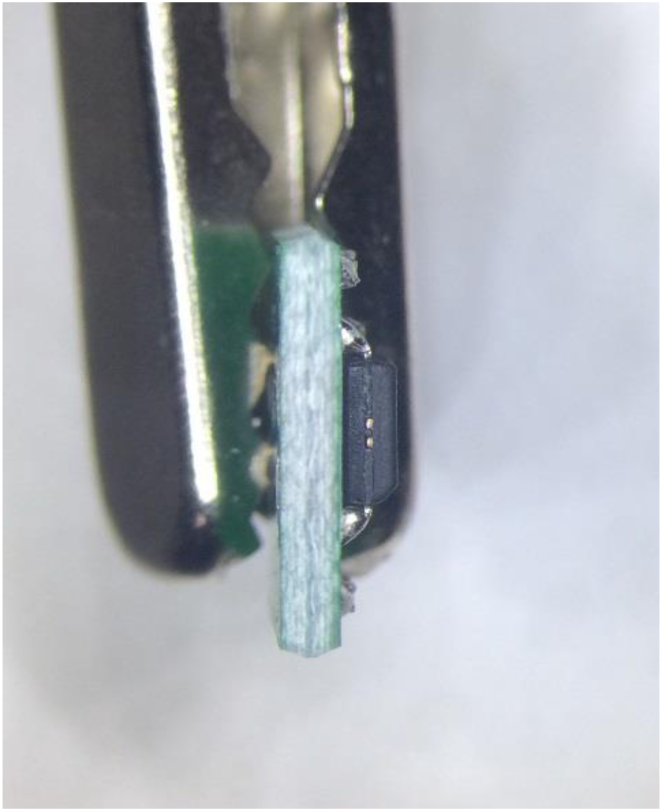
Hall sensor PCB with side view of chip. Note the careful alignment of the Hall sensor IC.

**Figure 4.**
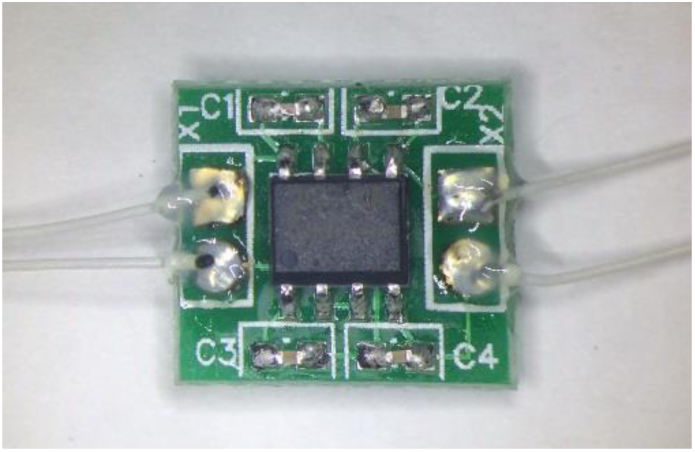
Hall sensor with leads soldered and coated in acrylic

###### 2. Making the Joystick Electrode

The next step is to create a joystick electrode. This electrode will consist of a metal tube with a small electrically isolated ball at the top. Two leads that run through the tube will eventually connect the ball to the joystick PCB via a DB9 connector. The electrode serves as the joystick manipulator as well as a capacitive touch sensor.

###### Materials

– Machined capacitive ball
– Cooner wire
– 1/8” steel tube
– 5-minute epoxy
– Silver conductive paint
– Dremel with
– 1/16” drill bit, abrading bit, cutting wheel bit
– Calipers
– Multimeter
– Tweezers

###### Instructions

Filing rod to length
– Measure 2.15” of the tube
– Secure tube gently in vise, being careful not to crush it
– Use dremel saw to saw off tube
– Use a metal file or a dremel with an abrading bit to remove material from end until tube has a length of 2.10 +/- 0.01”
– Make sure ends are free of metal burrs that could damage the wires

**Figure 5.**
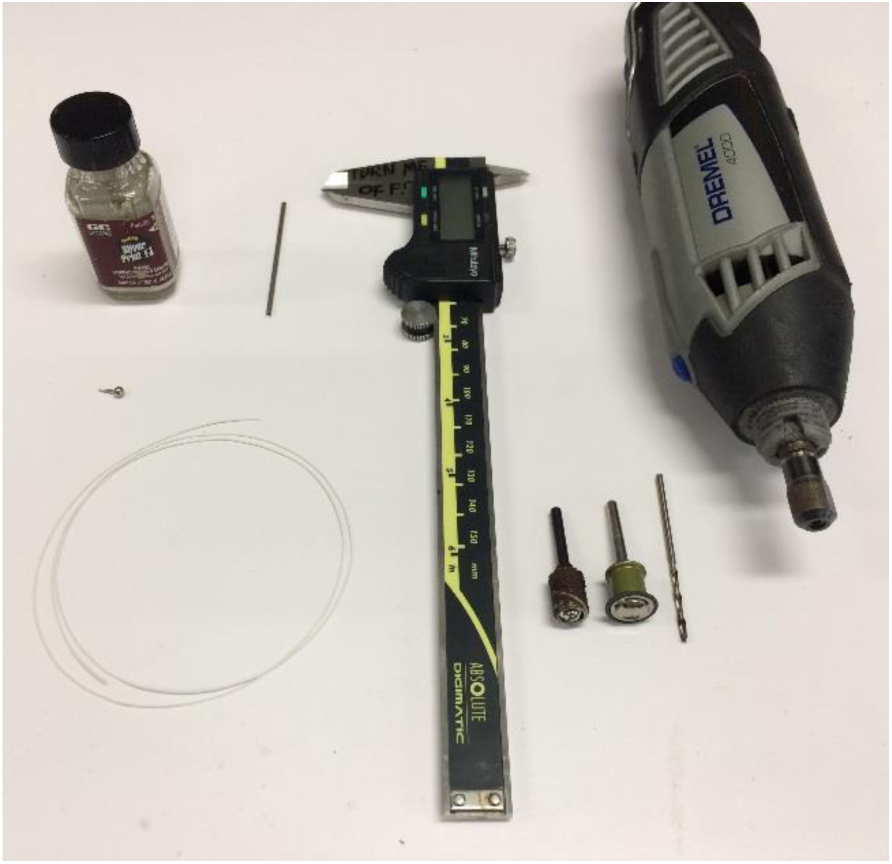
Joystick electrode materials. Clockwise from top left: silver paint, tube, calipers, dremel, dremel bit, cutting wheel, abrading head, Cooner wires, capacitive ball. Not pictured: epoxy

Cutting a side hole
– Fix tube gently in a vise
– Use 1/16” dremel bit to drill hole in **only one side** of tube approximately 1/2” away from bottom of rod
– Use a small file/blade/tweezers to remove any burrs from the edge of the hole that might interfere with or damage wires
– **Careful – tube is weakened at the side-hole spot**
– See Figure 6 for finished joystick electrode tube

**Figure 6.**
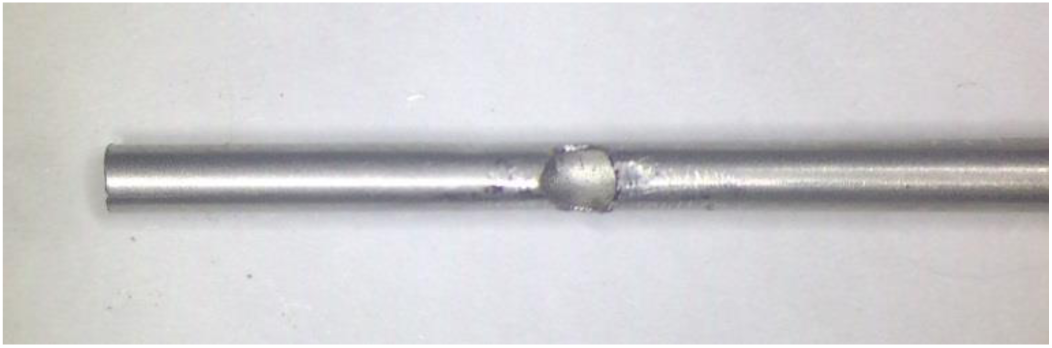
End of the joystick electrode tube, showing a completed side hole. Note the absence of sharp burrs on the edge of the hole.

Stripping, joining, and tinning electrode leads
– Cut two 6” lengths of wire
– Remove approximately 3/16” of insulation from both sides of the wires (suggested tools: tweezers or soldering iron tip)
– **Check that conductor is not damaged by stripping**
– Twist the bare ends of the two wires together on one side
– Tin the two free ends of the wires, and the one twisted-together end.
– Make sure the twisted-together & tinned end of the wires is not too large to fit in tube of the capacitive ball.
– See Figure 7 for completed electrode leads

**Figure 7.**
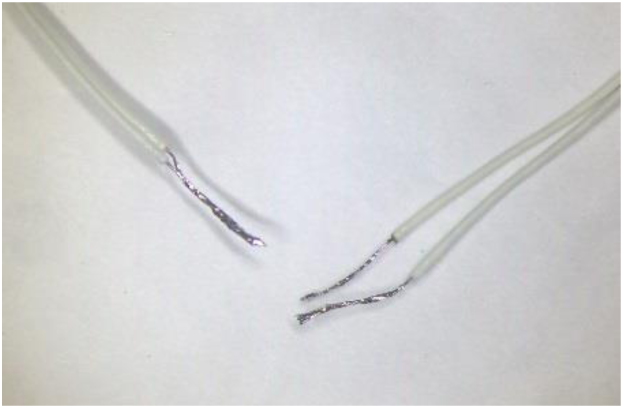
Joystick electrode leads completed, showing the wires twisted together at one end

Inserting wires into capacitive ball using silver paint
– First check that twisted & tinned wire pair fit into the neck of the capacitive ball.
– Shake up silver paint to mix
– Coat tinned wire with silver paint
– Paint tip of capacitive ball neck with silver paint
– Insert wires into neck of the ball
– Wait for paint to dry
– Note that paint alone will NOT result in a strong mechanical connection, so handle carefully
– See Figure 8 for result of this step.

**Figure 8.**
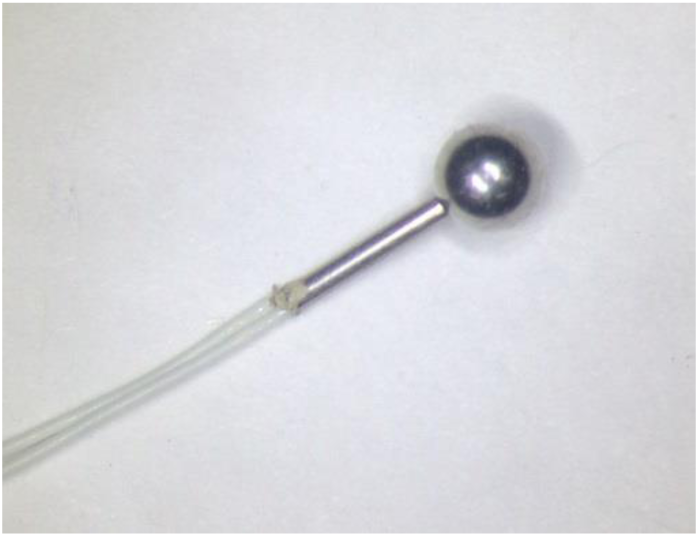
Capacitive ball with leads inserted

Apply epoxy to strengthen and insulate joint
– Mix small amount of two-part epoxy (equal parts) with a fine-tipped applicator like a pin
– Apply epoxy to joint between wires and ball neck
– Spread along wires and barely up to the ball
– Allow epoxy to set while ball is upside down (leads pointing upwards). This allows gravity to pull epoxy slightly onto the base of the ball, which helps prevent contact between the ball and the electrode tube.
**-** **Ensure that epoxy electrically insulates the neck of the ball so that it will NOT make electrical contact with the tube. ALL silver paint and metal below ball must be covered**
– **Ensure that epoxy does not stray TOO far up the sides of the ball itself; this could interfere with touch sensing**.
– **Ensure that epoxy layer doesn’t have bulges that will prevent the neck of the ball from inserting into the electrode tube**. It is possible, though difficult, to trim the epoxy layer later if bulges form during drying.
– See Figure 9 for result of this step.

**Figure 9.**
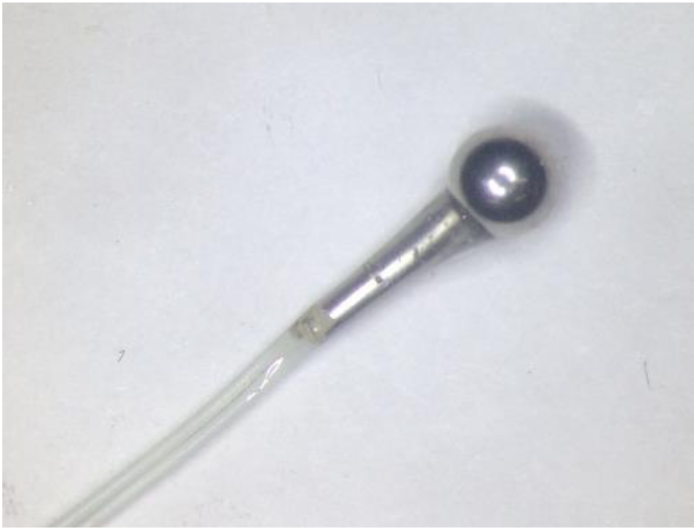
Capacitive ball with leads and a coat of epoxy. Note that the epoxy spreads out enough at the base of the ball to prevent it from contacting the tube when inserted, but not so much as to encroach on the touch-sensing area on the top and sides

Threading wires through tube
– Use cutters to eliminate any extra overhanging epoxy
– Thread wires through electrode rod – insert them on far side of the tube from the side hole
– Use tweezers to pull wire out through side-hole
– Gently pull wires until the ball settles onto the end of the tube
– If epoxy prevents the ball from getting close to the end of the tube, gently trim obstructing parts off.
– Pull ball slightly out again, and add 2nd layer of epoxy in same place as the first layer. **Do not wait for it to set**.
– Pull gently on the wire leads to reseat the ball at the end of tube. Push on the ball at the same time to assist.
– Wipe off excess epoxy below ball. **Careful not to wipe epoxy onto touch-sensitive sides and top of ball**.
– Spin around to visually check that the ball is approximately centered on rod. Adjust if it is not.
– **Use a multimeter to check that the two wire leads are electrically connected to the ball, but NOT to the tube**.
– Allow the epoxy to set – make sure gravity does not pull the ball out of place during setting.

Final test of electrode
– Use a multimeter to check that the two wire leads are electrically connected to the ball, but NOT to the tube.

###### 3. Cleaning/drilling out joystick gimbal assembly parts

The next step is to prepare the four parts of the gimbal for assembly. The gimbal consists of a platform that holds the Hall sensor, a cross that helps affix the gimbal accurately and securely in the baseplate, and a rectangle and square rotating pieces that provide the two degrees of freedom via the bearings that suspend them.

###### Materials

– Gimbal platform
– Gimbal square
– Gimbal rectangle
– 1/16”, 1/8”, and 1/4” drill bit
– 1/4” Reamer
– Bearing (for checking fit)
– Electrode tube (for checking fit)
– Magnet (for checking fit)

**Figure 10.**
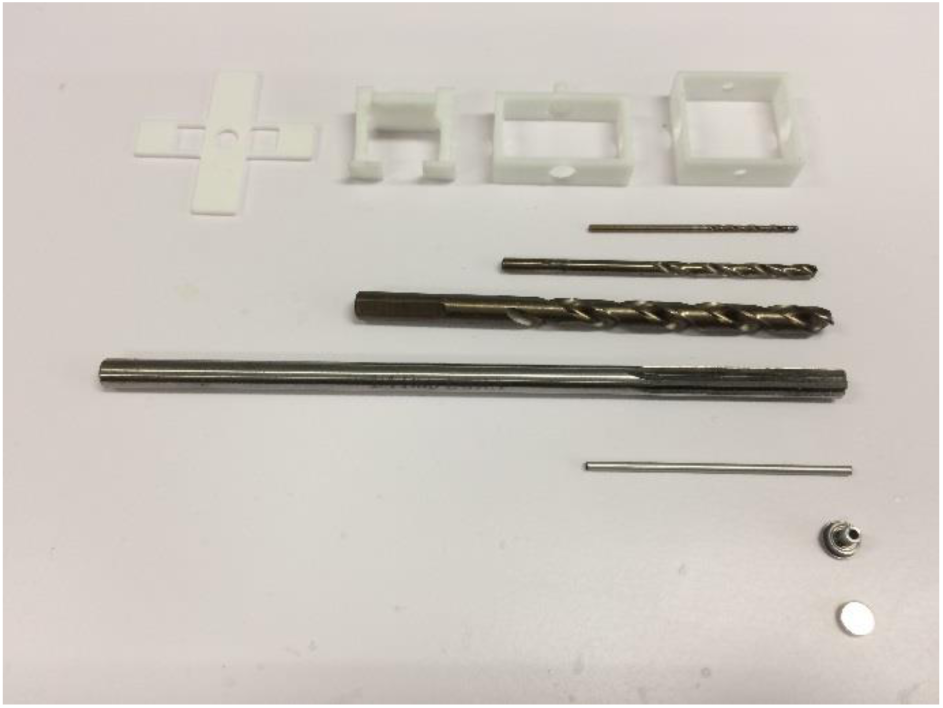
Gimbal preparation materials. Clockwise from top left: Cross (not used until next step), gimbal platform, gimbal rectangle, drill bits (1/16” 1/8” and 1/4”), %” reamer, sample tube, sample bearing, sample magnet

###### Instructions

Clean and inspect 3D printed parts
– The 3D printing process can leave small burrs and other imperfections.
– Examine parts and smooth/remove/fix any imperfections prior to using parts.
– Check that right angles are very close to 90°; significantly warped parts are likely to produce a failed joystick.

Gimbal platform: clearing out holes
– Use 1/8” drill bit **manually** to clear out the two 1/8” hole in the “shoulders” of the gimbal platform.
– Note: Using a power drill can result in a large amount of wobbling of the drill bit that can create holes that are too wide with non-vertical sides. This can introduce fatal levels of misalignment later. Using the drill bit manually is tedious, but more accurate. Using a tap wrench to hold the drill bit can make this easier on the hands.
– Take care not to change each hole’s orientation as you widen it; it’s best to clear out both holes simultaneously with the drill bit to ensure that the two shoulder holes remain coaxial. It is critical that the holes remain closely aligned, or the gimbal won’t rotate smoothly.
– Carefully widen hole until bearing shaft just fits through easily
– See Figure 11 for results of this step.

**Figure 11.**
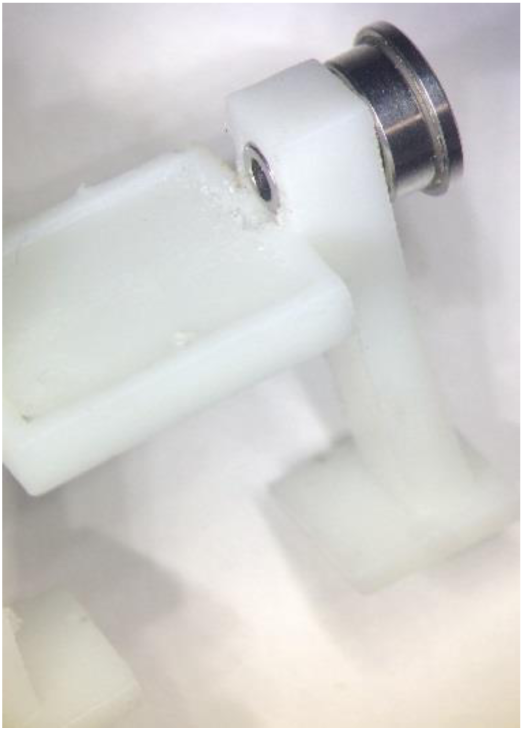
Bearing shaft fits easily into one of the two gimbal platform shoulder holes.

Gimbal square: clearing out holes
– Use 1/8” drill bit manually to clear out the two 1/8” holes in the gimbal square piece **if necessary** (see below).
– As before, drill both holes simultaneously to maintain them in a coaxial orientation
– Carefully widen hole until bearing shaft just fits through easily
– Use 1/4” drill bit to manually clear out the two 1/4” holes.
– Carefully widen holes until bearing fits in **snugly**

**Figure 12.**
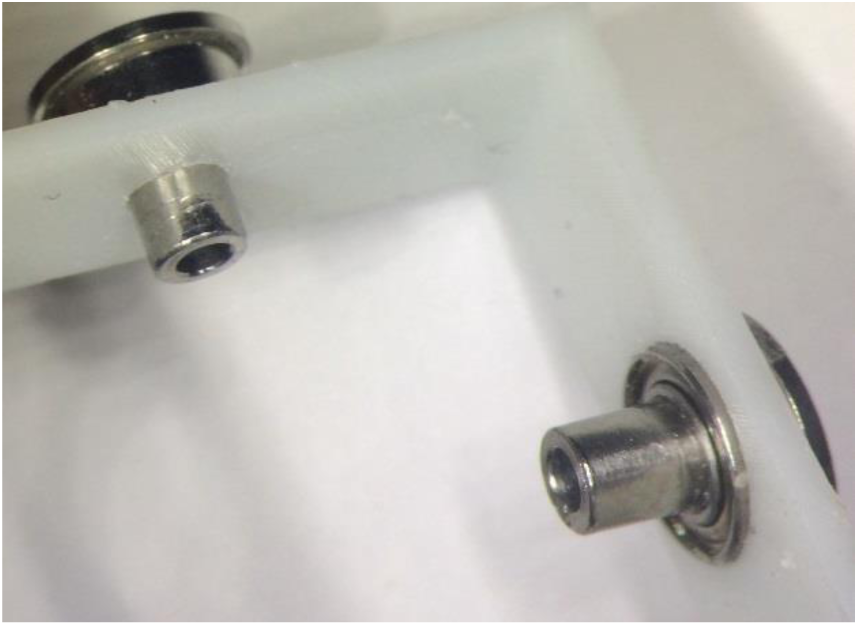
Gimbal square, showing bearings successfully fitting in two of the holes.

Gimbal rectangle: clearing out holes
– Use 1/4” drill bit manually to clear out the three 1/4” holes in the gimbal rectangle piece **if necessary** (see next step).
– Carefully widen short-side holes until bearing fits in **snugly**
– Carefully widen bottom long-side hole until a magnet fits **snugly**
– Clear out hole in electrode post hole with 1/16” drill bit
– **As you clear out the post hole, it must remain vertical or joystick will be non-vertical**
– Make sure electrode post can fit in **snugly**
– Ream out magnet hole on the inside of the rectangle. Be careful to hold the reamer vertically to avoid slanting the holes
– Ream directly through the magnet hole in one side of the rectangle through to the magnet hole on the other side of the rectangle.
– Put magnet in to make sure it fits snugly in each hole

**Figure 13.**
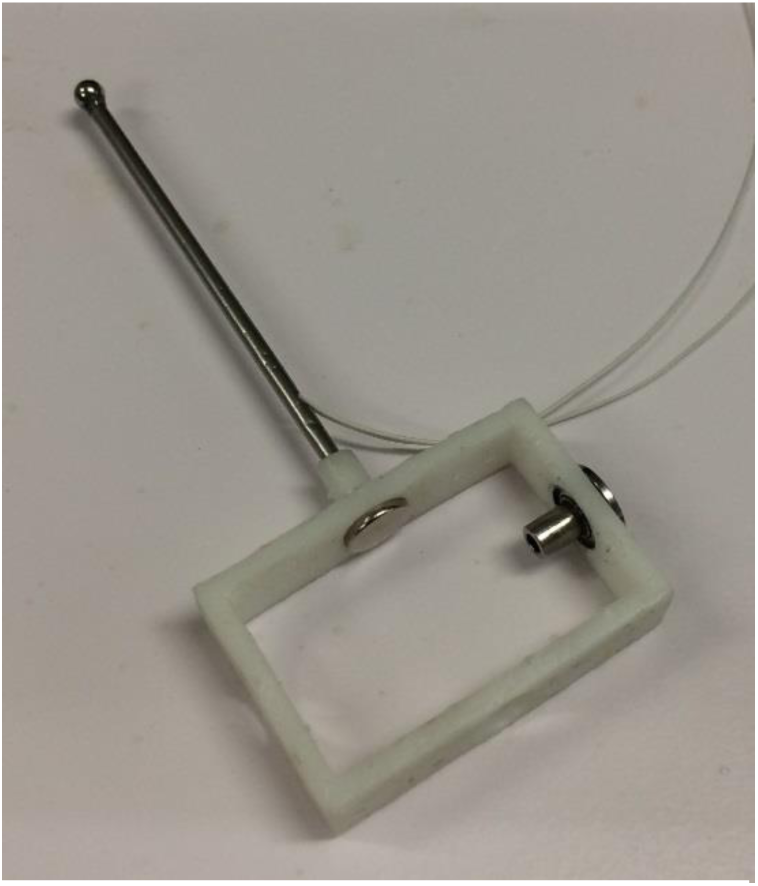
Gimbal rectangle, showing bearing, post, and magnet successfully fitting.

###### 4. Assembling joystick

The final step for creating the joystick is to assemble the previously created parts (the electrode, the gimbal, and the base), connect the joystick leads to the DB-9 connector, and house the joystick in the case.

###### Materials

– Gimbal parts (platform, square, and rectangle, cleaned and prepared)
– Ball bearings with shafts x4
– Completed Hall sensor PCB
– Completed JS electrode
– Magnetic tape
– Magnets x6
– Baseplate
– Plastic cross
– Super glue
– Epoxy
– Glue/epoxy applicator
– DB9 Female Solder Cup
– 4-40 Hex bolts & nuts x2
– JS case
– Case screws x4
– Rubber grommets x4
– Plastic umbrella
– 1/4-20 x 1” mounting bolt

**Figure 14.**
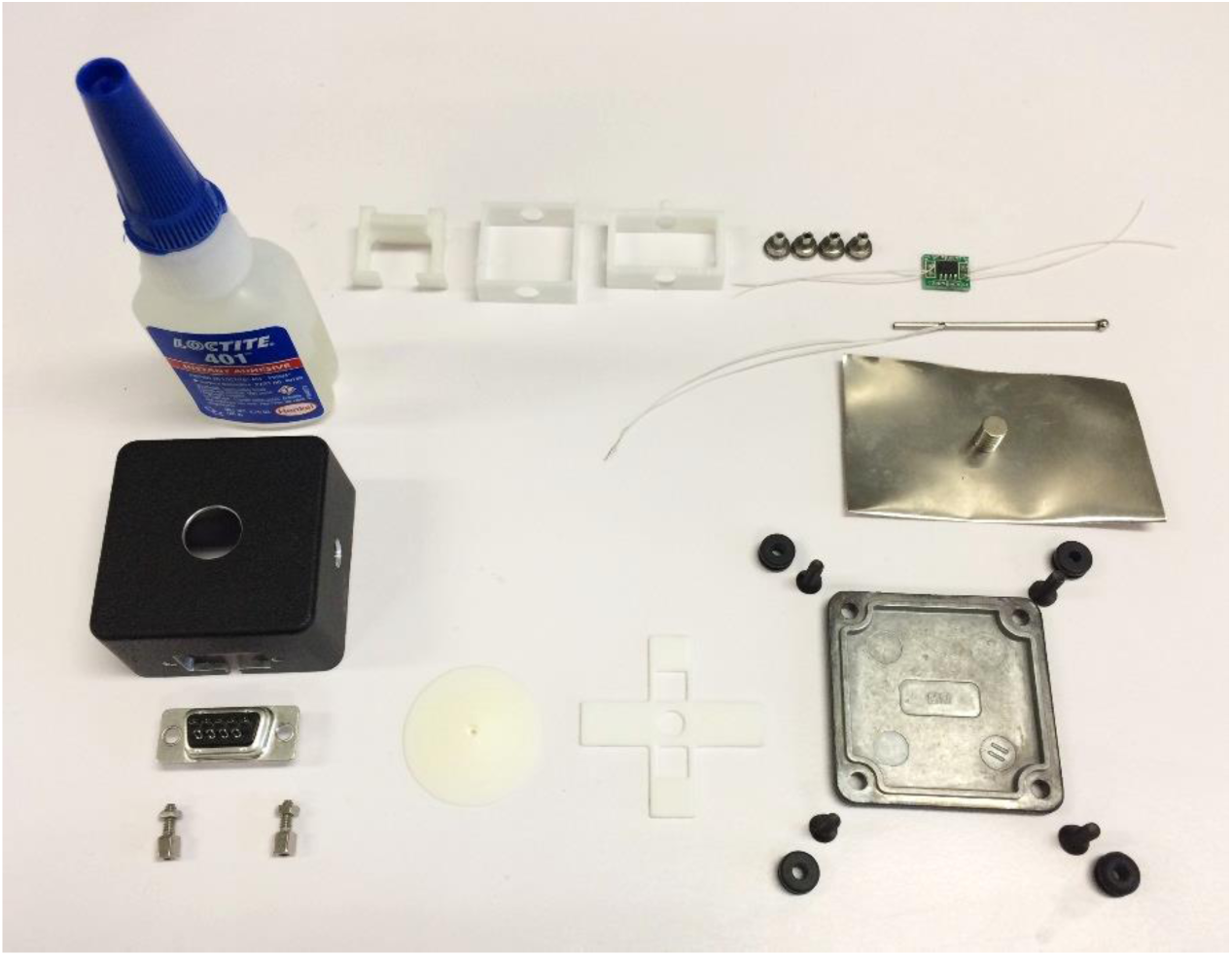
Materials for assembly of joystick. Clockwise from top left: Superglue, gimbal pieces, bearings, Hall sensor, electrode, magnetic tape, magnets, baseplate w/screws & grommets, cross, umbrella, DB-9 connector & screws, and housing top. Not pictured: 34-20 × 1” mounting bolt

###### Instructions

Seating Hall sensor
– Thread the four Hall sensor wires through holes in shoulders of platform gimbal
– Seat Hall sensor board flat on platform
– Mix epoxy
– Add a dab of epoxy in the center of the platform
– Press the Hall sensor down into place as you gently draw the wires through gimbal platform holes
– **CRITICAL:** Adjust the position of the Hall sensor PCB until the Hall sensor IC is perfectly square with gimbal platform in all dimensions. Any significant skewing, tilting, or off-center positioning will result in poor joystick position output. Note that the position of the PCB itself is irrelevant – it’s the Hall sensor chip itself that must be as close to perfectly oriented as possible. It can be helpful to allow the epoxy to partially set before fine-tuning the position of the Hall sensor, as the more-viscous partially-set epoxy will prevent the Hall sensor from shifting after alignment
– Let Hall sensor epoxy set

**Figure 15.**
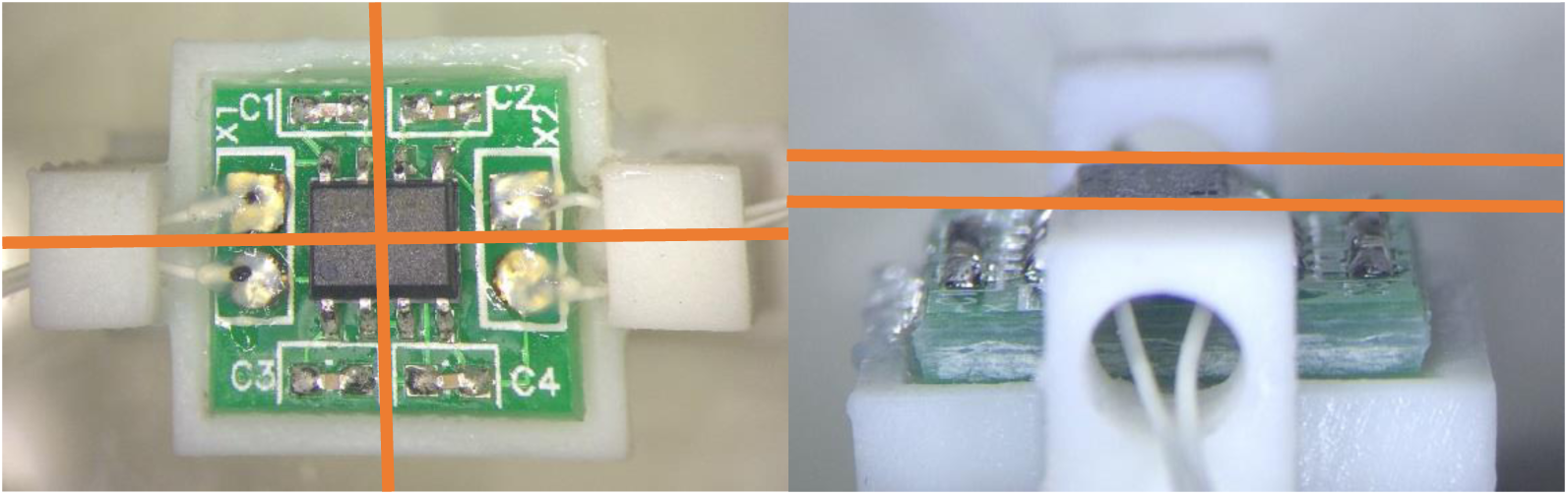
Alignment of Hall sensor chip

Preparing magnetic tape to hold top equilibrium magnet
– Cut small piece of magnetic tape in the same shape as the inside surface of the long side of the rectangle piece of the gimbal (1.10” × 0.32”)
– Place tape on inside bottom of rectangular piece
– Firmly press down magnetic tape until it is adhered to the rectangle piece.
– See Figure 16 to see the results of this step.

**Figure 16.**
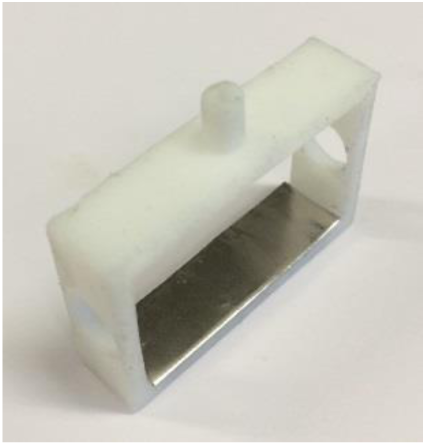
Rectangle piece with magnetic tape

Attaching square & rectangular pieces
– Gently thread the two pairs of Hall sensor wires through the two 1/4” holes in square piece
– Thread each pair of wire leads through a bearing. Insert the bearing into the 1/4” holes so the bearing shaft inserts into the gimbal platform shoulder holes.
– **Ensure square and platform pieces rotate extremely smoothly and freely against each other – you shouldn’t feel any kinks or hitches. You may need to make small adjustments to the position of the bearings and the position of the platform on the bearings to achieve smooth rotation**.
– Insert square piece into rectangle piece, and add bearings to connect
– **Ensure rectangle and square also rotate extremely smoothly and freely against each other**
– See Figure 17 and Figure 18 for results of this step.

**Figure 18.**
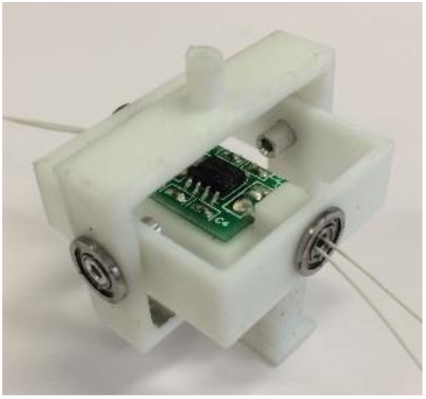
Gimbal square and rectangle suspended on platform

**Figure 17.**
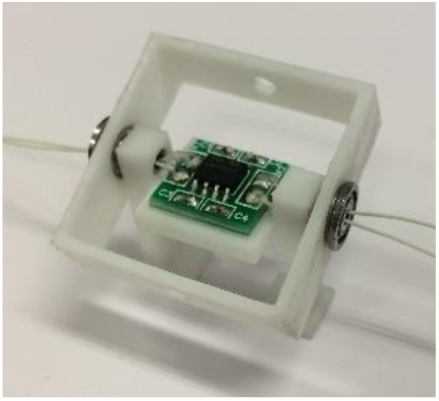
Gimbal square suspended on platform

Mount cross on baseplate
– Seat cross in baseplate
– **Make sure two square holes on plastic cross are not on top of words/markings on baseplate. In the correct orientation, the cross should lay flat. It may be necessary to file down the ends of the cross a little to achieve a good fit**.
– Add superglue with toothpick on edges of cross & baseplate ONLY on sides of cross without holes

**Figure 19.**
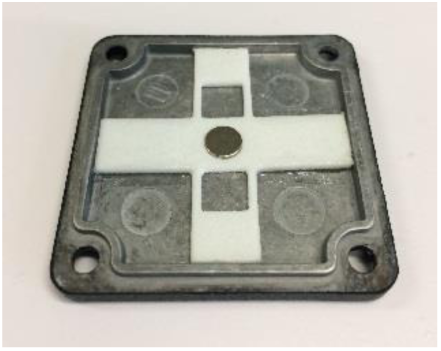
Cross with equilibrium restoring magnet glued into baseplate

Add equilibrium-restoring magnets
– Add superglue on center hole
– Press magnet into center hole
– **Ensure the magnet lays horizontally. Misalignment of this magnet can result in a non-central equilibrium position**
– Add another magnet **IN THE SAME ORIENTATION** as the baseplate equilibrium magnet on the rectangle piece in the hole on the opposite side from the magnetic tape
– Press fit the magnet until it is flush with the rectangle bottom
– Add second magnet on the bottom of the rectangle, to make a stack of two. This increases the equilibrium-restoring force of the joystick.
– See Figure 19 and Figure 20 for the results of this step.

**Figure 20.**
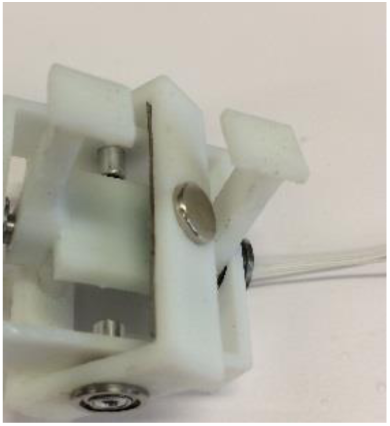
Gimbal rectangle with two equilibrium restoring magnets inserted. The first magnet is not visible.

Seat gimbal on baseplate
– Push-fit gimbal feet into holes in plastic cross in baseplate
– **Ensure that the arms of the platform are vertical and perpendicular to baseplate**
– Superglue down feet of platform
– Wait for superglue to dry
– See Figure 21 for the results of this step.

**Figure 21.**
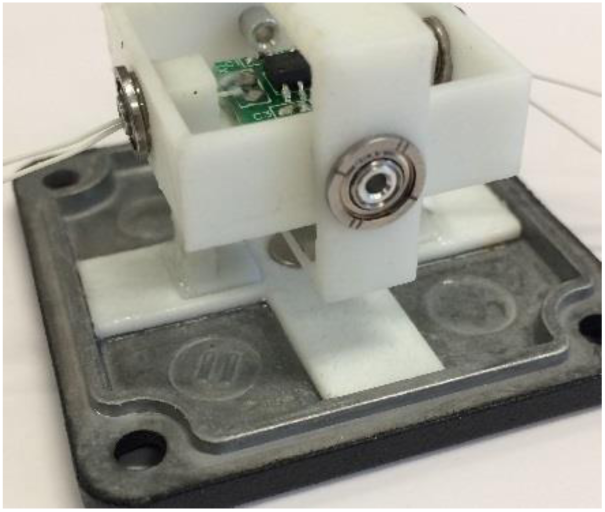
Gimbal seated on baseplate

Add Hall sensor magnets
– Press first magnet firmly into the top hole of the rectangle (this is made less challenging by using a non-magnetic stick to insert the magnet).
– Optionally glue the magnet it in if it isn't staying in with friction alone
– Make sure the magnet surface is parallel to the inner surface of the rectangle piece.
– Add two more magnets (making a total of three) into the top of the rectangle piece. This increases the magnitude of the signal from the Hall sensor, and therefore improves the signal to noise ratio.

Place joystick electrode into rectangle piece
– Insert the bottom (non-ball side) of the electrode into the collar sticking up out of the rectangle piece.
– Make sure that the electrode leads are facing the side of the Hall sensor PCB with capacitors 1 and 2.

**Figure 22.**
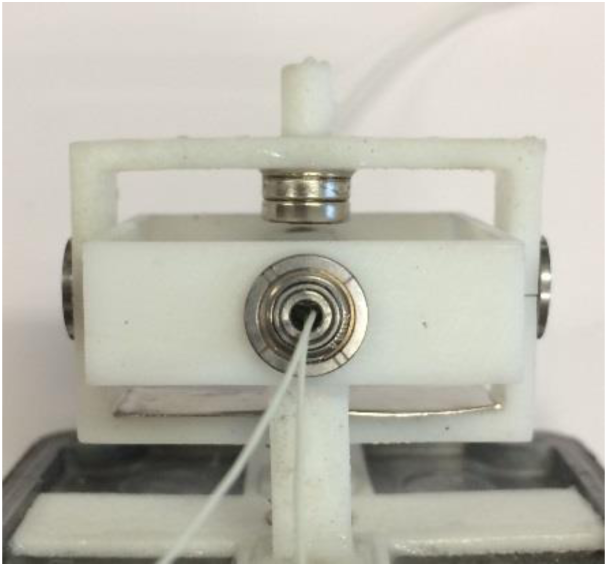
Gimbal with Hall sensor magnets installed

Centering joystick
– **This is a critical, delicate step! The quality of the joystick output is very sensitive to this step!**
– Repeat for each of the two degrees of freedom:
– Check whether the Hall sensor magnets are centered directly above the Hall sensor in equilibrium position
– Also use a ruler to check the verticality of joystick
– Adjust the position of the bearings in the square/rectangle (depending on which degree of freedom you are adjusting) to change position of Hall sensor magnets and angle of joystick.
– Prioritize the position of the Hall sensor magnets over joystick verticality, but maximize both. Some joystick non-verticality can be overcome by adjusting the housing screws later. However, if the Hall sensor magnets are not carefully centered over the Hall sensor at equilibrium, then the position output will be irrevocably distorted.
– Note that you must maintain a tiny air gap between the rectangle and square pieces to avoid introducing more friction into the joystick motion.

**Figure 23.**
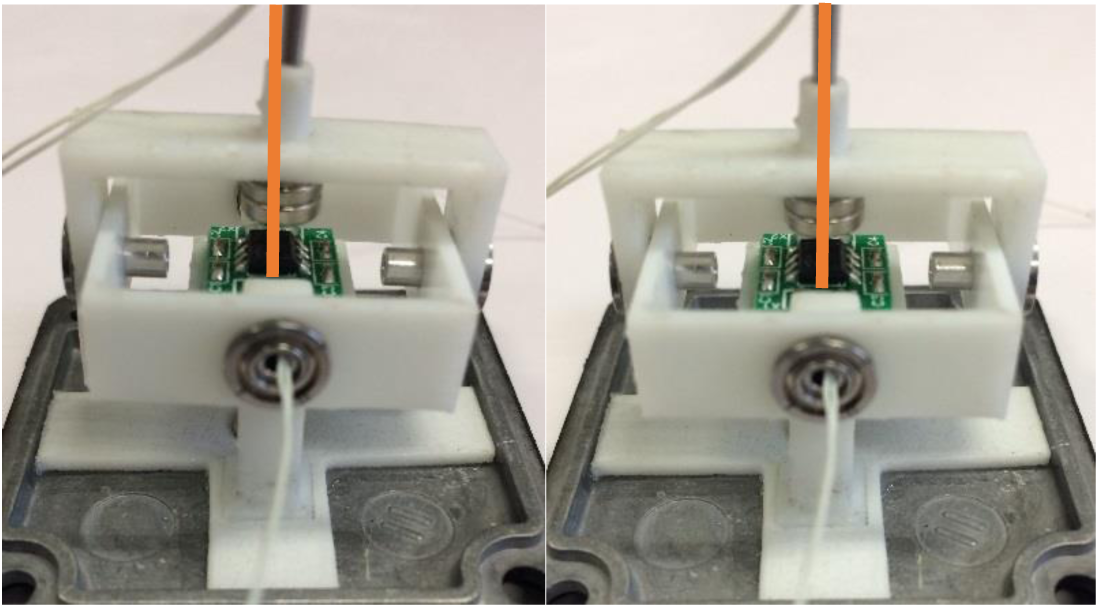
Gimbal centering process (one degree of freedom shown). Before centering (left), after centering (right)

Gluing 1st set of bearings
– Gently apply superglue with outside edge of bearing where it makes contact with square/rectangle (depending on degree of freedom) piece of gimbal
– Use a fine-tipped applicator (a toothpick or similar)
– Gently apply superglue to bearing shafts
– **Be careful not to superglue the inner and outer parts of the bearing together!**
– Allow glue to dry
– Test if joystick is “bouncy” – joystick should rebound & oscillate when disturbed
– Consider retesting that the joystick electrode has appropriate electrical connectivity before gluing it down – this is the last time you can easily replace it if it has become broken.
– Superglue down electrode wires

○ Dab glue on top surface of rectangle, press wires tightly against the electrode mounting collar
– Superglue joint between electrode tube and rectangle piece

Soldering joystick connector
– DB-9 female with solder cups
– Screw in hex screws with nut on either side
– Fill solder cups #1-#5, as well as cup #9 generously with solder
– Optional: Cut all leads down to a smaller size – this is a judgement call.
– Shorter wires leave less margin for error if a wire breaks. Longer wires are harder to pack into the case without obstructing the joystick motion.
– Solder the following connections into the DB-9 connector:

○ cup #1 – Either joystick electrode wire
○ cup #2 – Hall sensor GND wire

▪ The GND wire can be identified using a multimeter connectivity tester. The GND wire will be electrically connected to pin 8 of the Hall sensor IC, and the corner-facing terminal of the C1 capacitor
○ cup #3 – Hall sensor GND wire

▪ The 5V wire will be electrically connected to pins 2, 3, and 7 of the Hall sensor IC, as well as the non-corner-facing terminal of the C1 capacitor
○ cup #4 – Hall sensor X position wire

▪ Either Hall sensor wire that is **not** GND or 5V can be connected here (X and Y position signals can be easily swapped in software later)
○ cup #5 – Hall sensor Y position wire

▪ The remaining Hall sensor wire
○ cup #9 – The remaining joystick electrode wire
– Apply dental acrylic to faces of solder cups - cover all the solder, and a little on the wire insulation
– **Make sure that the dental acrylic doesn’t form a protrusion large enough that when the DB-9 is installed in the case, it will interfere with the motion of the gimbal**.
– Cure dental acrylic with the UV light
– See Figure 24 and Figure 25 for the results of this step

**Figure 24.**
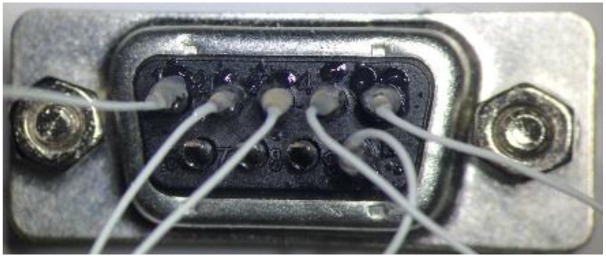
Completed DB-9 connector with all six wires soldered in and covered in acrylic

**Figure 25.**
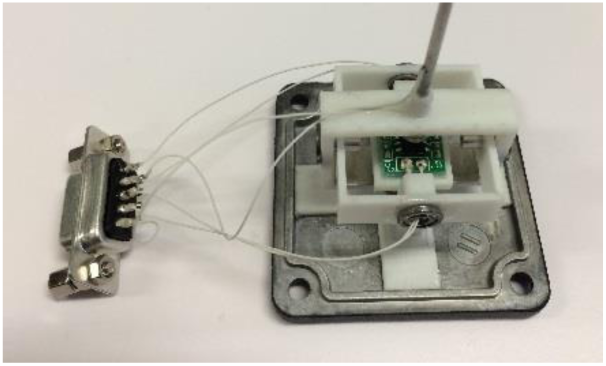
Joystick connected to DB-9

Encase joystick
– Insert the 1/4-20 × 1” mounting bolt into the inside of the side hole in the case and thread it all the way through to snug. It should go in the right-hand side of the case when the DB-9 hole is facing you and the joystick hole is on top.
– Carefully thread the six wires through the slot in the DB-9 hole of the case.
– Fasten the two DB-9 screws firmly into the casing. The inner nuts on each screw should allow the DB-9 to stand off a bit from the case, which helps prevent the joystick from bumping into the wires sticking out of the solder cups.
– Repeat for each base plate corner hole:

○ Insert a rubber grommet on one of the base plate corner holes between the base plate and the top of the housing
○ Insert a screw into the same hole from below the base plate.
○ **Make sure there are no wires caught in or around the grommet** – tightening the screw down on a wire will likely damage or sever the wire.
○ Tighten the screw **only** until it is just secure – do not overtighten!
– Use tweezers to gently push all exposed wire into the casing
– **Make sure joystick wires don't impede joystick's free motion**

○ Test joystick by gently displacing it then releasing it.
○ The joystick’s motion should be very bouncy (each bounce should last roughly 2-4 seconds before it settles into equilibrium)
○ Note that the range of motion parallel to the rectangle piece will naturally be smaller than in the other dimension because of the geometry of the case. The joystick motion will be limited by the hole in the bottom of the cage, so this restriction is not relevant and doesn’t need to be corrected.
○ If the joystick’s motion seems damped or restricted, it is likely because some of the wires are pressing against the gimbal.

▪ Use tweezers to gently rearrange the wires until they are no longer restricting the motion of the gimbal.
▪ If the wires become a persistent problem for joystick motion, you can also open the case back up and use some superglue to fix the middle of the wires to the baseplate to control their positioning. Take care to leave enough slack so the gimbal can move freely.
– See Figure 26 and Figure 27 for the results of this step.

**Figure 26.**
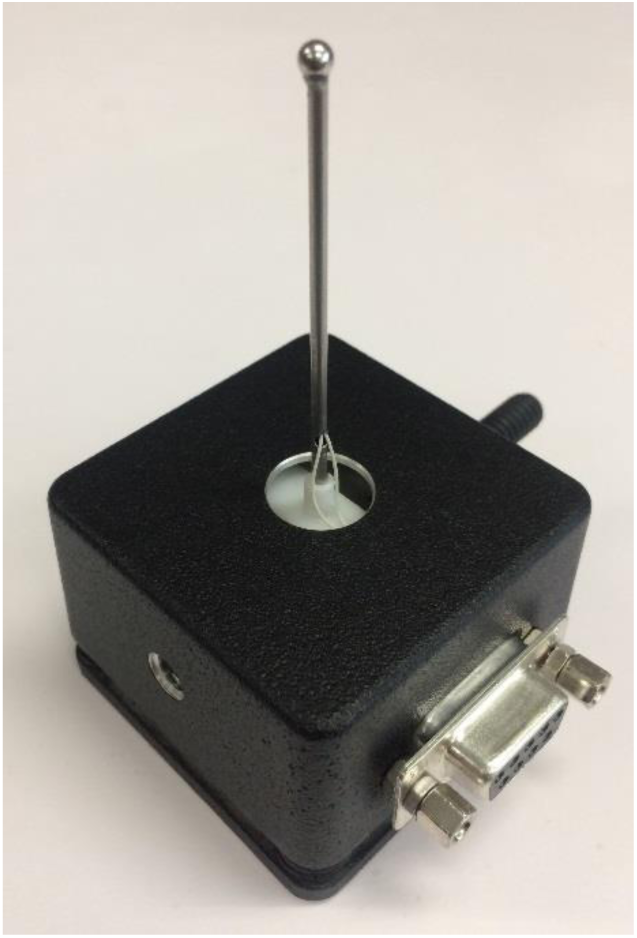
Finished joystick

**Figure 27.**
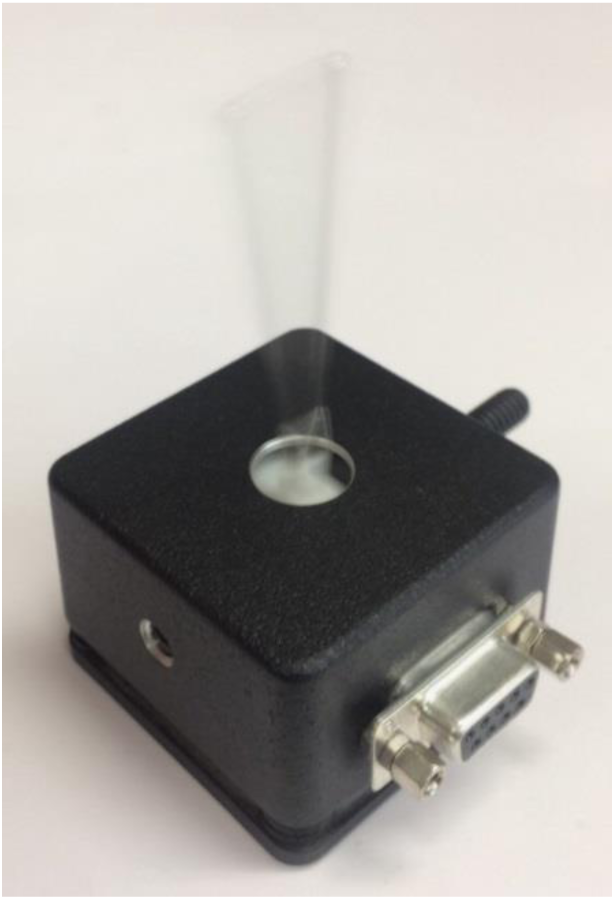
Finished joystick exhibiting a good bounce

#### Part II: Mount joystick on micromanipulator

##### Full list of Materials

| # | Part Description | Manufacturer/<br>Supplier | Manufacturer/Supplier Part Number | Datasheet/Drawing/<br>Modification |
| --- | --- | --- | --- | --- |
| 1 | Completed joystick |  |  |  |
| 2 | 1/2" XY Dovetail Translation Stage with baseplate | Thor Labs | DT12XY | https://www.thorlabs.com/thorproduct.cfm?partnumber=DT12XY |
| 3 | 1/4"-20 M6 countersunk cap screw | Thor Labs | (included with DB12XY) |  |
| 4 | Hex head bolt, washer, and locking washer | Thor Labs | (included with DB12XY) |  |
| 5 | RA bracket | N.A. (Custom Made) | N.A. (Custom Made) | AttachJSMicroManipulator from here |
| 6 | 8-32 x 1/4" bolt |  |  |  |
| 7 | 1/4" x 20 nut x2 |  |  |  |

##### Building Instructions

###### General notes

The micromanipulator provides a mounting connection between the joystick and the homecage, and provides the ability to precisely center the joystick in the cage-bottom hole, which permits accurate calibration and a uniform maximum joystick displacement in all directions.

###### 1. Connect the two DT12 translation stages to each other

###### 2. Attach the mounting adaptor to the translation stage

– Insert the hex head bolts, washers, and locking washers that come with the mounting adaptor into the two holes on either side of the center hole in the DT12XY baseplate. The locking washer should fit neatly into the hole. Do not tighten all the way.
– Insert the 1/4”-20 M6 countersunk cap screw into the center hole in the DT12XY baseplate. The threaded end will eventually be pointing upwards away from the micromanipulator.
– Slide the baseplate onto one of the two translation stages and center it
– Tighten the hex head bolts until the baseplate is secure

**Figure 28.**
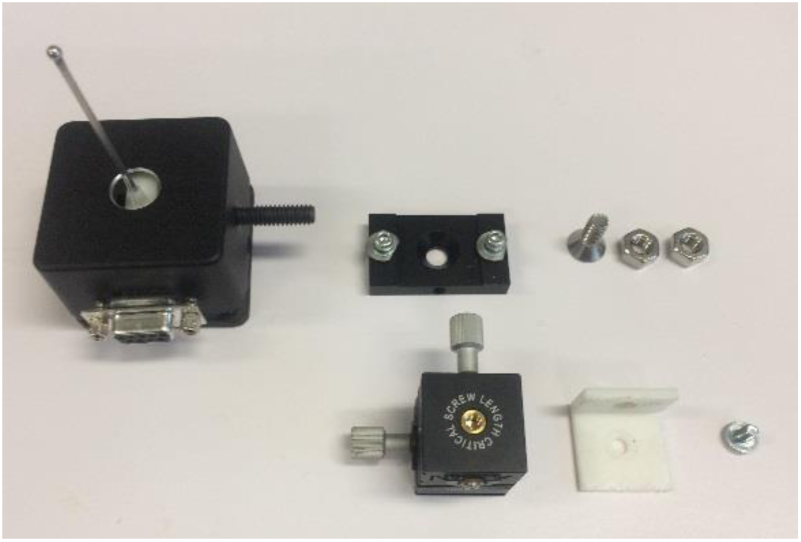
Materials for mounting joystick on micromanipulator. Clockwise from to pleft: Completed joystick, DT12B mounting adaptor, 1/4”x20 M6 countersunk cap screw, 1/4”×20 nut x2, 8-32×1/4” bolt, RA adapter, DT12 translation stages x2 (already attached)

**Figure 29.**
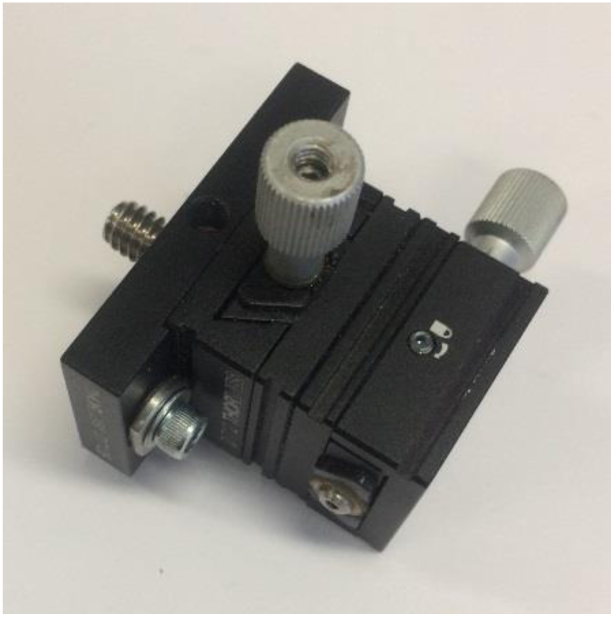
Assembled micromanipulator

###### 3. Mount the joystick on the micromanipulator

– Attach the large side of the RA bracket to the translation stage that does NOT have the DT12XY baseplate attached using the 8-32×1/4” bolt. This will be the bottom side of the micromanipulator. Tighten with a screwdriver.
– Insert the 1/4”-20×1” mounting bolt of the joystick housing into the other (smaller) side of the RA bracket.
– Thread on the other 1/4”-20 nut and tighten with a wrench.
– Ensure that the joystick housing sides are closely aligned with the RA bracket sides, so the joystick will be vertical when mounted.
– Visually check that the joystick is parallel to the micromanipulator top screw. If it isn’t, small adjustments can be made to the joystick housing screws, and the connection between the micromanipulator and the joystick to align it.
– See Figure 30 for completed mounting of joystick on micromanipulator

**Figure 30.**
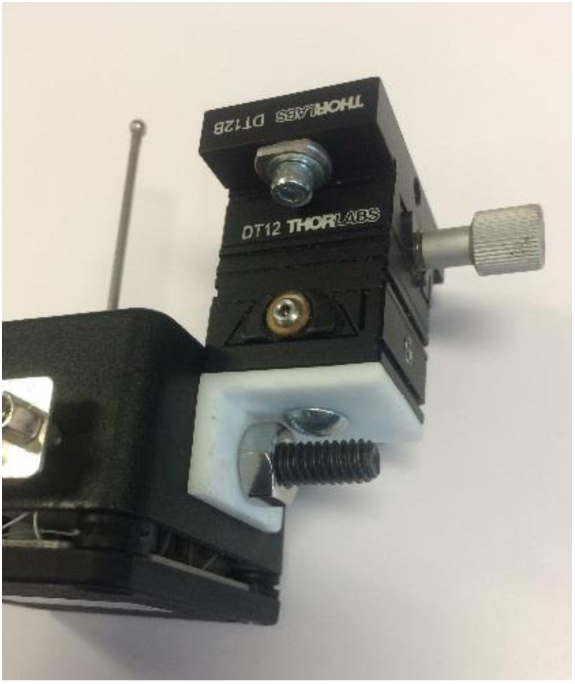
Joystick mounted on micromanipulator

#### Part III: Building a joystick board

##### Full list of Materials

| # | Part Description | Manufacturer/<br>Supplier | Manufacturer/Supplier<br>Part Number | Datasheet/Drawing/Modification |
| --- | --- | --- | --- | --- |
| 1 | Joystick Breakout PCB | N.A. (Custom Made) | N.A. (Custom Made) | Schematic and PCB here |
| 2 | Capacitor, 10 pF, SMT x2 | Samsung/Digikey | 1276-2154-1-ND | https://www.digikey.com/product-detail/en/samsung-electro-mechanics/CL10C100JB8NCNC/1276-2154-1-ND/3890240 |
| 3 | Capacitor, 1 nF, 0603 case x3 | Samsung/Digikey | 1276-1091-1-ND | https://www.digikey.com/product-detail/en/samsung-electro-mechanics/CL10C102JB8NNNC/1276-1091-1-ND/3889177 |
| 4 | Capacitor, 10 nF, SMT x2 | Samsung/Digikey | 1276-1103-1-ND | https://www.digikey.com/product-detail/en/samsung-electro-mechanics/CL10B103JB8NNNC/1276-1103-1-ND/3889189 |
| 5 | Capacitor, 0.1 μF (100 nF), 0603 case | Samsung/Digikey | 1276-6998-1-ND | https://www.digikey.com/product-detail/en/samsung-electro-mechanics/CL10B104KO8ZW6C/1276-6998-1-ND/7320640 |
| 6 | Capacitor, 0.22 μF, radial leads | Samsung/Digikey | 1276-1996-1-ND | https://www.digikey.com/product-detail/en/samsung-electro-mechanics/CL10B223JB8NNNC/1276-1996-1-ND/3890082 |
| 7 | Capacitor, 0.47 μF, 0402 case x2 | Samsung/Digikey | 1276-2082-1-ND | https://www.digikey.com/en/supplier-centers/s/samsung-electro-mechanics-america |
| 8 | Resistor, 160 $\Omega$, 0402 case | Vishay-Dale/Digikey | 541-160YCT | https://www.digikey.com/product-detail/en/vishay-dale/CRCW0402160RFKEDHP/541-160YCT-ND/2825954 |
| 9 | Resistor, 1 kΩ, 0402 case x5 | Panasonic/Digikey | P1.00KLCT | https://www.digikey.com/product-detail/en/panasonic-electronic-components/ERJ-2RKF1001X/P1.00KLCT-ND/97341 |
| 10 | Resistor, 2.6 kΩ, 0402 case | Panasonic/Digikey | P2.61KLCT | https://www.digikey.com/product-detail/en/panasonic-electronic-components/ERJ-2RKF2611X/P2.61KLCT-ND/194243 |
| 11 | Resistor, 3.3 kΩ, 0402 case x2 | Rohm/Digikey | RHM3.3KCDCT | http://www.digikey.com/product-detail/en/MCR01MRTF3301/RHM3.3KCDCT-ND/4083740 |
| 12 | Resistor, 9 kΩ, 0402 case | Panasonic/Digikey | P9.10KLCT | https://www.digikey.com/product-detail/en/panasonic-electronic-components/ERJ-2RKF9101X/P9.10KLCT-ND/17467224 |
| 13 | Resistor, 22 kΩ, 0402 case | Panasonic/Digikey | P22KJCT | http://www.digikey.com/product-detail/en/ERJ-2GEJ223X/P22KJCT-ND/146939 |
| 14 | Voltage comparator IC, SMT | TI/Digikey | TL331IDBVR | http://www.ti.com/lit/ds/symlink/tl331.pdf |
| 15 | Op Amp IC, SMT | TI/Digikey | LM358 | https://www.digikey.com/product-detail/en/texas-instruments/LM358DR/296-1014-1-ND/404838 |
| 16 | Voltage regulator | STMicroelectronics/Digikey | L7805CV | https://www.digikey.com/product-detail/en/stmicroelectronics/L7805CV/497-1443-5-ND/585964 |
| 17 | Touch sensor IC, SMT x2 | Atmel/Digikey | AT42QT1011 | http://www.atmel.com/images/Atmel-9542-AT42-QTouch-BSW-AT42QT1011_Datasheet.pdf |
| 18 | Transistor x2 | Micro Commercial Co/Digikey | 2N3904 | https://www.digikey.com/product-detail/en/micro-commercial-co/2N3904-AP/2N3904-APCT-ND/950591?utm_adgroup=Semiconductor%20Modules&gclid=EAlaI_QobChMlpO3hideR2gIVFFcNCh0mPwH6EAAYASAAEgIVBvD_BwE |
| 19 | 2-pin header x4 | Hirose Electric/Digikey | DF3A-2P-2DSA | http://www.digikey.com/product-detail/en/DF3A-2P-2DSA/H2094-ND/141512 |
| 20 | 3-pin header x2 | Hirose Electric/Digikey | DF3A-3P-2DSA | http://www.digikey.com/product-detail/en/DF3A-3P-2DSA/H2095-ND/141515 |
| 21 | 4-pin header | Hirose Electric/Digikey | DF3A-4P-2DSA | http://www.digikey.com/product-detail/en/DF3A-4P-2DSA/H2096-ND/141518 |
| 22 | DB-9 connector, male, through hole | Amphenol/Digikey | A34064-ND | https://www.digikey.com/product-detail/en/te-connectivity-amp-connectors/5747871-2/A34064-ND/1123840 |
| 23 | DB-15 connector, female, RA through hole | Amphenol/Digikey |  | https://www.digikey.com/product-detail/en/amphenol-commercial-products/17EBH-015-S-AM-0-10/17EBH-015-S-AM-0-10-ND/1242549 |
| 24 | Heat sink | Aavid | HS115-ND | https://www.digikey.com/product-detail/en/507302B00000G/HS115-ND/5849 |
| 25 | Heat sink compound, Silicone (type Z9) | GC Electronics | 10-8109 | [http://www.gcelectronics.com/order/DataSheets/10-8109%2010-8108%2010-8107-10%2010-8107-100%2010-8106%20Silicone%20(Z9).pdf](http://www.gcelectronics.com/order/DataSheets/10-8109%2010-8108%2010-8107-10%2010-8107-100%2010-8106%20Silicone%20%28Z9%29.pdf) |

##### Building Instructions

###### General notes

The joystick board collects and conditions the signals from the various sensors, including the joystick, and passes the resulting signals to the sb-RIO. The schematic contains the locations of where the components have to be soldered. If soldering, especially surface mount soldering, is new to you, there are many good guides available on the internet.

###### 1. Soldering the SMT components

###### 2. Soldering the through-hole components

Part IV: Building the lick port with lick sensor and solenoid valve

##### Full list of Materials

| # | Part Description | Manufacturer /Supplier | Manufacturer/Supplier Part Number | Datasheet/Drawing/Modification |
| --- | --- | --- | --- | --- |
| 1 | Lick spout, Coulbourn Instruments | Coulbourn Instruments | H24-01-TB-01 | http://www.coulbourn.com/v/vspfiles/assets/manuals/habitesth-serieshardwareuserguide.pdf |
| 2 | 1/16" ID food-grade plastic tubing | US Plastics | 56400 | http://www.usplastic.com/catalog/item.aspx?sku=56400&gclid=EAlaIqobChMlIcKbn7_m1gIVCYVpCh3DSwUWEAQYASABEglvWvD_BwE |
| 3 | 1/16" barbed plug | VWR | P0-1NK | https://us.vwr.com/store/product/14459897/barbed-plugs-eldon-james |
| 4 | Solenoid valve | The Lee Company | LHDA2433215H | http://www.theleeco.com/electro-fluidic-systems/solenoid-valves/lhd/3-port-ported-style.cfm |
| 5 | Photointerrupter switch | Lite-On Inc./Digikey | LTH-301-32 | https://www.digikey.com/product-detail/en/lite-on-inc/LTH-301-32/160-1936-ND/3198349 |
| 6 | 3-pin receptacle | Hirose/Digikey | DF3-3S-2C | http://www.digikey.com/product-detail/en/DF3-3S-2C/H2084-ND/141482 |
| 7 | 4-pin receptacle | Hirose/Digikey | DF3-4S-2C | https://www.digikey.com/product-detail/en/DF3-4S-2C/H2085-ND/141485 |
| 8 | Crimp terminals | Hirose/Digikey | DF3-2428SC | https://www.digikey.com/product-detail/en/DF3-2428SC/H1500-ND/141644 |
| 9 | Stranded, insulated wire, approximately 26 AWG |  |  |  |
| 10 | Heatshrink tubing |  |  |  |

##### Building Instructions

###### General notes

The lick port consists of a lick spout, a solenoid valve, and a lick sensor, and associated cables. The lick sensor and the solenoid are controlled via the joystick PCB. The procedure for the lick sensor is very similar to the nosepoke sensor. Note that while this research used a beam-break lick sensor, we have subsequently transitioned to using a capacitive lick sensor.

###### 1. Extract the transmitter & receiver from the plastic photointerrupt switch housing

– Use a cutting tool (for example a Dremel with a cutting wheel) to carefully remove the transmitter and receiver from the plastic housing. Take care not to damage the transmitter and receiver.
– Paint the back side of both the transmitter and receiver black to block spurious light signals from the environment.
– See Figure 32 for the results of this step.

**Figure 31.**
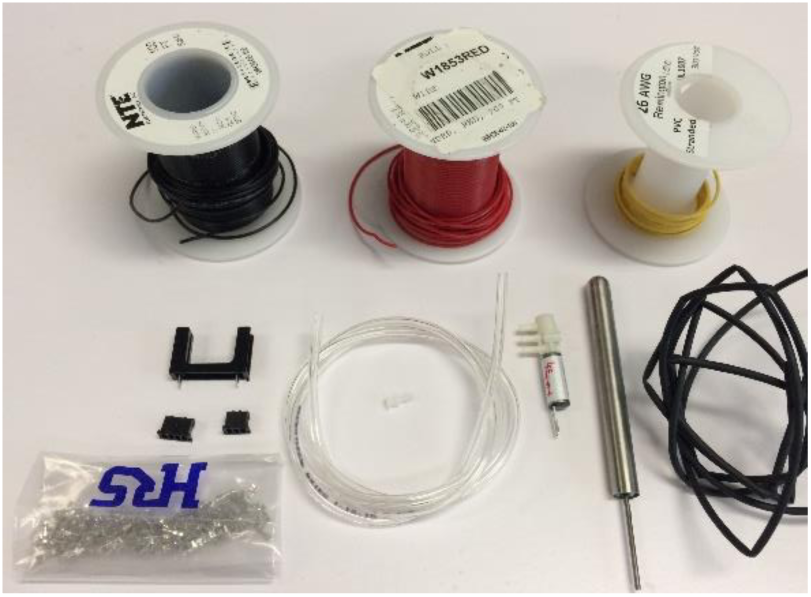
Materials for lick port/sensor/valve. Clockwise from top left: Wire, heatshrink tubing, water spout, solenoid valve, 1/16” tubing, 1/16” plug, crimp terminals, 3-pin and 4-pin receptacles, phototransistor interrupt switch

###### 2. Cut and strip wire leads

– Cut two ground wires, and two transmit/receive wires, approximately 8 inches each. Color coding is recommended.
– Use a wire stripper to strip the ends of the wires.
– See Figure 34 for the results of this step.

**Figure 32.**
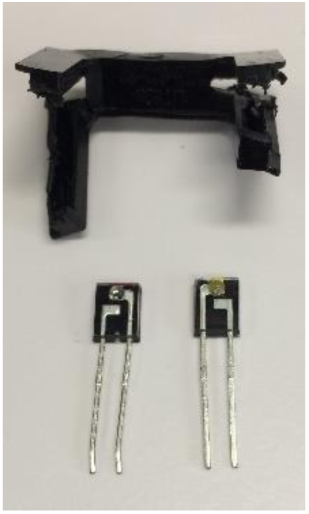
Transmitter (left) and receiver (right) extracted from their housing (top) and coated

###### 3. Solder wires to transmitter and receiver

– The receiver needs an “LickOut” wire (lick signal) and a ground wire, and the transmitter needs a 5V wire and a ground wire.
– See Figure 33 to identify terminals of the transmitter and receiver.
– Solder the ground wire, LickOut, and 5V wires onto the corresponding terminals.
– Cover bare metal connections with heatshrink tubing.
– See Figure 37 for the results of this step

**Figure 33.**
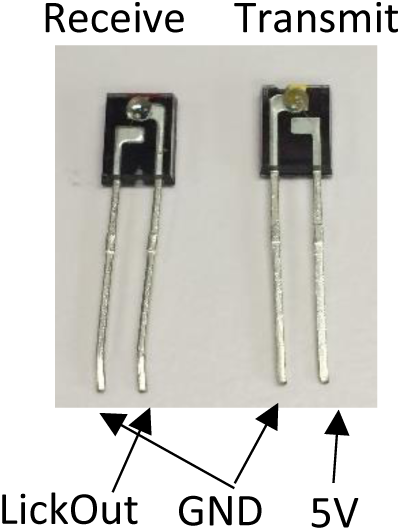
Identifying the terminals of the lick sensor transmitter and receiver

**Figure 34.**
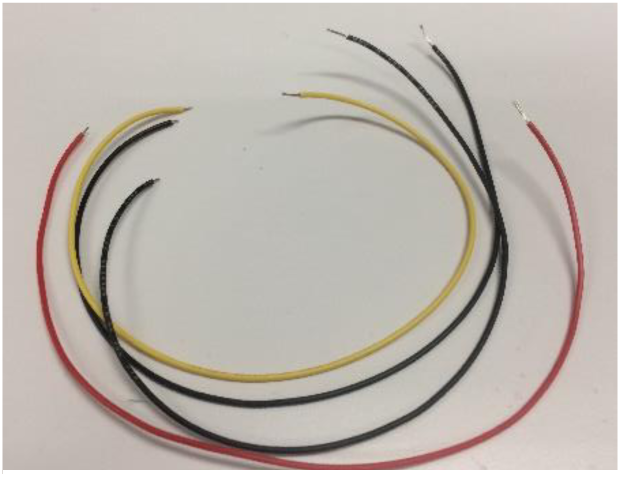
Cut and stripped wires for the lick sensor

###### 4. Add receptacle

– Add crimp terminals to the three stripped ends of the lick sensor wires
– Optional: braid wires
– Insert the crimped ends of the wires into the 3-pin receptacle in the order shown in Figure 35

###### 5. Glue lick sensor to lick spout

– Use superglue to affix the transmitter and receiver on either side of the lick spout so the beam from the transmitter’s lens passes right in front of the lick spout hole and then into the receiver’s lens.
– Make sure that the wires stick close together and close to the sides of the lick spout for about 2 inches, or the spout won’t fit in the custom lick spout holder later.
– Line up the lenses so the beam will pass the lick spout hole at a distance of approximately 1 mm.
– See Figure 38 for the results of this step.

**Figure 35.**
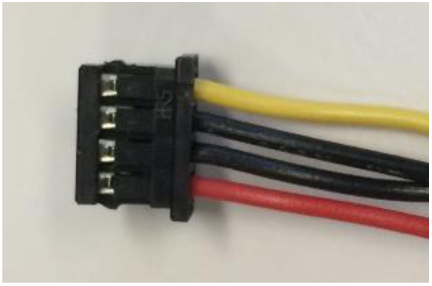
Lick sensor wire receptacle, showing connection order. Red = 5V, Black = Ground, Yellow = signal out

###### 6. Add wires to solenoid valve

– Cut two wires to a length of approximately 8” each.
– Strip both ends of the wires.
– Solder the wires onto the solenoid valve. The solenoid valve is nonpolar, so the order of the wires is not important.
– Add heatshrink tubing to the bare wire joints for insulation and strength.
– Add crimp terminals to the other ends of the wire, and insert into the 3-pin receptacle as shown in Figure 39.

**Figure 36.**
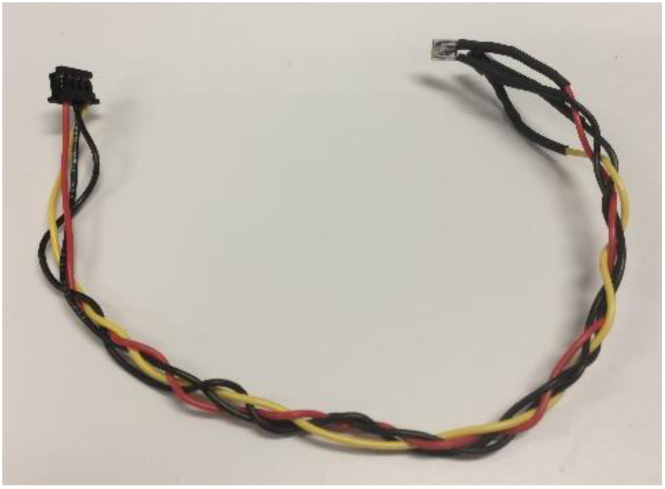
Assembled lick sensor

**Figure 37.**
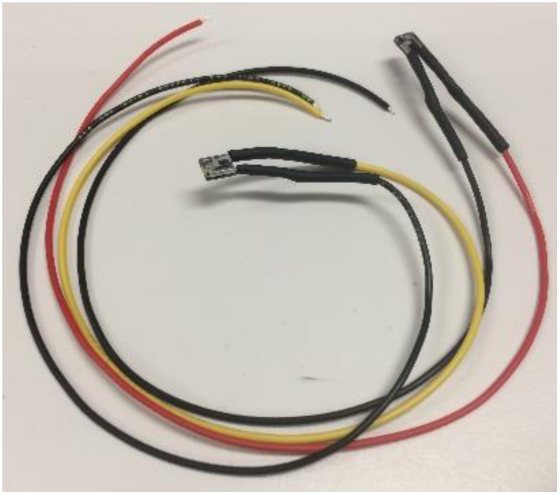
Transmitter and receiver with wires and heatshrink tubing

**Figure 38.**
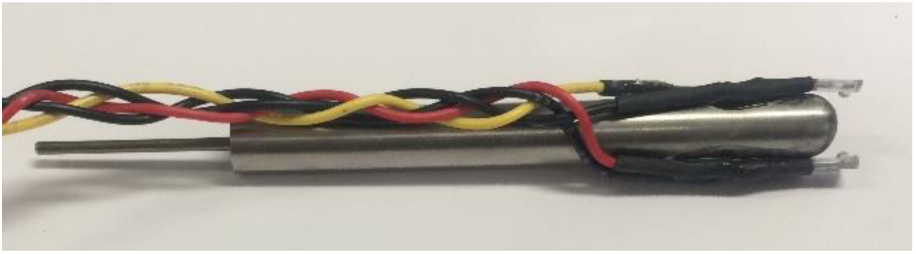
Lick sensor attached to water spout

**Figure 39.**
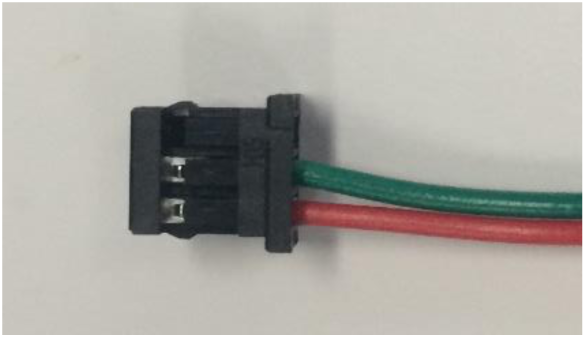
Solenoid valve wires inserted into receptacle. Only the bottom and middle spots should be filled, although the order doesn't matter.

###### 7. Add tubes to solenoid valve

– Cut an approximately 1” section of 1/16” tubing.
– Push the 1” piece of tubing onto the valve outlet closest to the wire terminals, then add a 1/16” plug to seal that outlet
– Cut two more approximately 12” lengths of tubing
– Push the two 12” lengths of tubing onto the other two valve outlets.
– See Figure 41 for the results of this step.

**Figure 40.**
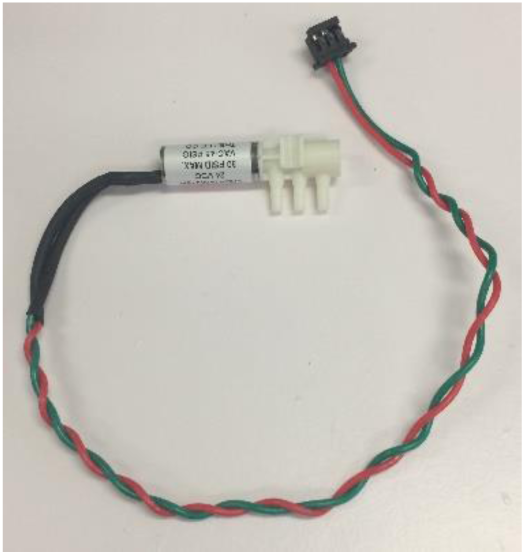
Solenoid valve with wires connected

###### 8. Connect the valve to the lick spout

– Push the free end of one of the 12” sections of tubing from the valve onto the lick spout tube that extends from the bottom. Push the tubing far enough on to make a good seal.
– See Figure 42 for the completed assembly of the solenoid valve, lick spout, and lick sensor.

**Figure 41.**
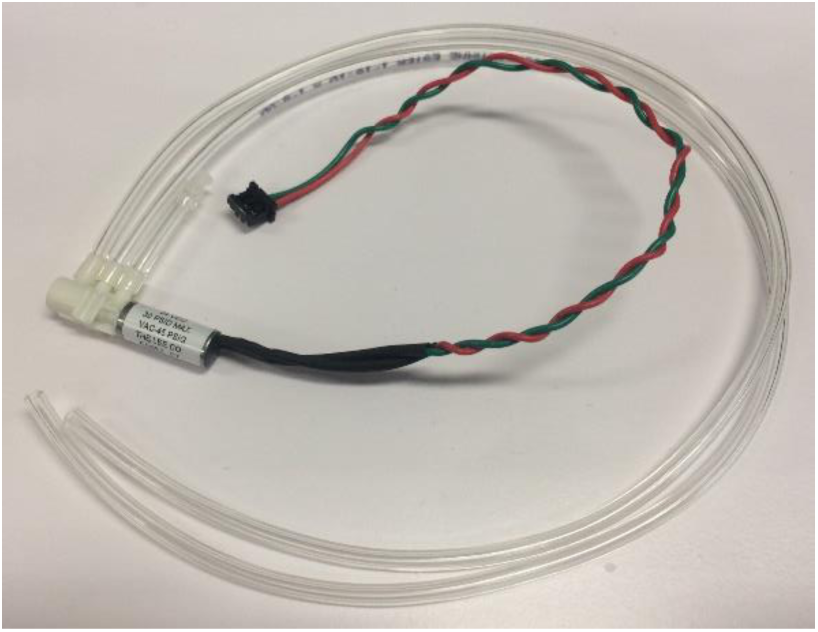
Solenoid valve with wires and tubes connected

**Figure 42.**
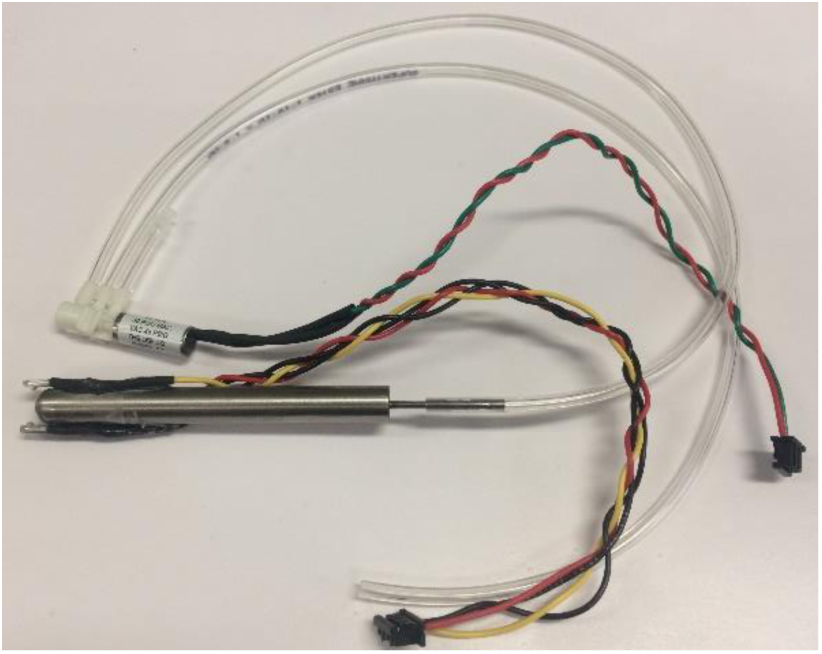
Completed solenoid valve/water spout/lick sensor assembly

#### Part V: Building a fixed post touch sensor

##### Full list of Materials

| # | Part Description | Manufacturer /Supplier | Manufacturer/Supplier Part Number | Datasheet/Drawing/Modification |
| --- | --- | --- | --- | --- |
| 1 | Machined ball for touch sensor electrode | N.A. (Custom Made) | N.A. (Custom Made) | N.A. (Custom Made) |
| 2 | Stainless Steel 304 Hypodermic Tubing, 15 Gauge, 0.072" OD, 0.0625" ID, 0.00475" Wall, 36" Length | Amazon |  | https://www.amazon.com/gp/product/B004WPQJHC |
| 3 | Machined shafts for bearings (Stainless Steel 316L Seamless Round Tubing, 1/8" OD, 0.069" ID, 0.028" Wall, 36" Length) | Amazon |  | https://www.amazon.com/gp/product/B004XN9LLE |
| 4 | Ultra-Flexible PVC Lead Wire, 38 AWG | Cooner Wire Company | NUF38-1650T-WHITE | https://www.coonerwire.com/pvc-hookup-wire-nuf-series/ |
| 5 | Fixed post holder | N.A. (Custom Made) | N.A. (Custom Made) |  |
| 6 | Dental Acrylic |  | Flow-It ALC Flowable Composite, P/N: N11H |  |
| 7 | Coltolux LED UV curing light | Coltene | C7970100115 | https://nam.coltene.com/products/restoration/curing-lights/coltolux-led/coltoluxR-led/ |
| 8 | Superglue (Loctite 401) | Loctite | 401 | http://www.loctite.sg/sea/content_data/93806_NEWCA401EN.pdf |
| 9 | 5-minute epoxy (HARDMAN® Extra-Fast Setting Epoxy) | Hardman | 04001 | http://www.royaladhesives.com/Files/Hardman-Structural/DOUBLE-BUBBLE_04001_RED_TDS.PDF |
| 10 | Silver conductive paint | GC Electronics | 22-023 | http://www.gcelectronics.com/order/DataSheets/22-023,%2022-024%20Silver%20Print%20II%20Conductive%20Paint.pdf |
| 11 | 2-pin receptacle | Hirose/Digikey | DF3-2S-2C | https://www.digikey.com/product-detail/en/DF3-2S-2C/H2083-ND/141479 |
| 12 | Crimp terminals | Hirose/Digikey | DF3-2428SC | https://www.digikey.com/product-detail/en/DF3-2428SC/H1500-ND/141644 |
| 13 | Stranded insulated wire, approximately 26 AWG |  |  |  |
| 14 | Heatshrink tubing, small (optional) |  |  |  |

##### Building Instructions

###### General notes

The procedure for creating the fixed touch post is almost the same as creating the joystick electrode. Therefore, rather than repeat the instructions, refer to the section “2. Making the Joystick Electrode” above, and to the differences listed below.

###### 1. Fabricating the fixed post tube

– The length of the fixed post tube should be at least 1 inch, but the exact length is not important.
– The tube does NOT need a side-hole. The leads can come directly out the bottom of the tube.

###### 2. The fixed-post holder

– Since the tube does not sit on a gimbal as in the case of the joystick electrode, the post needs some support and stabilization. This is provided by a custom 3D printed fixed post holder.
– Slide the post holder onto the tube. Do NOT glue it on yet.

###### 3. The connector

– Rather than a DB-9 connector, the fixed post connects to the joystick board via a twisted pair of wires that lead to a 2-pin receptacle, which plugs into a header on the JS board.
– Carefully strip and solder the free ends of the Cooner wires from the fixed post onto two 26 AWG wires twisted into a pair, approximately 8” long.
– Add heatshrink tubing or dental acrylic to reinforce the connections between the Cooner wires and the 26 AWG wires.
– Apply crimp terminals to the other ends of the 26 AWG wires, and insert them into the 2-pin receptacle. The order doesn’t matter.
– See Figure 43 for the completed fixed post touch sensor

**Figure 43.**
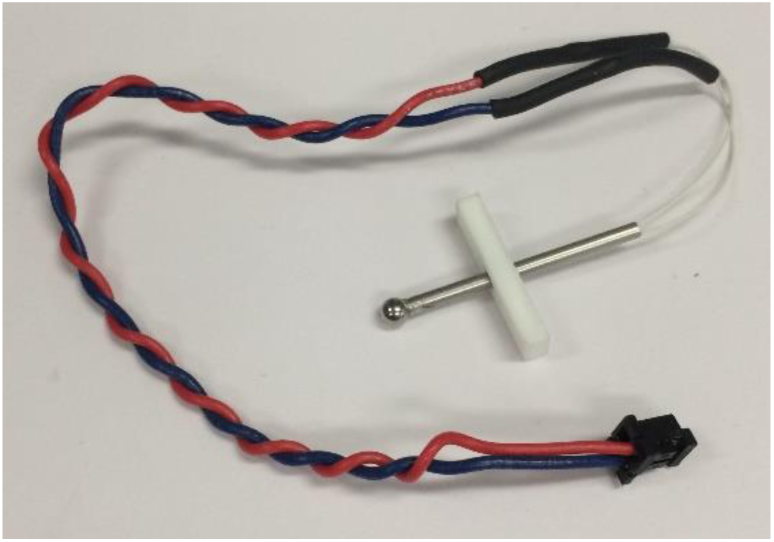
Completed fixed post touch sensor

#### Part VI: Building the nosepoke sensor

##### Full list of Materials

| # | Part Description | Manufacturer /Supplier | Manufacturer/Supplier Part Number | Datasheet/Drawing/Modification |
| --- | --- | --- | --- | --- |
| 1 | Photointerrupter switch | Lite-On Inc./Digikey | LTH-301-32 | https://www.digikey.com/product-detail/en/lite-on-inc/LTH-301-32/160-1936-ND/3198349 |
| 2 | Resistor, 200Ω, axial lead | Stackpole Electronics/Digikey | CF14JT200RCT-ND | https://www.digikey.com/product-detail/en/stackpole-electronics-inc/CF14JT200R/CF14JT200RCT-ND/1830333 |
| 3 | 3-pin receptacle | Hirose/Digikey | DF3-3S-2C | http://www.digikey.com/product-detail/en/DF3-3S-2C/H2084-ND/141482 |
| 4 | Crimp terminals | Hirose/Digikey | DF3-2428SC | https://www.digikey.com/product-detail/en/DF3-2428SC/H1500-ND/141644 |
| 5 | Stranded, insulated wire, approximately 26 AWG |  |  |  |
| 6 | Heatshrink tubing |  |  |  |

##### Building Instructions

###### General notes

The nosepoke sensor detects when the mouse pokes its nose through the hole in the cage that leads to the lick spout. It operates by shining an infrared (IR) beam from a transmitter LED on one side of the hole to a receiver phototransistor on the other side.

When the mouse pokes its nose in, the IR beam is interrupted.

Note that circuitry on the joystick PCB is necessary for biasing and interpreting the transmitter and receiver.

###### 1. Detach the transmitter/receiver from each other

– Cut away the plastic bridge connecting the transmitter/receiver

###### 2. Cut and strip wire leads

– Cut three ground wires (~4”), a transmit/5V wire (~7”), and a receive wire (~8”). Color coding is recommended.
– Use a wire stripper to strip the ends of the wires.
– See Figure 45 for the results of this step.

**Figure 44.**
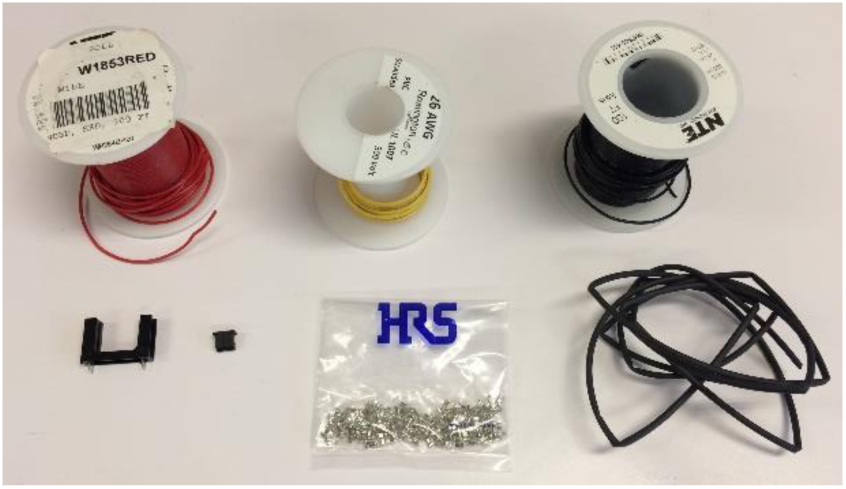
Nosepokr sensor materials. Clockwise from top lest: three spools of 26 AWG stranded wire, heatshirnk wrap, crimp terminals, 3-pin receptacle, IR transmitter & receiver. Not pictured: 200 Ω resistor.

**Figure 45.**
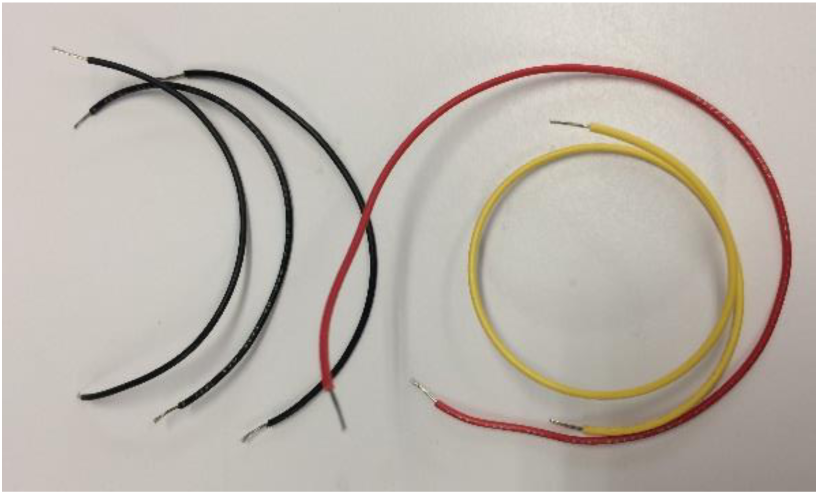
Cut and stripped wires for the nosepoke sensor

###### 3. Add wires and resistor to transmitter and receiver

– The receiver needs an “NPout” wire (nosepoke signal) and a ground wire, and the transmitter needs a 5V wire and a ground wire. The transmitter also requires a 200 Ω current limiting resistor. We will incorporate this resistor as part of the transmitter’s power wire.
– See Figure 46 to identify terminals of the transmitter and receiver.
– Solder a ground wire onto each ground terminal
– Solder the NPout and 5V wires onto the corresponding terminals
– Cut and strip a break in the 5V wire, then solder in the 200 Ω resistor to close the gap.
– See Figure 50 for the results of this step

###### 4. Solder wires and resistor onto transmitter and receiver

– Solder the two ground wires together along with a common ground wire.
– Cover bare metal connections with heatshrink wrap.
– See Figure 47 for the results of this step.

###### 5. Add receptacle

– Add crimp terminals to the three stripped ends of the nosepoke sensor wires
– Insert the crimped ends of the wires into the 3-pin receptacle in the order shown in Figure 48

**Figure 46.**
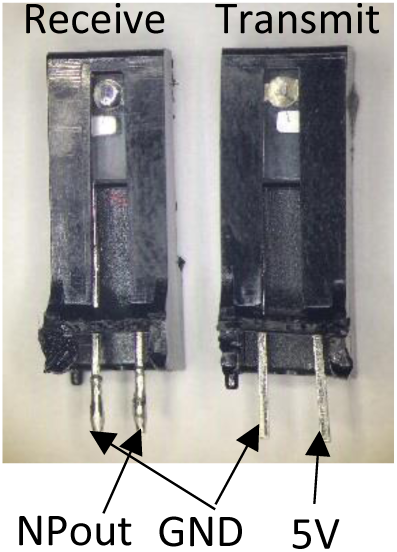
Identifying the terminals of the transmitter and receiver

**Figure 47.**
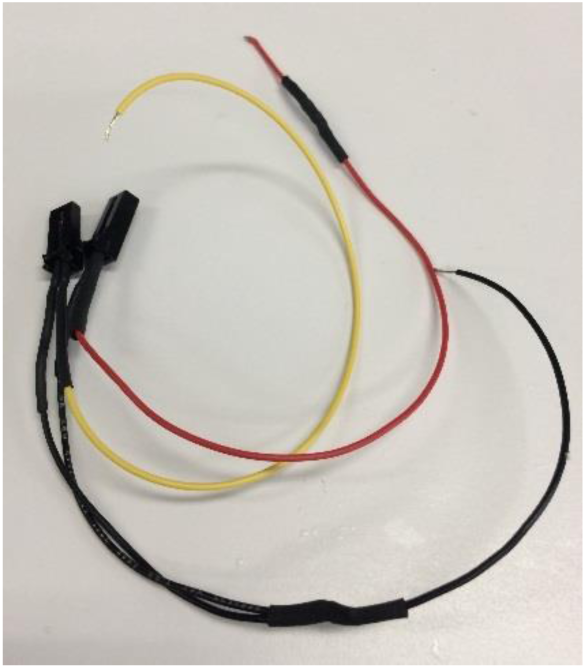
Ground wires soldered together, bare metal covered in heatshrink wrap

**Figure 48.**
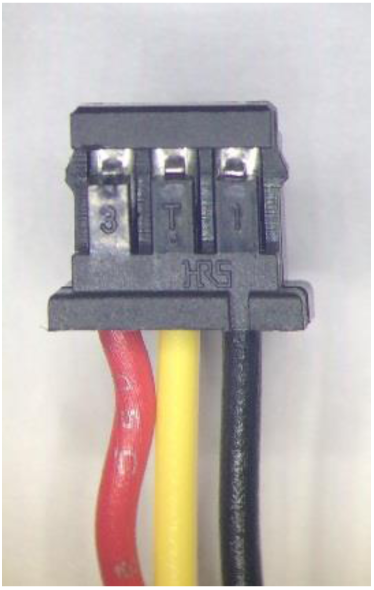
Nosepoke wire receptacle, showing connection order. Red = 5V, Yellow = signal out, Black = Ground

**Figure 49.**
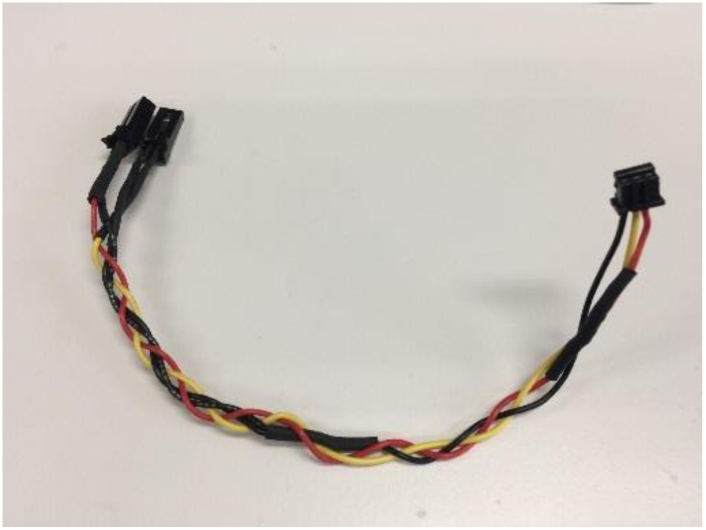
Completed nosepoke sensor

**Figure 50.**
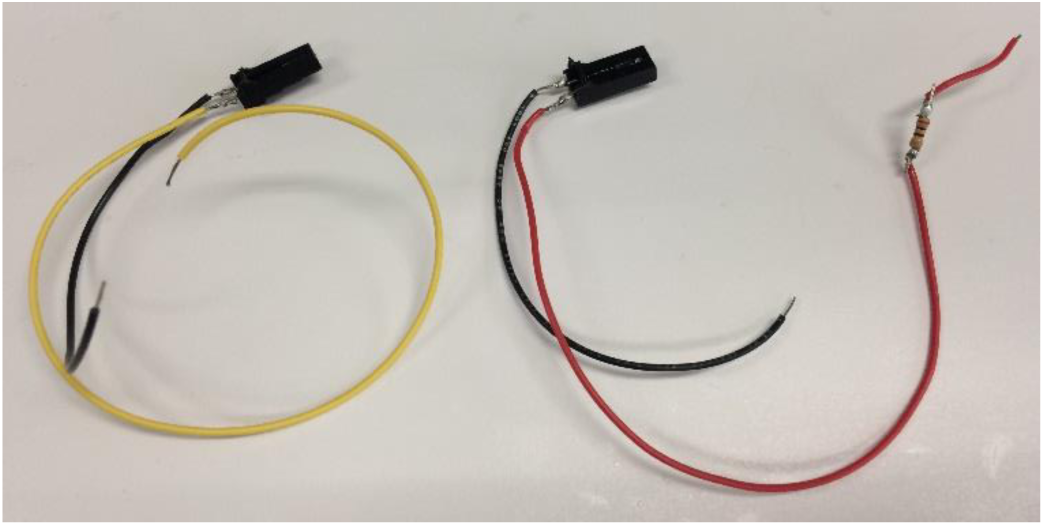
Nosepoke sensor with wires and resistor soldered

#### Part VII: Building the masking light

##### Full list of Materials

| # | Part Description | Manufacturer /Supplier | Manufacturer/Supplier Part Number | Datasheet/Drawing/Modification |
| --- | --- | --- | --- | --- |
| 1 | LED light strip |  |  |  |
| 2 | 2-pin receptacle | Hirose/Digikey | DF3-2S-2C | https://www.digikey.com/product-detail/en/DF3-2S-2C/H2083-ND/141479 |
| 3 | Crimp terminals | Hirose/Digikey | DF3-2428SC | https://www.digikey.com/product-detail/en/DF3-2428SC/H1500-ND/141644 |
| 4 | 5-minute epoxy (HARDMAN® Extra-Fast Setting Epoxy) | Hardman | 04001 | http://www.royaladhesives.com/Files/Hardman-Structural/DOUBLE-BUBBLE_04001_RED_TDS.PDF |
| 5 | Stranded, insulated wire, approximately 26 AWG |  |  |  |
| 6 | Heatshrink tubing |  |  |  |

##### Building Instructions

###### General notes

The masking light is an LED with wires and a 2-pin connector. It plugs into the joystick board and will be attached to the homecage above the lick port. Its purpose is to prevent mice from visually discriminating between trials when the optogenetic laser is on or off due to laser light leaking from the cannula.

###### 1. Cut one LED unit off of the strip

– Cut one of the LED units off of the strip along the printed divider lines.
– Peel and cut away the plastic coating on the edge of the LED unit that covers the copper terminals marked “+” and “-”
– See Figure 53 for the results of this step.

###### 2. Solder wires onto the LED

– Cut and strip two wires approximately 8” long
– Solder one end of each wire onto the exposed terminals on the LED unit marked “+” and “-”. Color coding is recommended.
– Coat the exposed junction with epoxy or dental acrylic to insulate and strengthen the joint.
– See Figure 52 for the results of this step.

**Figure 51.**
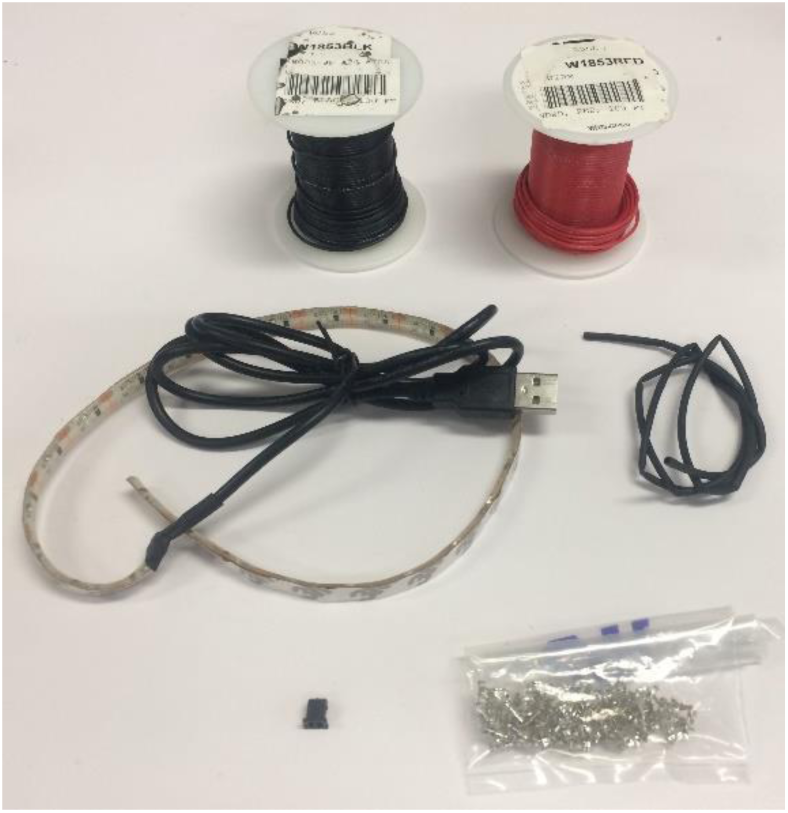
Materials for masking light. Clockwise from top left: wire, heatshrink tubing, crimp terminals, 2-pin receptacle, LED light strip

###### 3. Add receptacle

– Add crimp terminals to the two stripped ends of the wires
– Optional: twist wires
– Insert the crimped ends of the wires into the 2-pin receptacle in the order shown in Figure 55.

**Figure 52.**
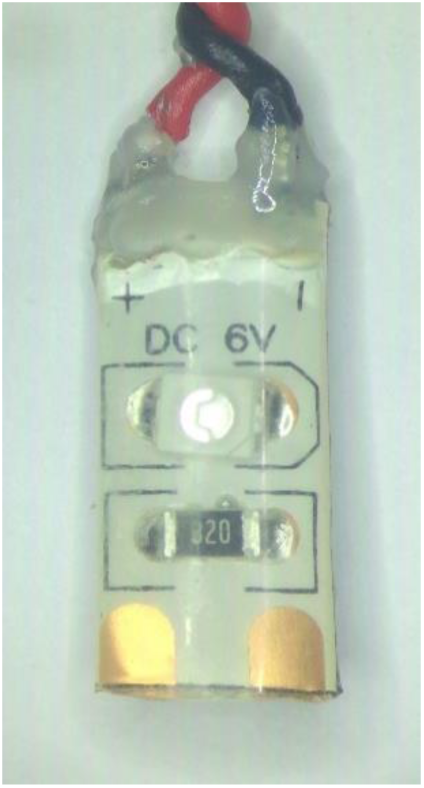
LED with terminals coated

**Figure 53.**
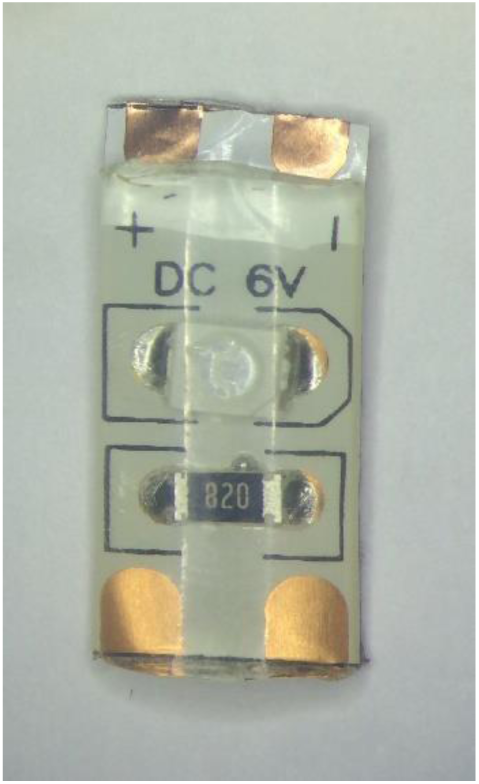
LED unit cut from strip, with terminals exposed

**Figure 54.**
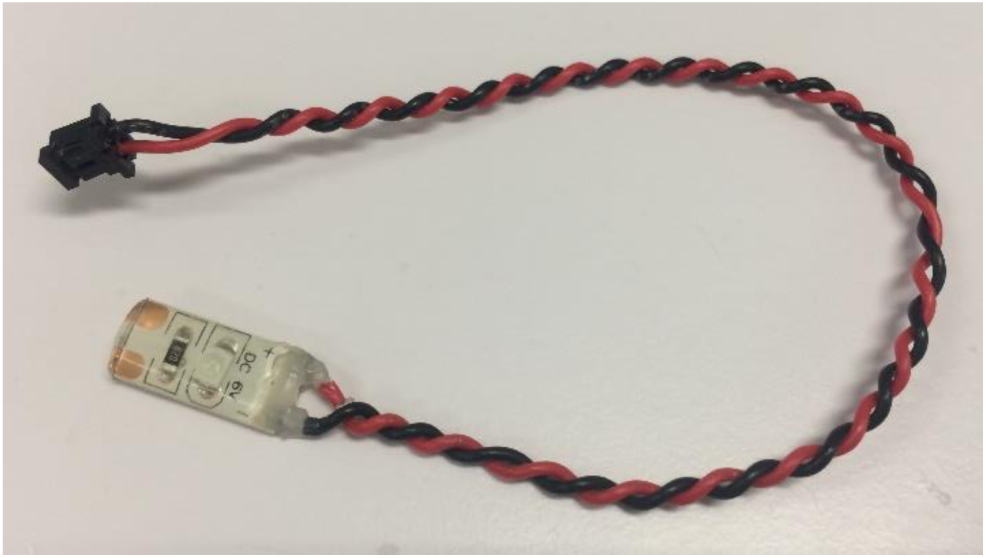
Completed masking light

**Figure 55.**
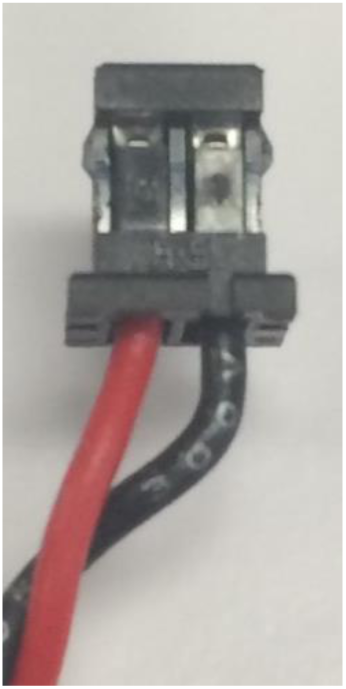
Receptacle for LED

**Figure 56.**
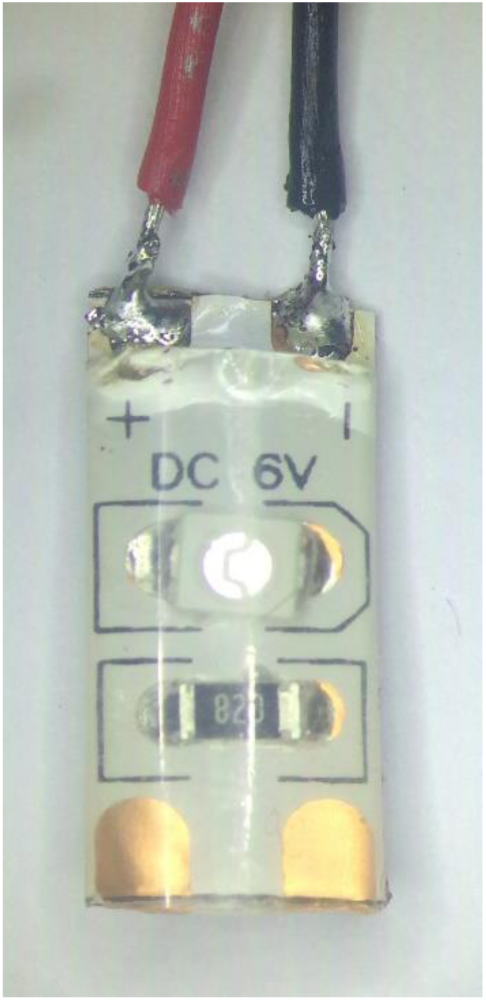
LED with wires soldered onto terminals. Note that the terminals are marked “+” and “-” to indicate which one should be connected to the 5V and GND wires

#### Part VIII: Building the water reservoir

##### Full list of Materials

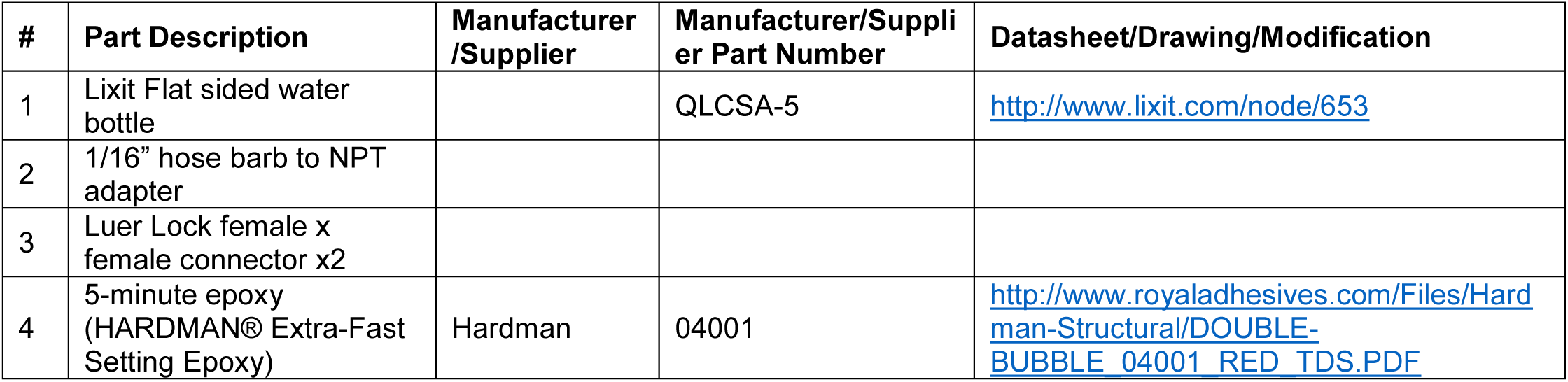

##### Building Instructions

###### General notes

The water reservoir is a modified consumer rodent water bottle. The reservoir has a hose barb port at the bottom to connect to the lick spout intake tube. It also has two luer lock ports – one for filling on top, and one for draining to a pre-set level. These can be connected to an automatic water system to easily maintain the appropriate water height in each bottle, and therefore maintain a constant dispense volume at the lick spout. The constant dispense volume can be calibrated by adjusting the duration of solenoid valve opening.

###### 1. Disassemble the bottle

– Remove the bottle from its packaging
– Unscrew the cap, separate the rubber seal and the metal spout. Discard the spout.

###### 2. Install bottom outlet adapter

– Using a utility knife, cut off the narrower rubber neck of the rubber seal, leaving a thin rubber gasket behind
– Insert the adapter in the opening in the bottle cap
– Using a combination of pushing, stretching, and screwing, push the rubber gasket onto the threaded part of the adapter, until both the adapter flange and the gasket are firmly flush against either side of the cap. If a large vise is available, it’s also possible to simply push-fit the adapter into the rubber seal using a vise, even without cutting off the seal neck or widening the opening.
– See Figure 58 and Figure 59 for the results of this step.

**Figure 57.**
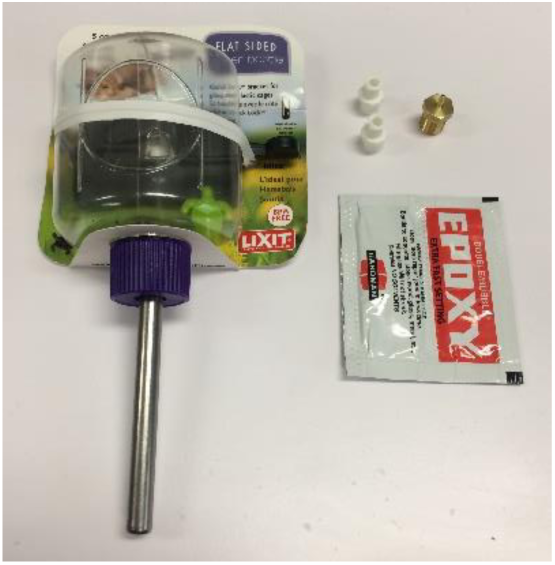
Reservoir materials. Clockwise from top left: Water bottle, luer lock connectors x2, hose barb x NPT adapter, epoxy

**Figure 58.**
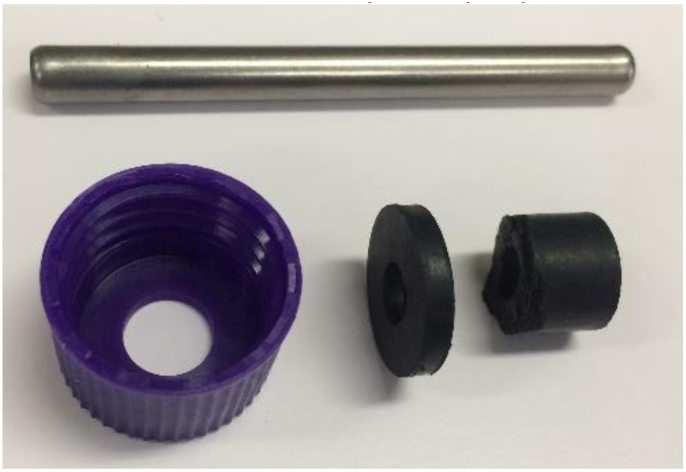
Neck removed from rubber seal. The metal spout at the top and the extra rubber on the right, can be discarded.

###### 3. Add fill and drain ports

– Optional: remove the bracket to make it easier to manipulate the bottle.
– Drill a 1/4” hole in the top and flat side of the bottle. The side hole should be approximately 1 cm below the top surface of the bottle.
– Insert the luer lock connectors into the two holes.
– Use a generous amount of epoxy between the bottle and the luer lock connector and around the lip of the connector to make a watertight seal.
– See Figure 60 for the results of this step.

**Figure 59.**
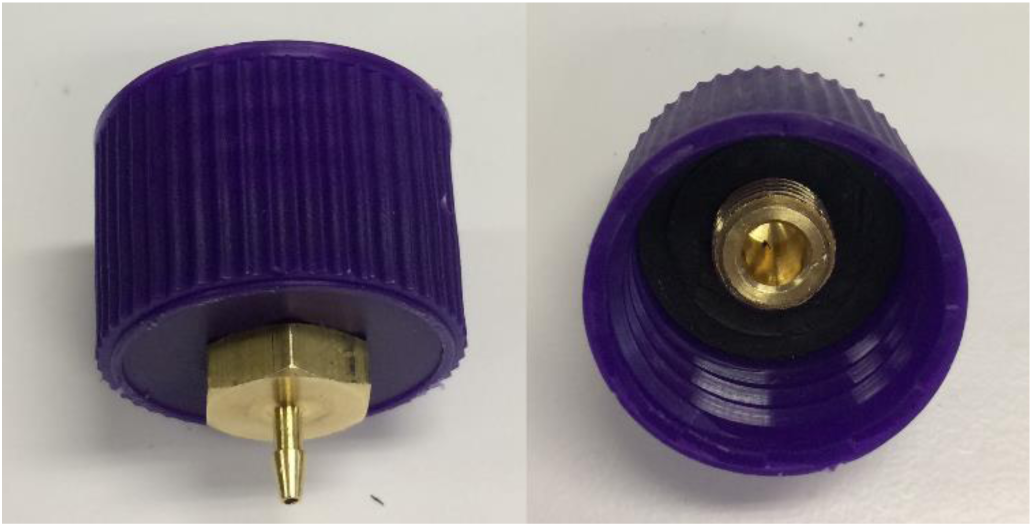
Adapter and seal installed in cap. Outside view, left, inside view, right.

###### 4. Add pressure relief hole

– Drill a 1/16” hole somewhere on the top surface of the reservoir to relieve pressure during operation.

###### 5. Screw cap back on and test seals

– Screw the cap back on
– Test to make sure the three ports are watertight by filling the bottle with water, blocking the ports off (attaching a piece of crimped tubing to each port is an easy way), and watching for drips when squeezing the bottle gently
– Make sure there are no plastic fragments left in the reservoir that could clog up the ports or tubing.
– See Figure 61 for the completed water reservoir.

**Figure 60.**
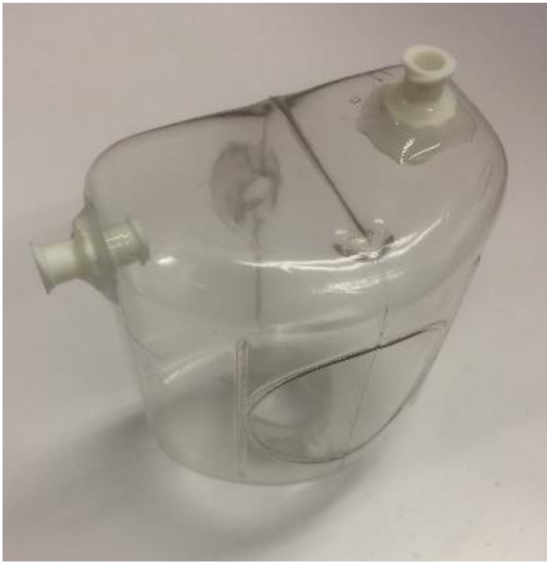
Luer lock fill and rain ports inserted and epoxied

**Figure 61.**
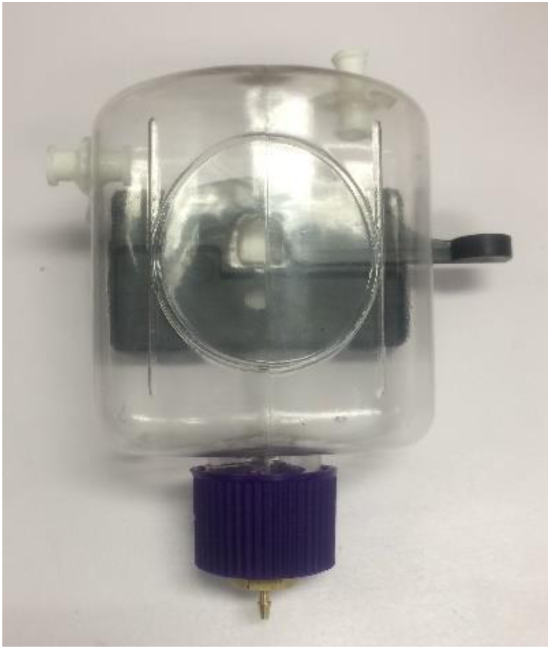
Completed water reservoir

#### Part IX: Assembling the home cage

##### Full list of Materials

| # | Part Description | Manufacturer /Supplier | Manufacturer/Supplier Part Number | Datasheet/Drawing/Modification |
| --- | --- | --- | --- | --- |
| 1 | Joystick mounted on micromanipulator |  | N.A. (Custom Made) |  |
| 2 | Protective umbrella |  | N.A. (Custom Made) |  |
| 3 | Joystick board |  | N.A. (Custom Made) |  |
| 4 | Fixed post touch sensor |  | N.A. (Custom Made) |  |
| 5 | Nosepoke sensor |  | N.A. (Custom Made) |  |
| 6 | Lick port |  | N.A. (Custom Made) |  |
| 7 | Masking light |  | N.A. (Custom Made) |  |
| 8 | Machined home cage |  |  |  |
| 9 | Water reservoir |  | N.A. (Custom Made) |  |
| 10 | L brackets x2 |  | N.A. (Custom Made) |  |
| 11 | Walls x2 |  | N.A. (Custom Made) |  |
| 12 | Lick sensor metal support panels x2 |  | N.A. (Custom Made) |  |
| 13 | Lick spout holder |  | N.A. (Custom Made) |  |
| 14 | 1/4"-20 nut |  |  |  |
| 15 | 4-40 x 3/8" length screws x10 |  |  |  |
| 16 | 4-40 nuts x8 |  |  |  |
| 17 | 8-32 bolts and nuts x2 |  |  |  |
| 18 | 4-40 D-Sub hex head screw x2 |  |  |  |
| 19 | 8-32 x 3/8" thumb screw |  |  |  |
| 20 | Copper sheet |  | 6" X 24"/ 10 Mil (30 ga.) Copper Sheet | https://basiccopper.com/6x2410milcos.html |

##### Building Instructions

###### General notes

The completed homecage contains all the components assembled in the previous sections. It is designed to be part of a scalable, high throughput, largely automated mouse training and neurobiology and behavior research system. Note that it can be helpful to have a block to set the homecage on so the components underneath aren’t supporting any weight. It can also be helpful to have a heavy block in the back end of the cage to prevent it from tipping.

**Figure 62.**
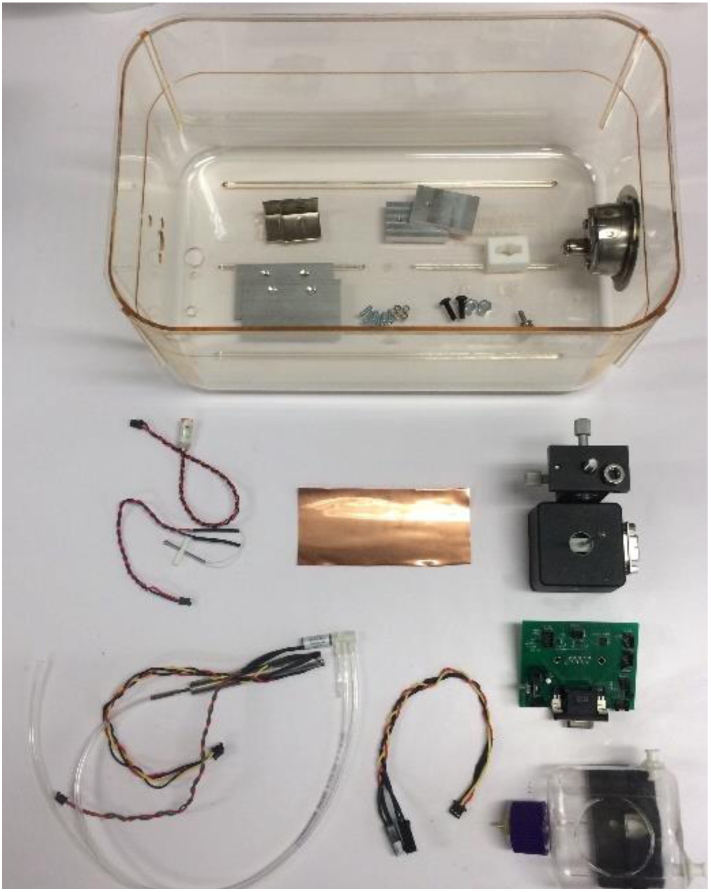
Home cage materials. Clockwise from top left: Homecage (top not pictured), L brackets, walls, lick port support panels, bolts & nuts, spout holder, joystick, joystick board, reservoir, nosepoke sensor, lick port, fixed post touch sensor, copper sheet

**Figure 63.**
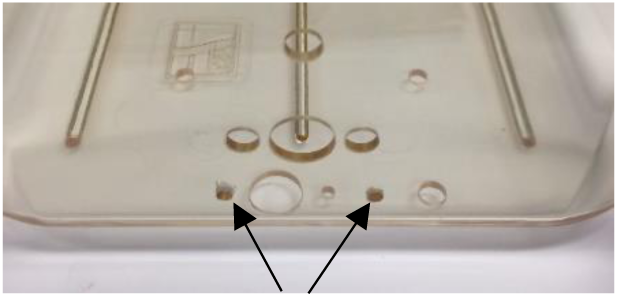
Two holes for lick port support panels on front of cage

###### 1. Drill holes for the lick port support panels

– Mark two points 3/4” above the bottom front edge of homecage, and 1 1/4” apart, centered on the nosepoke hole.
– Drill 1/8” holes on the two marks
– Using two 4-40 screws, screw each lick port support panel onto the front of the home cage. See Figure 64 for the results of this step.

###### 2. Install the lick port

– Insert the lick spout holder between the two lick port support panels, and screw it in with two 4-40 screws.
– Add a thumbscrew to the top hole on the lick spout holder. This will fun ction as a setscrew to hold the lick spout at the desired position. You may need to clear out the hole with a drill to get the thumbscrew in.
– Insert the lick spout into the lick spout holder, angle it so a mouse can just access it from the nosepoke hole, and tighten it into place with the thumbscrew. See Figure 66 for the results of this step.

**Figure 64.**
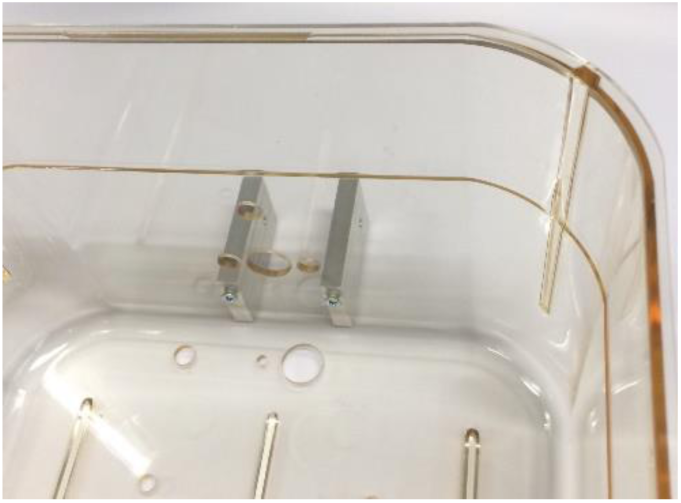
Lick port support panels installed (view from inside)

**Figure 65.**
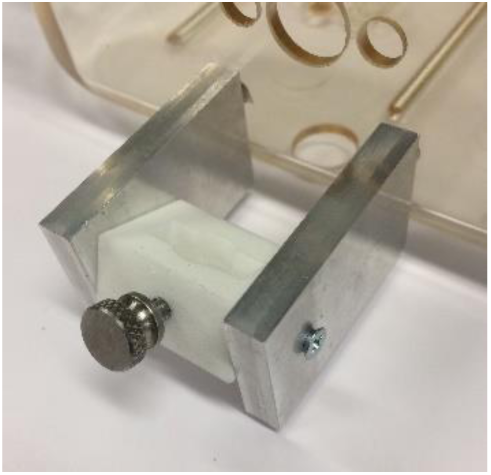
Water spout holder installed

###### 3. Adding the nosepoke sensor and masking light

– Press the nosepoke sensor transmitter and receiver through the two holes on either side of the nosepoke hole.
– You may need to either widen the nosepoke holes slightly, or round the corners of the nosepoke sensor slightly to allow it to fit – a Dremel tool works for this.
– There should be approximately 1/4” of the sensor exposed on the inside of the cage.
– Ensure the transmitter and receiver are facing each other
– Apply superglue on either side of the LED on the masking light
– Press masking light onto hole in top center of homecage, above the nosepoke hole, allow the glue to dry.
– See Figure 68 for the results of this step.

**Figure 66.**
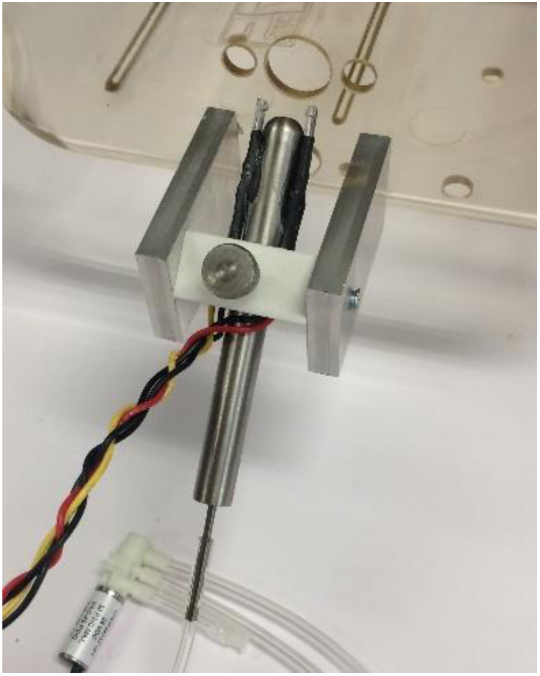
Lick port installed

###### 4. Add protective umbrella on joystick electrode

– The umbrella prevents bedding from falling out of the cage hole and into the joystick housing.
– Drill out the center hole on the protective umbrella with a 1/8” drill bit.
– Gently insert the electrode into the umbrella until it is roughly 1/4” above the joystick housing. Glue it in place with dental acrylic.
– See Figure 67 for the results of this step.

**Figure 67.**
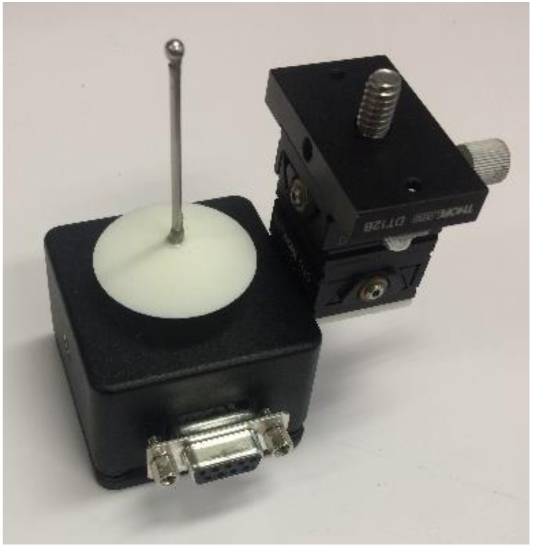
Joystick with umbrella

**Figure 68.**
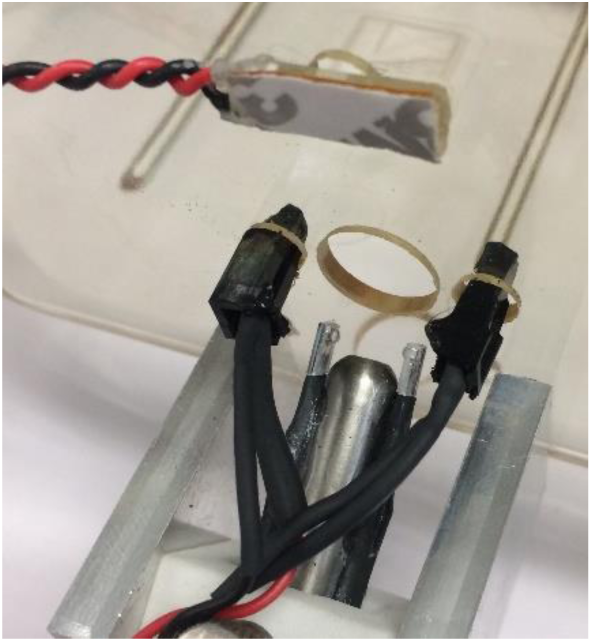
Nosepoke sensor and masking light installed

###### 5. Mounting joystick under cage

– Insert micromanipulator mounting screw and joystick post into the two holes on the bottom of the cage at the front. Loosely secure the micromanipulator with the 1/4” nut.
– Check to make sure that the top of the joystick ball is 11/16” above the cage floor.
– Make sure joystick and micromanipulator remain vertical, and the micromanipulator and joystick housing remain square to the cage, as you tighten the nut
– Tighten the nut well with a wrench so it won't shift.
– You may want to use threadlock to prevent slipping.
– Use the two adjustment knobs on the micromanipulator to precisely center the joystick in the joystick hole.
– See Figure 69 for the results of this step.

**Figure 69.**
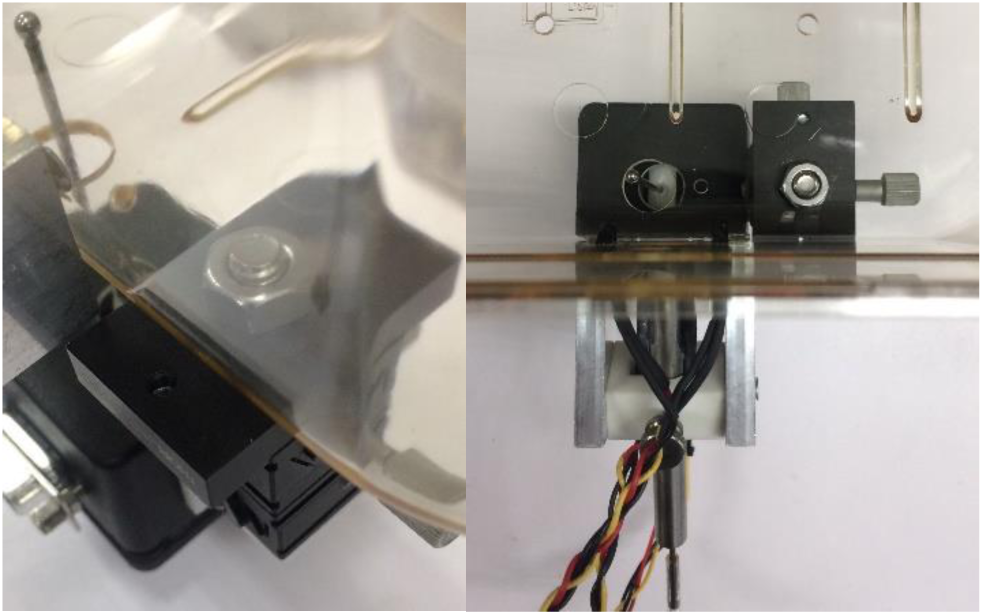
Joystick mounted on cage. Left: View from outside. Right: View from above (Note that the umbrella is missing in this photo).

###### 6. Add fixed post

– Take care not to bump the delicate joystick electrode during this process.
– Gently push post through the hole next to the joystick hole in the cage bottom from below. If there is any resistance, stop and widen the hole slightly with a 1/8” drill bit. Do not force the delicate capacitive ball through the hole!
– Superglue post spacer to underside of the cage. Do not glue the post to the spacer yet, as the height needs to be precisely adjusted.
– Position the fixed post electrode so the fixed post ball is about 1/2 an electrode ball below the joystick ball. This offset gives the animal better stability for manipulating the joystick.
– Make sure the fixed post is vertical
– Glue fixed post electrode in place
– ensure that the post doesn’t shift or tilt as the glue dries.
– See Figure 70 for the results of this step.

**Figure 70.**
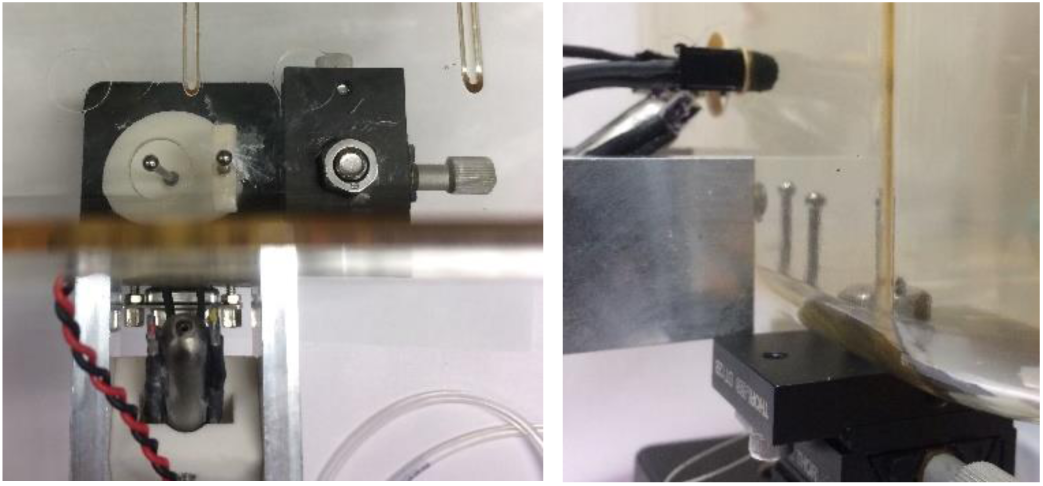
Fixed post touch sensor. Left: Top view. Right: Side view.

###### 7. Plug in JS board

– Gently plug the joystick board DB9 connector into the joystick DB9 connector. Make sure you don’t change the alignment of the joystick in the process.
– Recommended: Screw joystick board into joystick housing DB9 connector using approximately 1/2” 4-40 screws
– Connect cables as shown in Figure 72.
– See Figure 71 for the results of this step.

**Figure 71.**
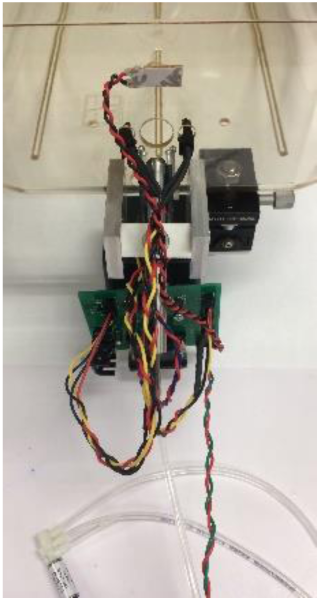
All sensors installed

**Figure 72.**
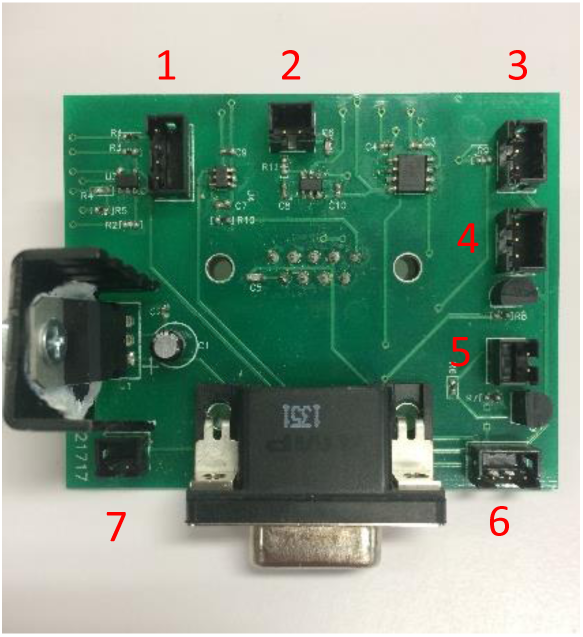
Joystick board connectors: 1: Lick sensor 2: Fixed post touch sensor 3: Nosepoke sensor 4: Solenoid valve 5: Masking light 6 & 7: No connection (extra)

###### 8. Add walls

– Countersink one side of wall holes to prevent the screw head from sticking out
– Screw wall into L-bracket
– Screw L-bracket into cage bottom so mouse has approximately 1 1/2” of space between walls to walk through.
– Take care not to damage joystick and fixed post electrodes.
– See Figure 73 for the results of this step.

**Figure 73.**
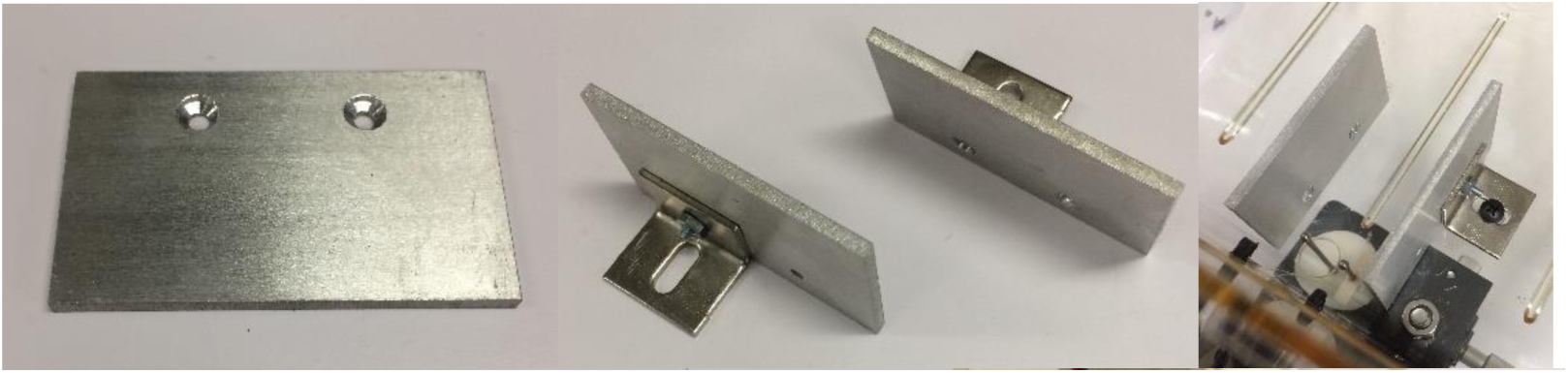
From left to right: Wall showing countersunk holes, walls with brackets attached, walls installed

###### 9. Add gutter

– The gutter will redirect any water drips away from the joystick and electronics.
– Cut a roughly 2”x6” rectangle of the sheet copper
– Fold a lip up along the long sides to contain drips
– Cut away the lip along about 1/2” of the start of the gutter, to provide an attachment flap
– Glue or tape the gutter to the lick port support panels underneath the lick port
– Flare out the sides of the lip under the lick port to maximize the protected area
– See Figure 74 for the results of this step.

**Figure 74.**
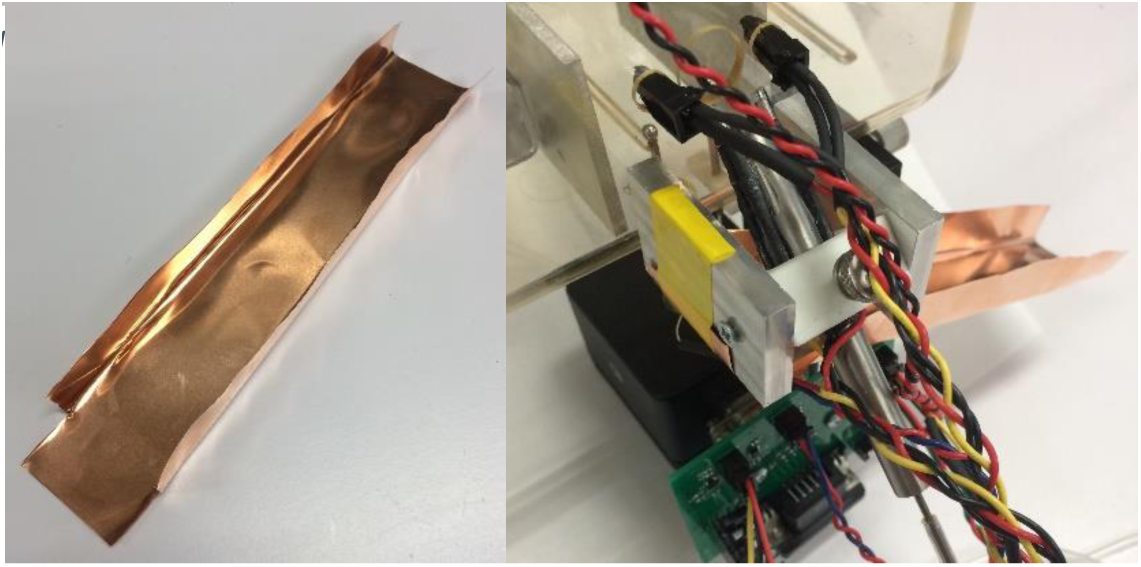
Gutter. Left: Formed, Right: Installed

###### 10. Assemble cage lid

– The cage lid consists of a commercially available cage lid machined to have a large opening on the top. A custom cover fits in the large opening in the cage lid, and has a long slot to allow freely moving mice to be attached to electrodes or fiber optics. The reservoir also attaches to front of the cage lid
– Mark the corresponding locations of the four holes around the perimeter of the opening in the lid on the custom cover.
– Drill four 1/8” holes where the four marks are on the custom cover
– See Figure 75 for the custom cover with drilled holes
– Remove the protective film from both sides of the custom cover
– Screw the custom cover onto the cage lid with 4-40 x 3/8” screws, and secure with nuts.
– Remove the adhesive backing on the mounting bracket for the water reservoir and firmly press it onto the front lip of the cage lid off to one side.
– See Figure 77 for the results of this step.

**Figure 75.**
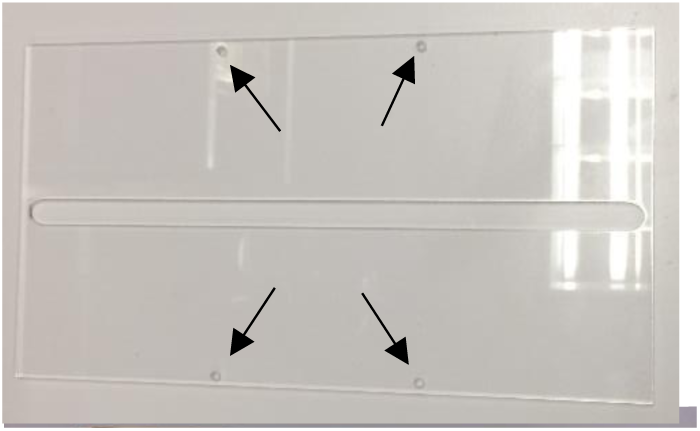
Custom cover with four holes drilled

**Figure 76.**
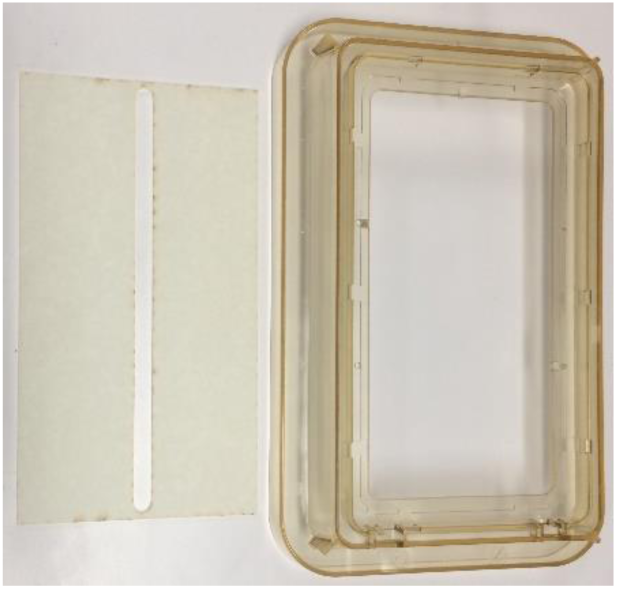
Custom cover and cage lid

**Figure 77.**
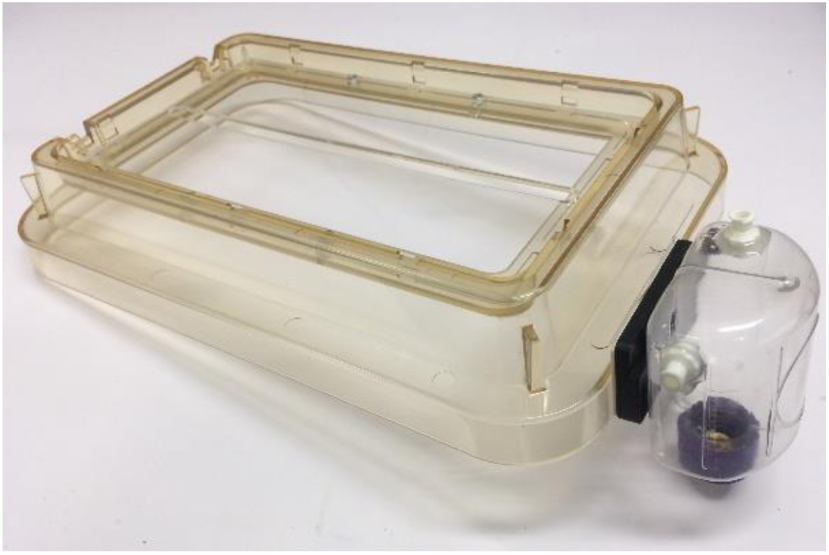
Completed cage lid

**Figure 78.**
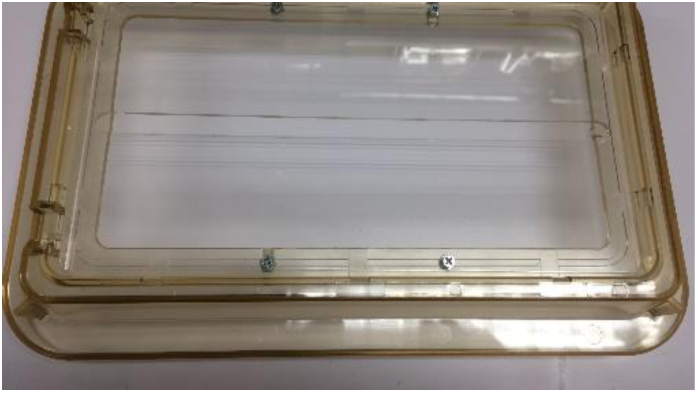
Cage lid with custom cover

**Figure 79.**
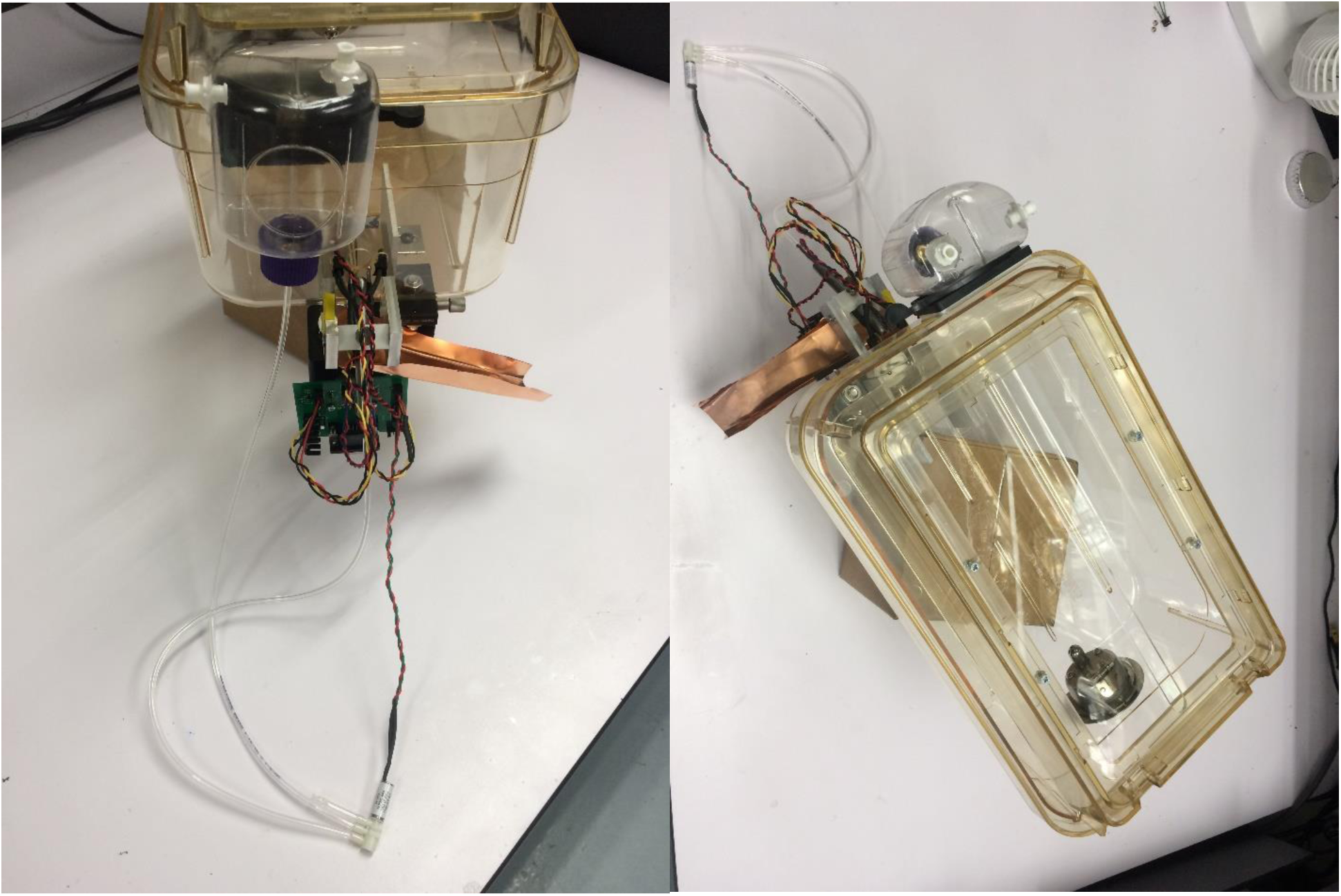
Completed home cage

#### Part X: Building the optogenetic system

##### Full list of Materials

| # | Part Description | Manufacturer /Supplier | Manufacturer/Supplier Part Number | Datasheet/Drawing/Modification |
| --- | --- | --- | --- | --- |
| 1 | Doric Laser Diode Fiber Light Source, 2 channel, 450 nm |  | LDFLS_450/075_450 /075 | http://doriclenses.com/life-sciences/laser-diode-fiber-light-sources/910-laser-diode-fiber-light-source-2-channel-model.html#ld_wavelengths-450_nm_75_mw/ld_wavelengths_2-450_nm_75_mw |
| 2 | Fiber commutator |  | FRJ_1x2i_FC-2FC | http://doriclenses.com/life-sciences/fiber-optic-rotary-joints/809-1x2-fiber-optic-rotary-joints-intensity-division.html |
| 3 | Mounting plate for commutators |  | N.A. (Custom Made) |  |
| 4 | Fiber optic cable, male to male SMA, Thor Labs |  | FG200UCC 200um 0.22NA |  |
| 5 | Fiber optic cable, male SMA to ferrule insert |  |  |  |
| 6 | Doric commutator mounting bracket |  | Holder_FRJ_large | http://doriclenses.com/life-sciences/holders-for-rotary-joints/819-holder-for-frj1x2-frj1x4-erj-aerj-hrj-oe-and-ledfrj-1ch.html |
| 7 | 1/4"-20 x 1" screws |  |  |  |
| 8 | 1/4"-20 nuts |  |  |  |

##### Building Instructions

###### General notes

The optogenetic system consists of a laser light source.

The light is sent via fiber optic cable to a fiber optic cranial implant on a mouse. To allow the mouse to freely turn in the cage, an optical commutator is used in the optical path. The commutator mounts on a custom plate that sits in the rack above each homecage.

###### 1. Screw commutator into mounting bracket

– See Figure 81 for the results of this step.

**Figure 80.**
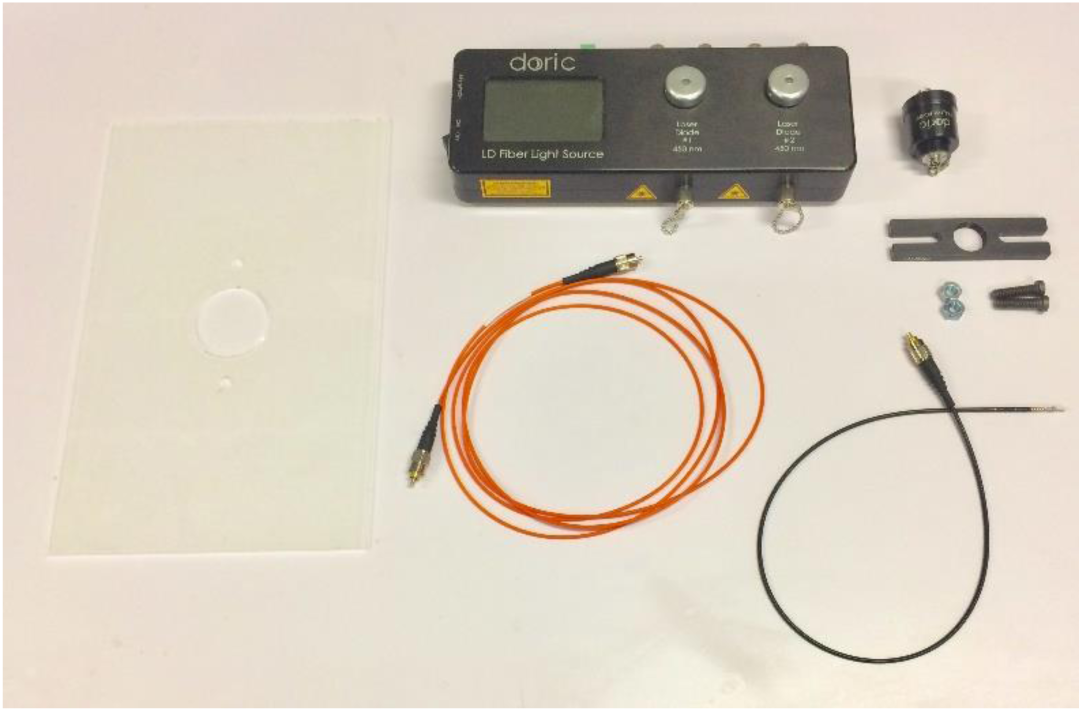
Optogenetic system materials. Clockwise from top left: Custom mounting plate, laser light source, commutator, commutation mounting bracket, 1/4 “x20 bolts and nuts, fiber optic cable (black, male SMA to ferrule), fiber optic cable (orange, male to male)

###### 2. Bolt bracket to custom mounting plate

– Use the 1/4”x20 bolts and nuts to fix the commutator roughly in the middle of the opening in the plate.
– Note that the bracket should go on top, so in the unlikely event that the screws loosen and come out, the commutator will not fall.
– See Figure 82 for the results of this step.

**Figure 81.**
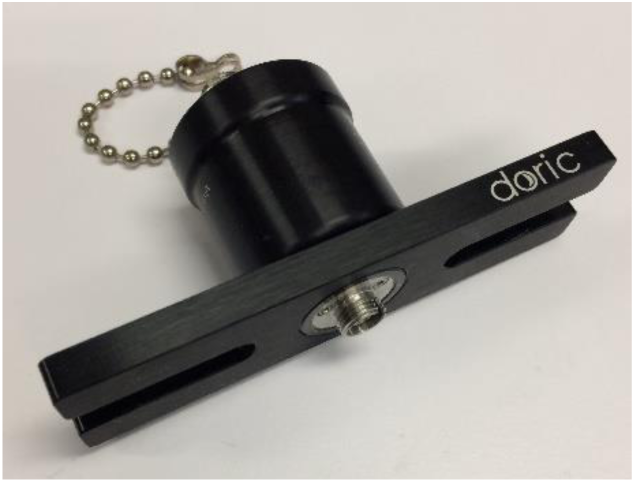
Commutator screwed into

###### 3. Connect the fiber optic cables

– The male-male cable should go on top (the non-rotating port). The other end of this cable will connect to the laser light source.
– The male-ferrule cable should go on the bottom (the rotating port). The other end of this cable will connect to the implant.
– See Figure 84 for the results of this step.

**Figure 82.**
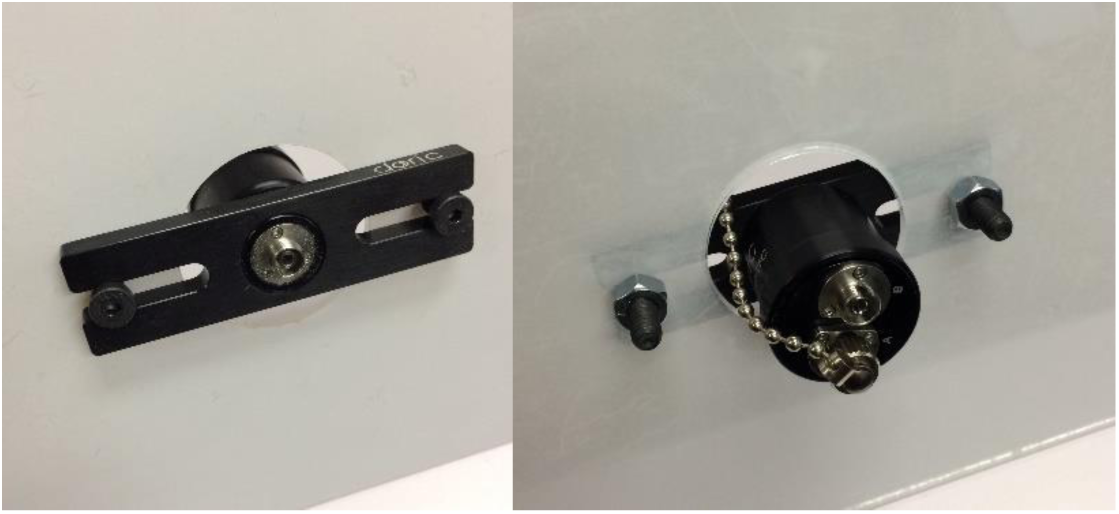
Commutator and bracket mounted to custom plate. Left: View from top. Right: View from bottom

###### 4. Use tape to hold excess cable in place

– Take the extra fiber optic cable that connects the laser to the commutator and coil it on the top of the mounting plate.
– Use tape to fix the coil in position so it won’t shift during the experiment.
– Note that changes in the radius of curvature of the fiber optic cable during the experiment will cause the light power level reaching the implant to change.
– See Figure 83 for the results of this step.

**Figure 83.**
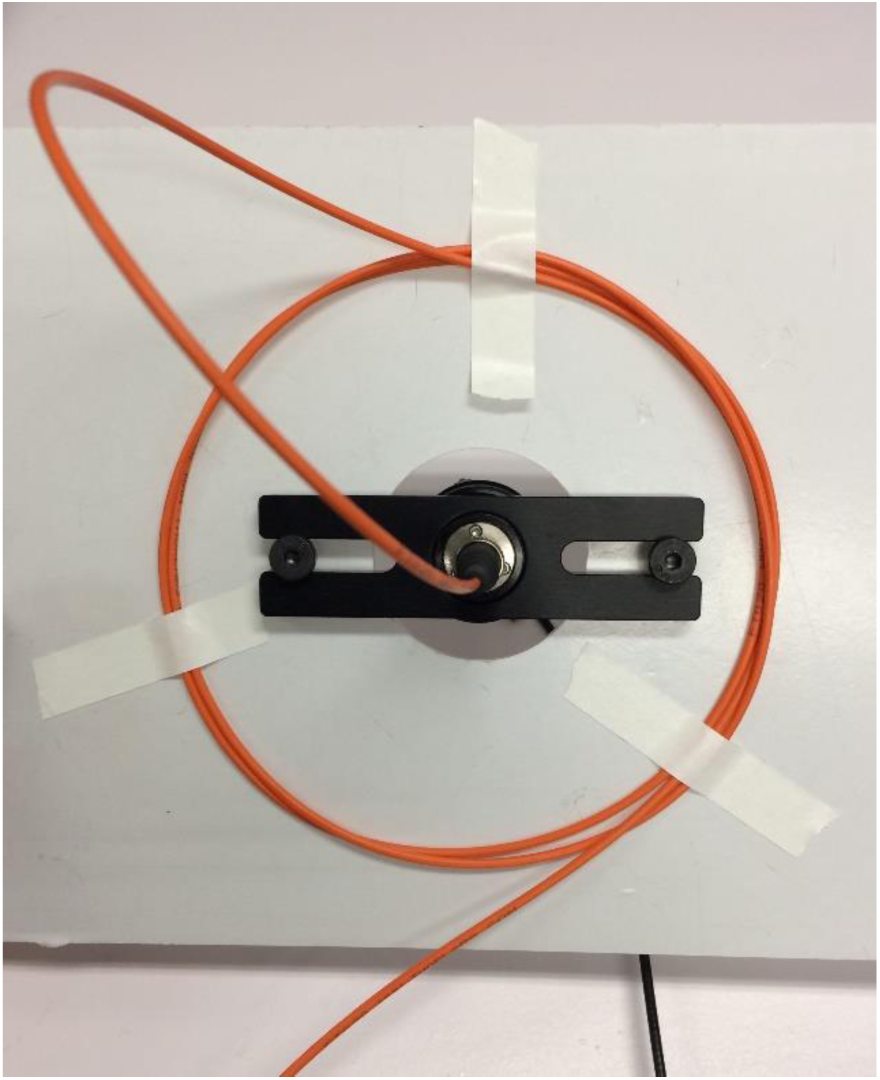
Cable management

**Figure 84.**
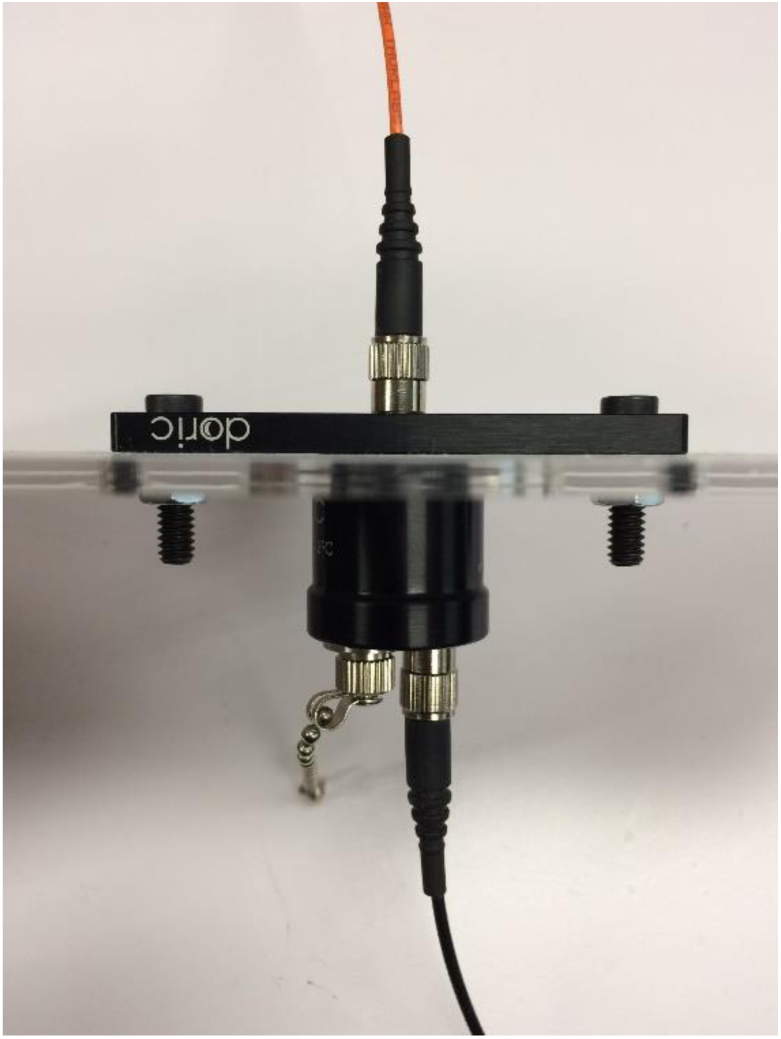
Fiber optic cables connected

#### Part XI: Building the RIO breakout board

##### Full list of Materials

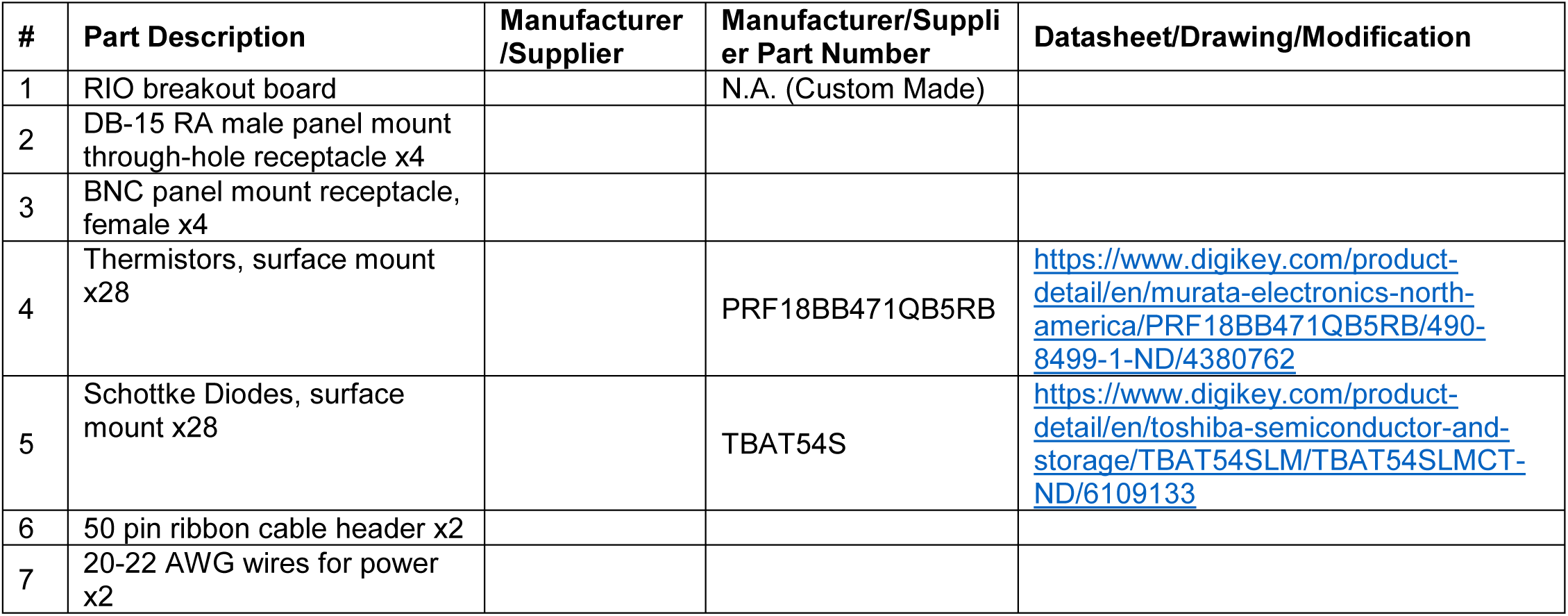

| # | Part Description | Manufacturer /Supplier | Manufacturer/Supplier Part Number | Datasheet/Drawing/Modification |
| --- | --- | --- | --- | --- |
| 1 | RIO breakout board |  | N.A. (Custom Made) |  |
| 2 | DB-15 RA male panel mount through-hole receptacle x4 |  |  |  |
| 3 | BNC panel mount receptacle, female x4 |  |  |  |
| 4 | Thermistors, surface mount x28 |  | PRF18BB471QB5RB | https://www.digikey.com/product-detail/en/murata-electronics-north-america/PRF18BB471QB5RB/490-8499-1-ND/4380762 |
| 5 | Schottke Diodes, surface mount x28 |  | TBAT54S | https://www.digikey.com/product-detail/en/toshiba-semiconductor-and-storage/TBAT54SLM/TBAT54SLMCT-ND/6109133 |
| 6 | 50 pin ribbon cable header x2 |  |  |  |
| 7 | 20-22 AWG wires for power x2 |  |  |  |

##### Building Instructions

###### General notes

The RIO breakout board is a component of the DAQ system. It plays two roles – protecting the analog and digital IO ports of the expensive sbRIO board, and routes the sbRIO IO channels into four groups and connects them to DB-15 and BNC receptacles so they can be easily connected to the homecages and laser light sources. Each RIO breakout board services a single sbRIO board, and supports up to four homecages.

###### 1. Solder the SMT components

###### 2. Solder the through-hole components

###### 3. Cut and strip power wires

– Cut and strip two wires, about 1 foot each.
– Color coding is recommended – color scheme used here: red=high, white=low
– Solder into power connection through holes in RIO breakout board.

#### Part XII: Building the data acquisition system (DAQ)

##### Full list of Materials

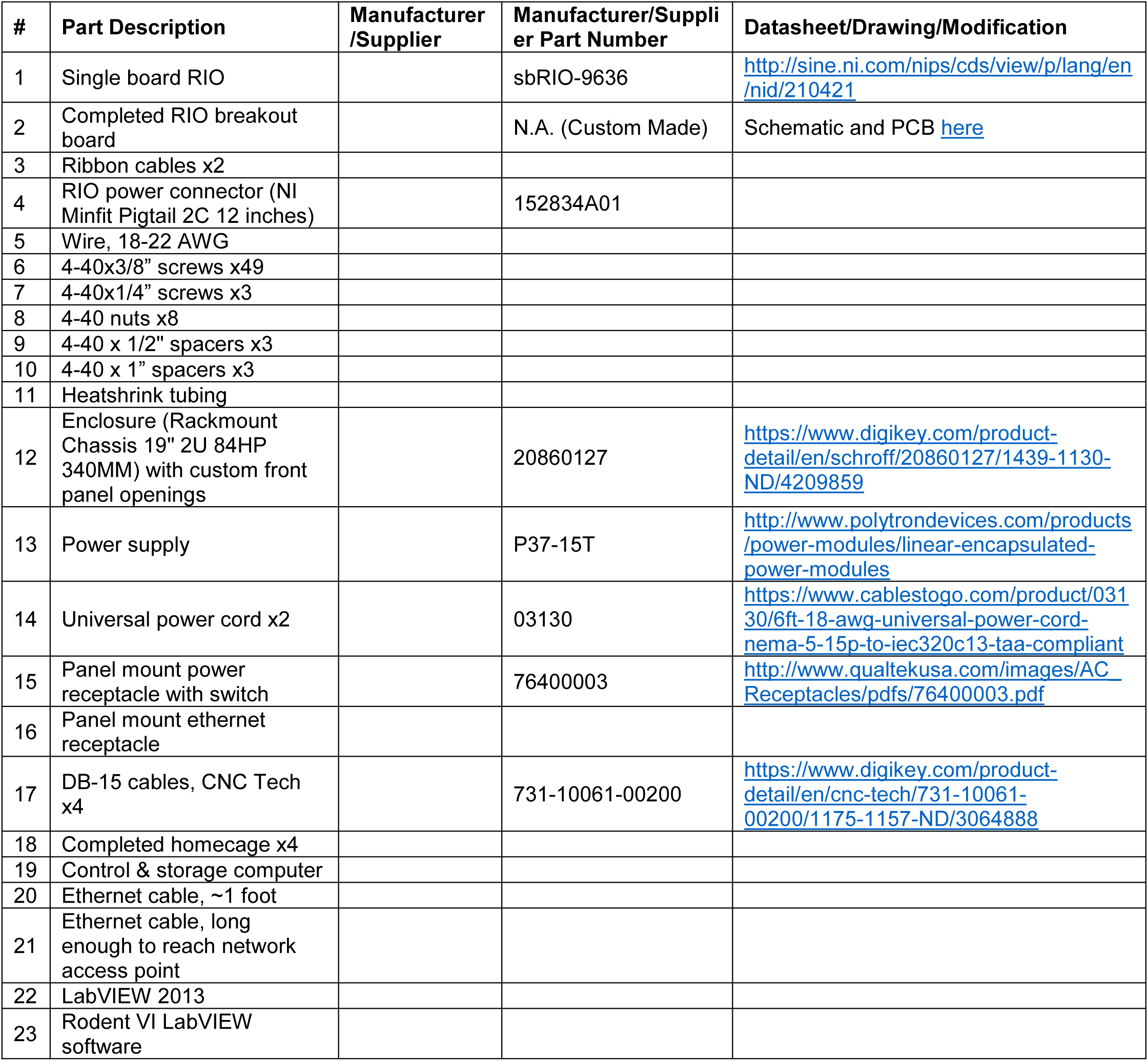

##### Building Instructions

###### General notes

The data acquisition system consists of a single-board RIO DAQ from National Instruments, which has a large array of digital and analog input/output (IO) channels, a custom PCB that protects and distributes those channels, an enclosure for the DAQ, DB-15 cables to connect the DAQ to each homecage, and a control/storage computer which controls the DAQ via LabVIEW software, and stores the sensor data it receives. Each DAQ system can host up to four homecages.

**Figure 85.**
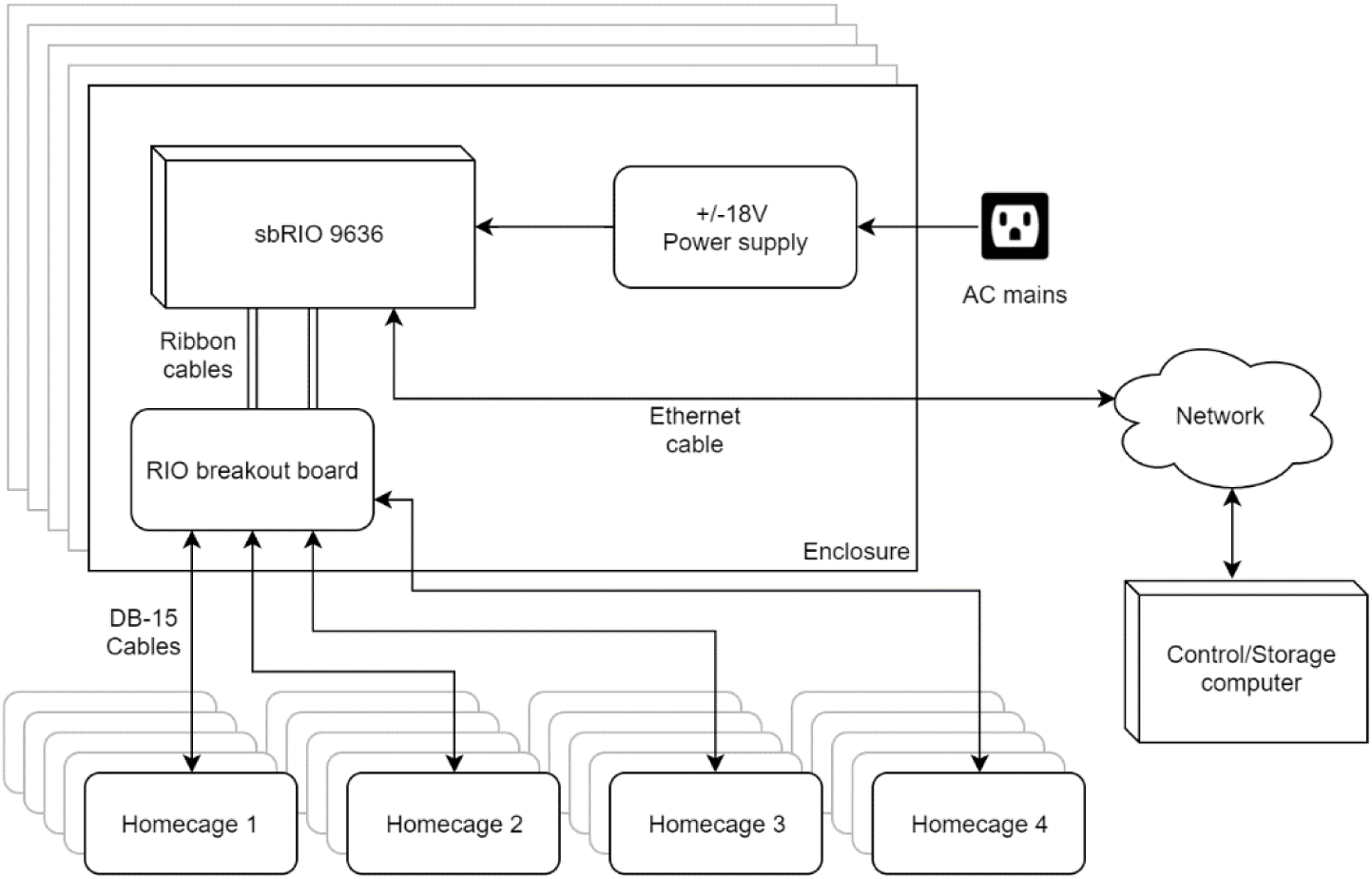
DAQ system schematic

**Figure 86.**
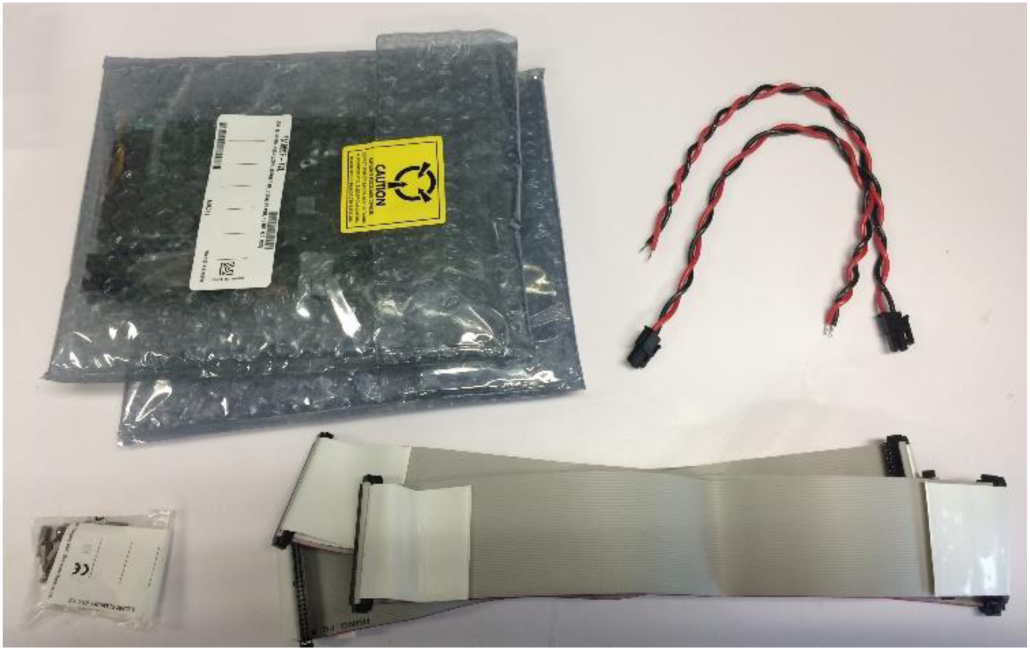
sbRIO supplies. Clockwise from top left: sbRIO-9636 x2, NI Minifit Pigtail x2, NI 50 pin ribbon cable x2, mounting hardware (not used)

**Figure 87.**
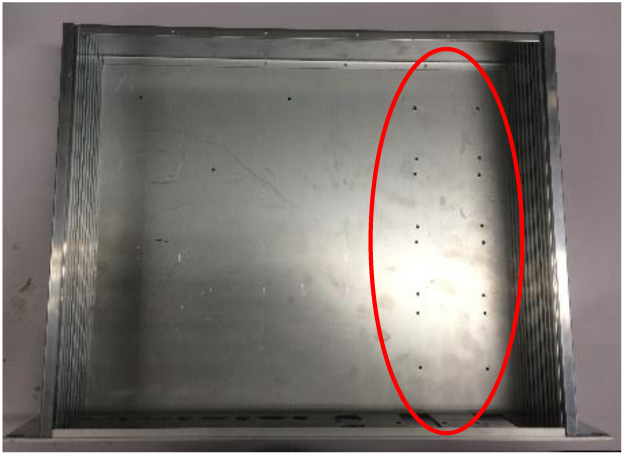
Enclosure base plate with power supply mounting holes drilled

**Figure 88.**
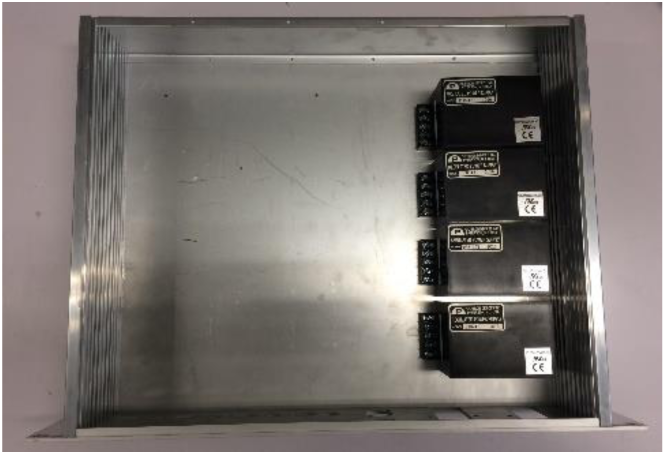
Four power supplies mounted

**Figure 89.**
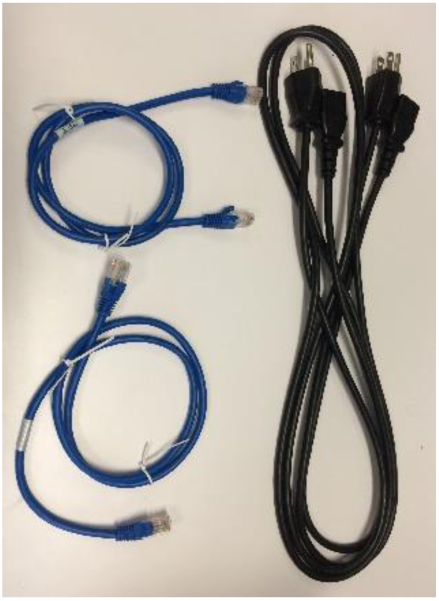
Required cords: 2 short ethernet cords, 2 power cords

**Figure 90.**
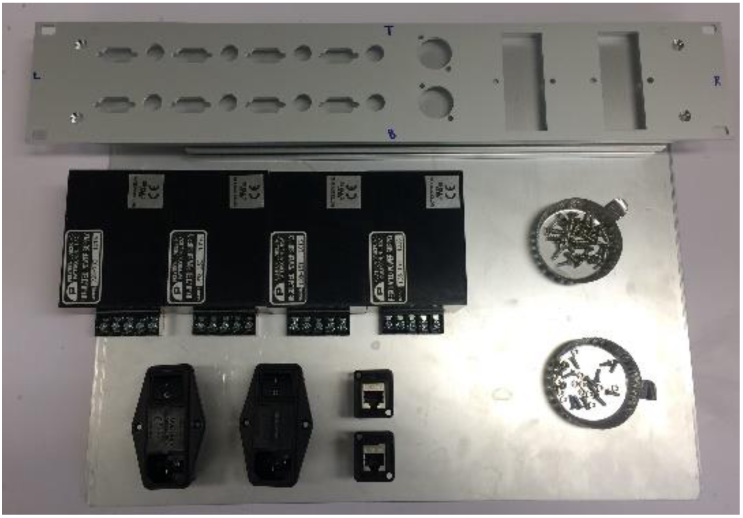
Enclosure materials. Clockwise from top left: custom cut front, sides, and back panels, enclosure hardware, 4-40 screws and nuts, ethernet jacks x2, power jacks x2, power supplies x4

###### 1. Assemble base and sides of enclosure

– See manufacturer instructions

###### 2. Mount power supplies on enclosure base

– Each power supply has four 4-40 mounting holes.
– Drill four mounting holes (1/8”) for each power supply along the right side of the base plate of the enclosure
– Screw each power supply in using 4-40×1/4” screws
– See Figure 87 and Figure 88 for the results of this step.

**Figure 91.**
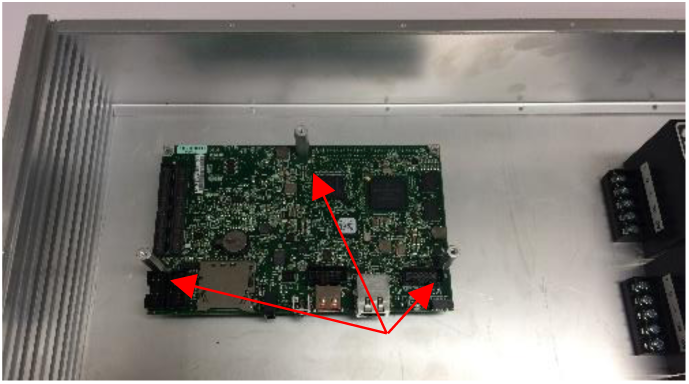
2nd layer of standoffs installed

**Figure 92.**
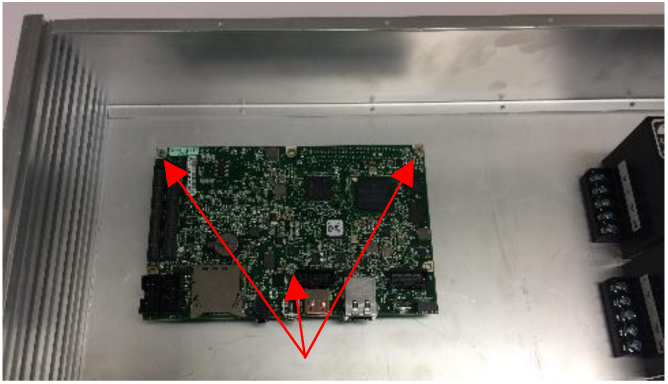
sbRIO #1 installed on standoffs

**Figure 93.**
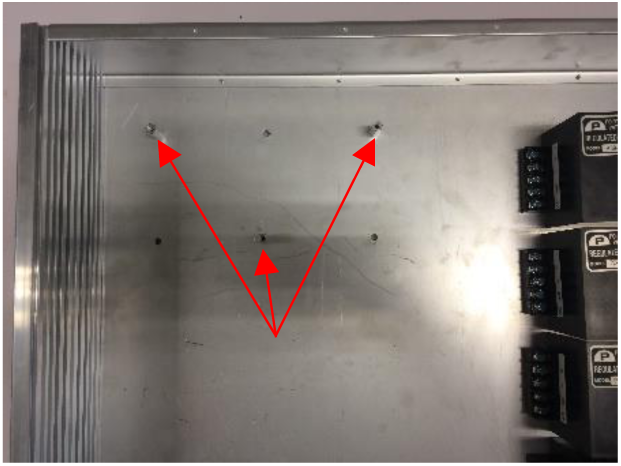
Baseplate with three standoffs installed. Note that the image shows the other three holes drilled – these will be unused and unnecessary.

###### 3. Mount sbRIOs on enclosure base

– Drill mounting holes (1/8”) for three of the six mounting holes in the sbRIO board. See Figure 93.
– Note that the mounting standoffs provided by National Instruments do not provide enough clearance for the ribbon cables when mounting two sbRIOs on top of each other.
– Note that unless you have male to female standoffs, you can only use three mounting holes for each sbRIO.
– Screw the shorter three standoffs into the enclosure base
– Mount an sbRIO on top of the three standoffs. See Figure 92.
– Screw the longer three standoffs into the three unused mounting holes on the first sbRIO. See Figure 91.
– Plug two ribbon cables into the two 50-pin headers on the sbRIO board. These cables carry the DIO and AIO channels to and from the sbRIO. Make sure the side of the ribbon marked red is on the side of the header with channel #1. See Figure 96.
– Screw the 2^nd^ sbRIO onto the 2^nd^ layer of standoffs. Take care that the ribbon cables from the first sbRIO are neatly exiting the space between the sbRIOs. See Figure 94.
– Plug the other two ribbon cables into sbRIO #2. Again take care that the orientation is correct. See Figure 95.

**Figure 94.**
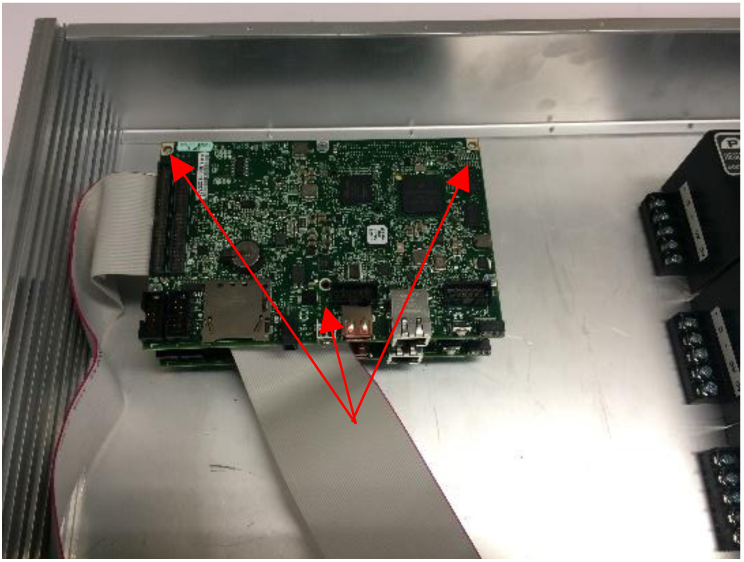
sbRIO #2 installed

**Figure 95.**
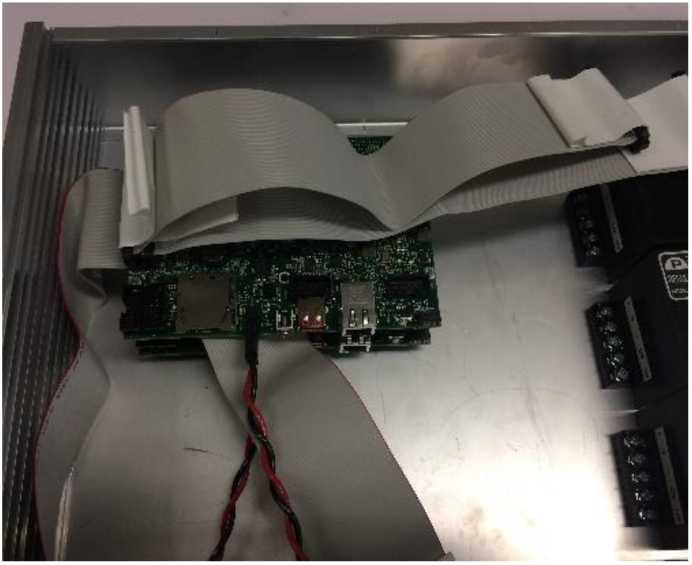
sbRIO #2 ribbon cables plugged in

**Figure 96.**
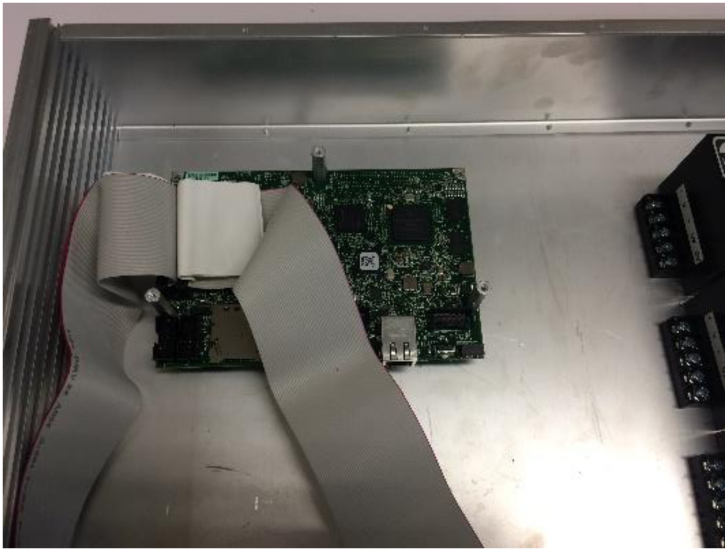
sbRIO #1 installed with ribbon cables

###### 4. Mount RIO breakout boards on enclosure faceplate

– Insert the RIO breakout board DB-9 and BNC ports into the faceplate cutouts.
– Secure the RIO breakout boards with 4-40×3/8” screws on either side of each DB-9 port
– Plug the four ribbon cables into the RIO breakout boards. In the orientation shown, the left ribbon cables from the sbRIOs go into the left headers on the RIO breakout boards. Pin #1 goes on the right.
– See Figure 97 for the results of this step.

**Figure 97.**
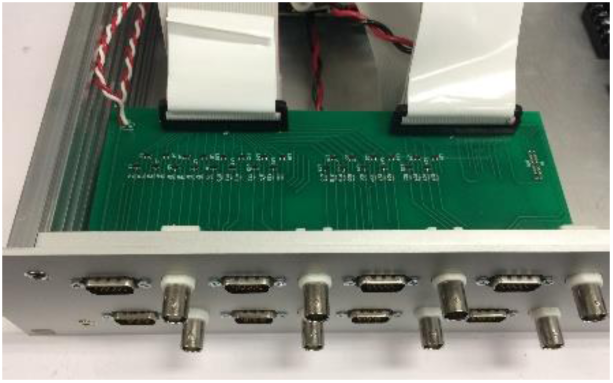
Two RIO breakout boards installed with ribbon cables.

**Figure 98.**
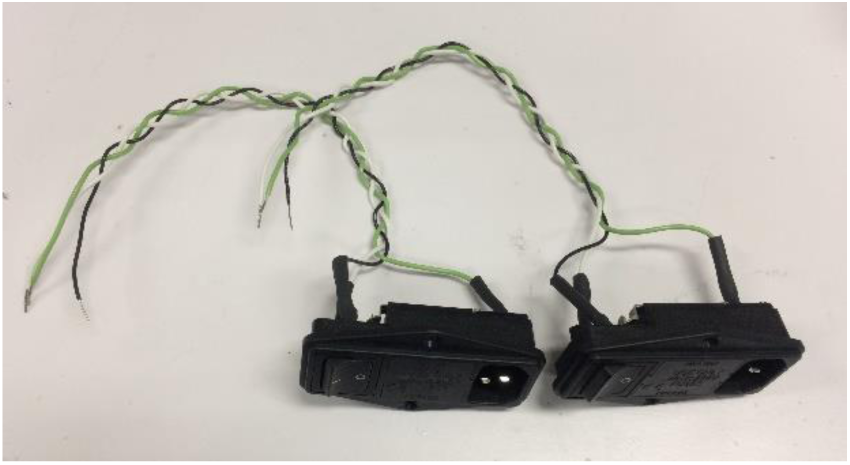
AC power receptacles

###### 5. Cut and strip power wires

– Cut three pairs of 12” wires (two each for AC hot, neutral, and ground), and three pairs of 6” wires (two each for AC hot, neutral, and ground). Color coding is recommended – the color scheme used here is

  ○ AC hot = black
  ○ AC neutral = white
  ○ AC ground = green
– See Figure 99 for the results of this step.

###### 6. Solder wires onto power receptacles, and mount on faceplate

– Solder the longer wires onto the two power receptacles. Use heatshrink tubing to protect the solder joint.
– Use 4-40 screws and nuts to mount each receptacle on the faceplate.
– See Figure 98 for the results of this step.

**Figure 99.**
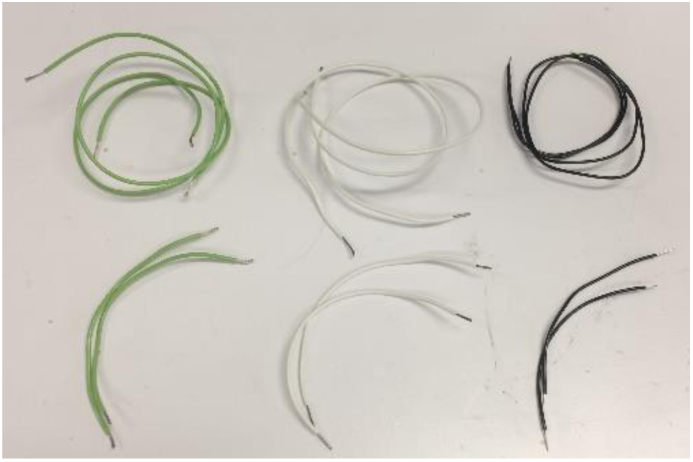
Power supply wires, cut and stripped

**Figure 101.**
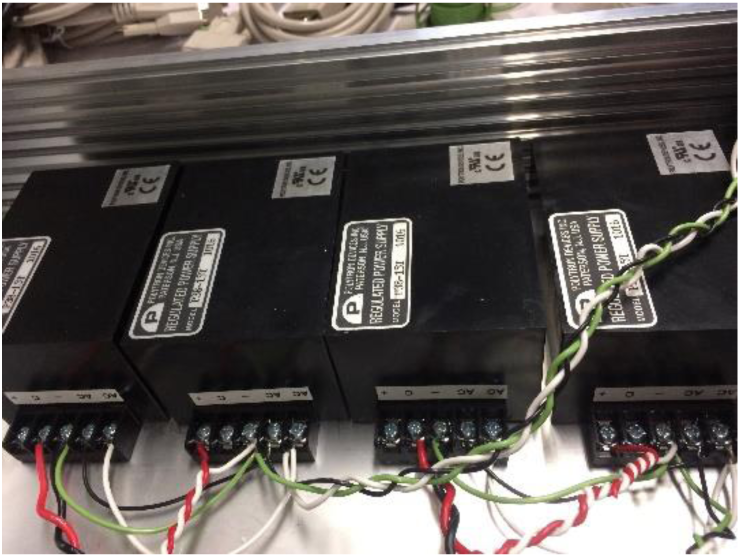
Power supply connection photo

**Figure 100.**
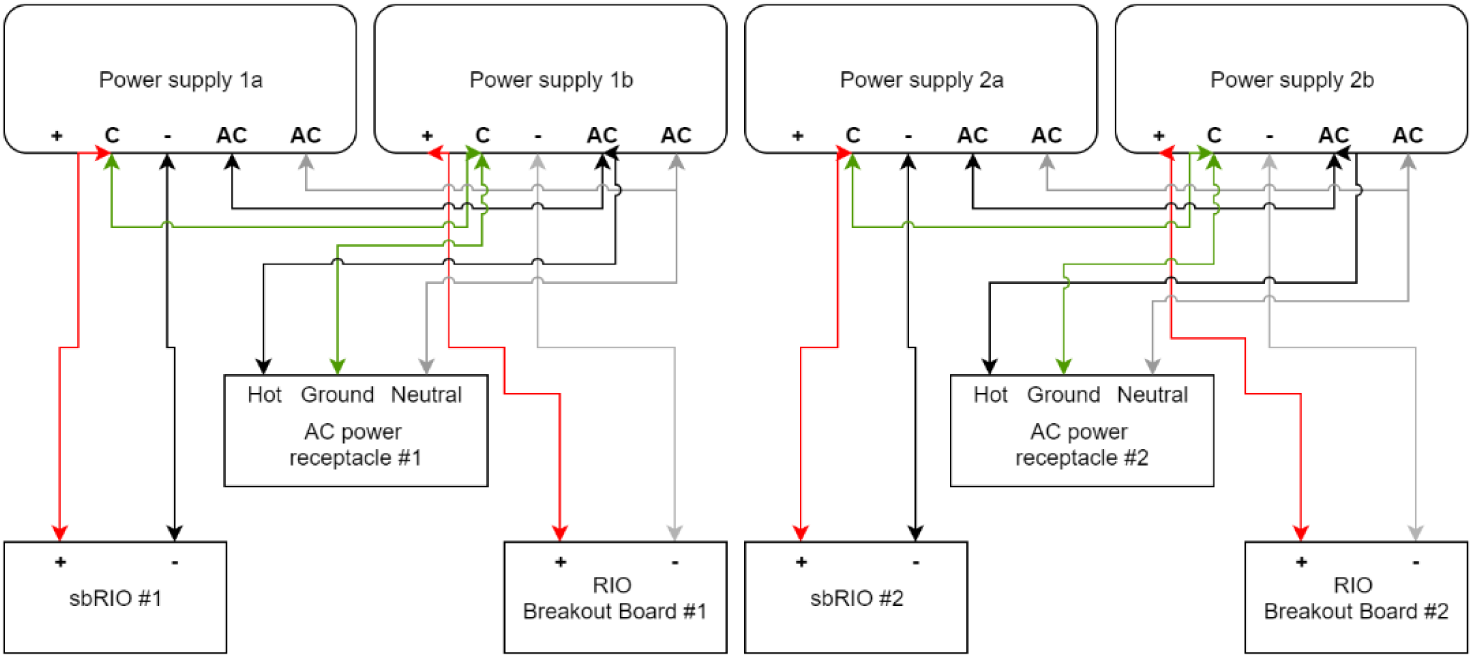
Power supply connection schematic. Compare to corresponding photo.

###### 7. Connect power wires to power supplies

– Connect the wires to the screw terminals on the power supplies as shown in Figure 100 and Figure 101.

###### 8. Connect sbRIO and RIO breakout board power cables

– Plug minfit power connectors into each sbRIO power jack.
– Connect bare wire ends of the sbRIO power cables and the RIO breakout board power cables to power supplies as shown in Figure 100 and Figure 101.

###### 9. Mount ethernet jacks in enclosure, and connect to sbRIO boards

– Mount ethernet jacks on faceplate with 4-40 screws and nuts
– Connect each sbRIO ethernet jack to the panel-mounted jack using the short ethernet cables.

**Figure 102.**
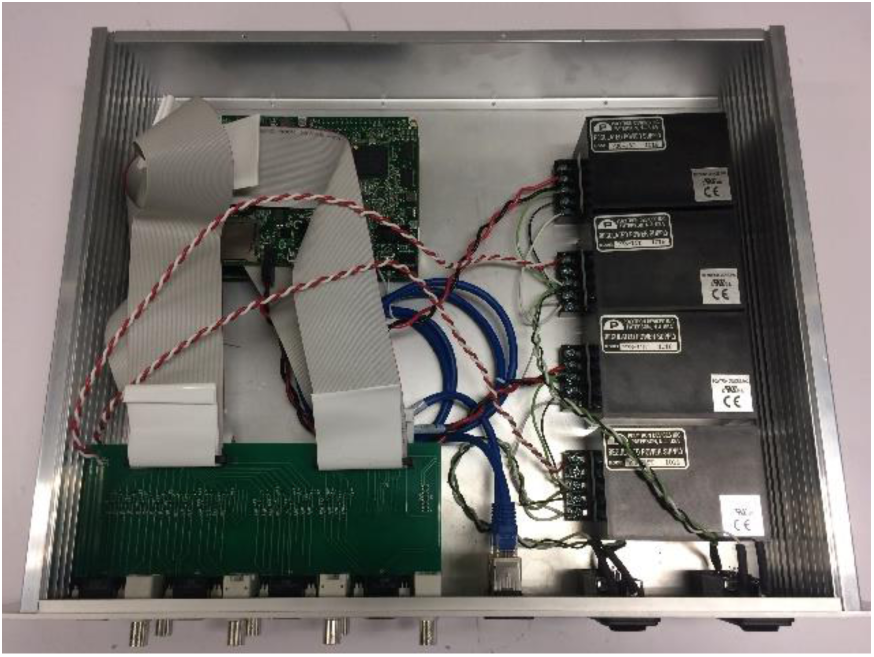
Completed DAQ system

###### 10. Optional: close enclosure with lid

###### 11. Connect to sbRIO using LabVIEW, and compile software onto FPGA

#### Part XIII: Building the water distribution system

##### Full list of Materials

| # | Part Description | Manufacturer /Supplier | Manufacturer/Supplier Part Number | Datasheet/Drawing/Modification |
| --- | --- | --- | --- | --- |
| 1 | 1/16" ID tubing |  |  |  |
| 2 | 1/8" ID tubing |  |  |  |
| 3 | 1/4" ID tubing |  |  |  |
| 4 | 1/2" ID tubing |  |  |  |
| 5 | Luer lock T connectors |  |  |  |
| 6 | Luer lock to 1/16" barb adapter |  |  |  |
| 7 | Luer lock to 1/8" barb adapter |  |  |  |
| 8 | 1/4" barb T |  | 57024-04 |  |
| 9 | 3/8" NPT male coupler |  |  |  |
| 10 | Thread sealant tape |  |  |  |
| 11 | Pneumadyne 8-station 3/8" NPT aluminum manifold |  | M10-125-8 | https://www.pneumadyne.com/station-npt-input-aluminum-manifold-p-2633.html |
| 12 | Supply pump: Brushless DC Pump |  | AD20P-1230D |  |
| 13 | Return pump: Flojet 3.8 GPM 45 PSI self-priming pump |  | 04300501 |  |
| 14 | Globe valve x2:<br>Milwaukee Valve 125 SWP 200 WOG 3/8" NPT to 3/8" NPT |  | 502 1/8"-3" |  |
| 15 | Plastic bucket with a lid, roughly 2 gallons |  |  |  |

##### Building Instructions

###### General notes

This closed loop water recirculation system is designed so it is easy and quick to accurately refill all the reservoirs in a rack on a daily basis; this allows the water dispense volume to remain calibrated down to the uL over the course of the experiment. These instructions, and the schematic in Figure 103, are for a water system for 8 homecages. This design can be easily scaled up or down to accommodate a different number of homecages.

**Figure 103.**
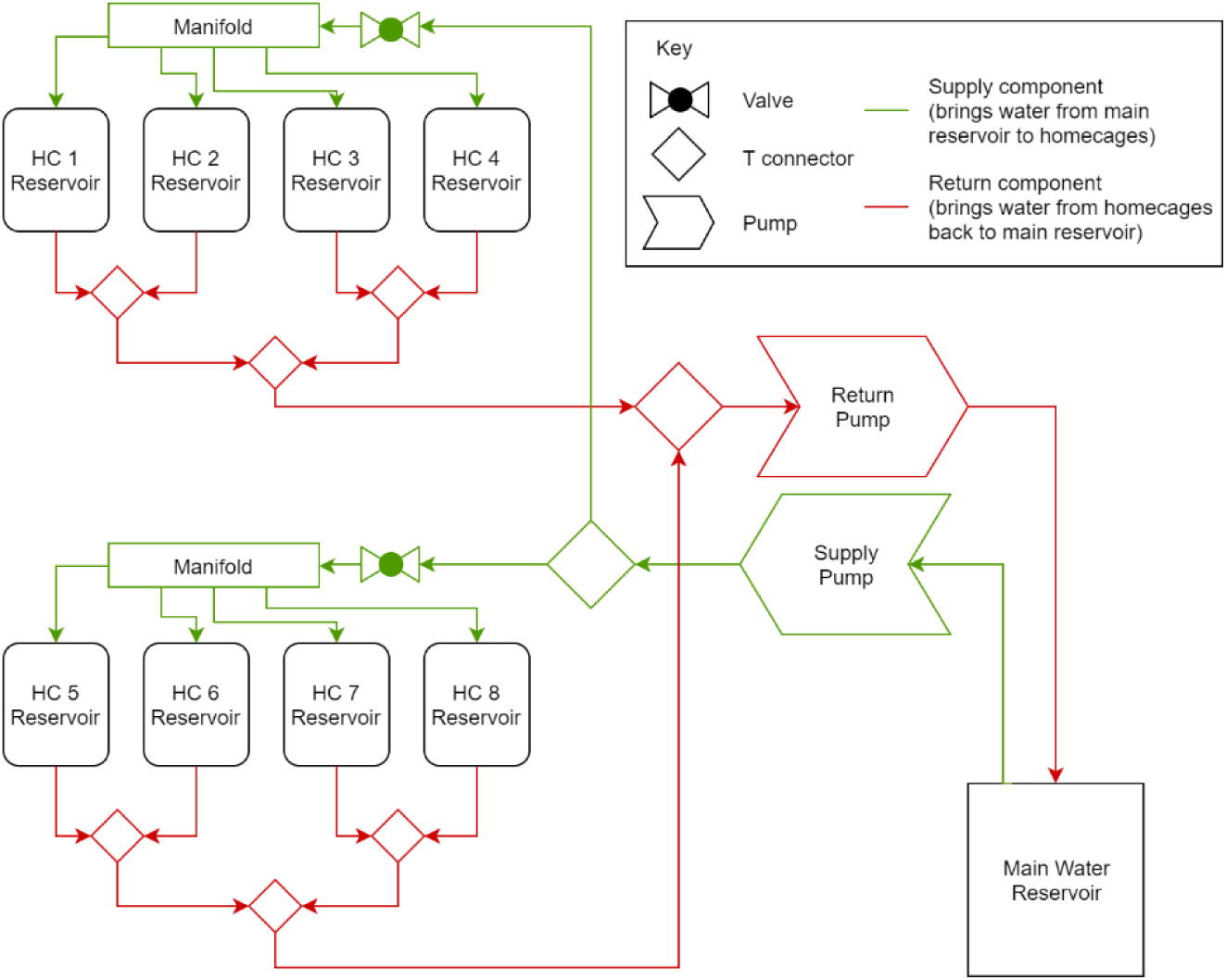
Water distribution system schematic

**Important**: Whenever there are NPT threaded connectors, you must seal the thread with thread sealant tape and tighten with a wrench for a watertight seal.

###### 1. Cut return tubing sections to length

– The tubing lengths required for this setup depend on the rack geometry in which they will be installed, and convenient mounting locations for the pumps. Refer to Figure 104 for the topology of the tubing.

###### 2. Connect return tubing

– The return tubing forms a branching tree structure, with luer lock T connectors at each branch, and luer lock to barb adapters connecting the tubing to the Ts. A 3/8” to 1/4” barb adapter steps up tubing diameter towards the diameter of the return pump inlet.
– Firmly connect all tubing onto all the barbs (make sure tubing passes over all the ridges on each barb for a watertight connection), and screw in all the luer lock connections.
– Refer to Figure 104 for the return tree tube connections

**Figure 104.**
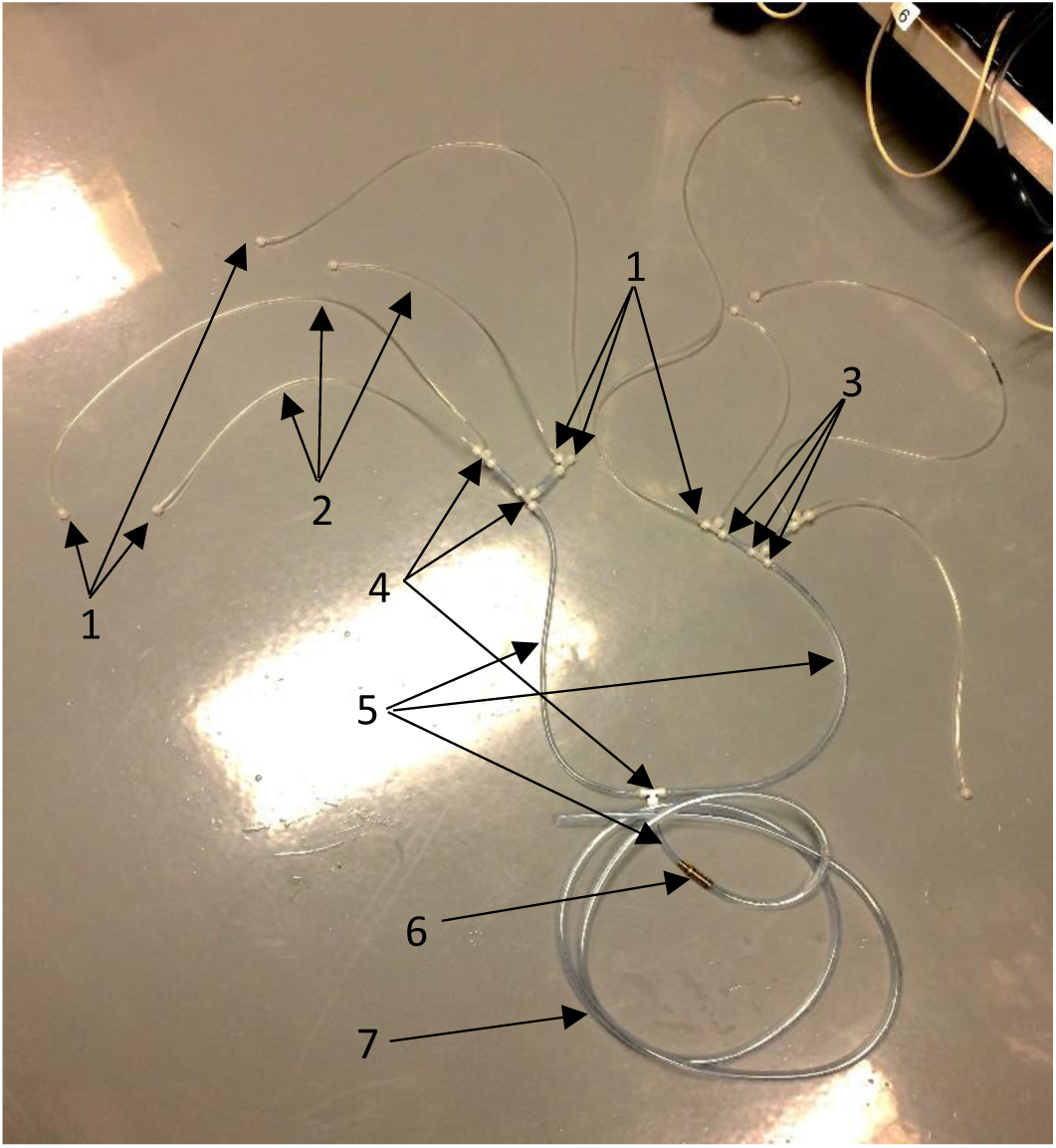
Return tubing. Examples of each component are numbered above and identified below. 1. Luer lock to 1/16” barb adapters 2. 1/16” tubing 3. Luer lock to 1/8” barb adapters 4. Luer lock T connectors 5. 1/8” tubing 6. 1/8” to 1/4” adapter 7. 3/8” tubing

###### 3. Connect return tubing to return pump

– Using a 1/4” to 3/8” barb adapter, and a short segment of 3/8” tubing, connect the return tree to the return pump.
– Attach a section of 3/8” tubing to the return pump outlet – this will lead to the main water reservoir.
– Optional: Mount the pump on a panel that allows it to neatly fit in one of the rack spaces.
– The white strips on either side of the pump are foam strips that reduce the transfer of vibration to the rack when the pump is turned on.
– See Figure 106 for the pump setup. Note that the pump is mounted on a black plastic panel in this figure.

###### 4. Assemble manifold

– Screw in 3/8” NPT to 1/16” barb adapters into the eight ports of each manifold.
– Block off four ports on each manifold with a small piece of 1/16” tubing and plug
– Screw in a 3/8” NPT male coupler to the end port on each manifold
– Screw the globe valves onto the couplers
– Screw a 3/8” NPT to 1/8” barb adapter onto each globe valve
– See Figure 105 for the results of this step. Note that the manifold is mounted on a black plastic rack-adapter panel in this figure.

**Figure 105.**
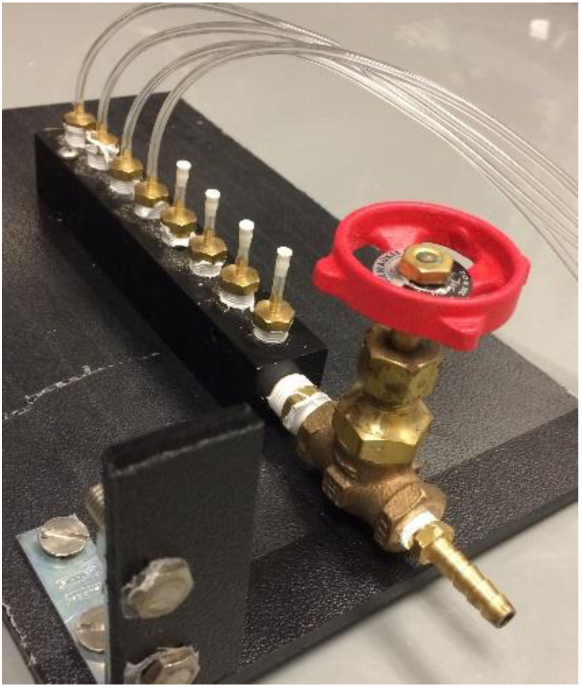
Assembled manifold

**Figure 106.**
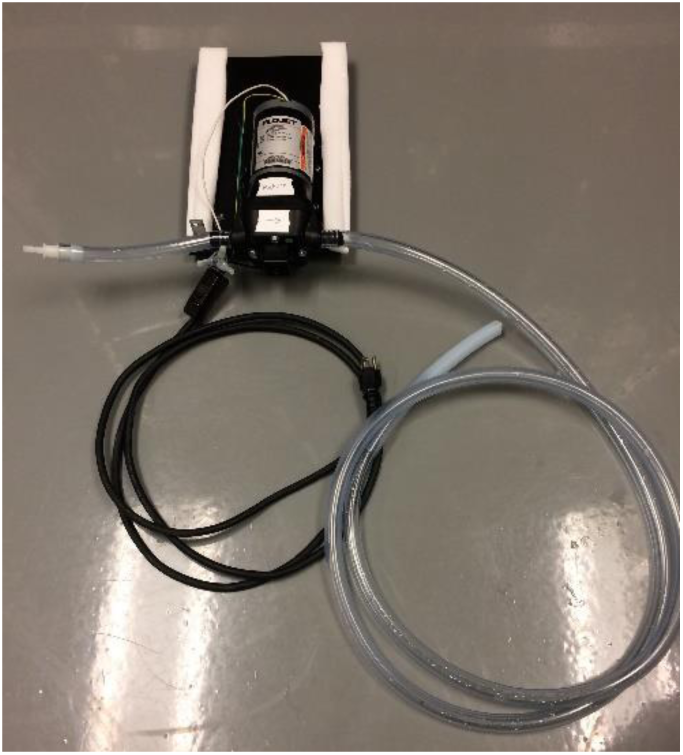
Return pump, showing the inlet tube on the left, the outlet tube on the right.

###### 5. Cut supply tubing sections to length

– As with the return system, the tubing lengths will depend on the geometry of the setup.
– Refer to Figure 107 for the topology of the supply system

###### 6. Connect supply tubing

– Refer to Figure 107 for tubing connections

###### 7. Recommended: Add priming valve

– To make priming the supply pump easier, add a T in the main supply line and add a valve that can be opened to manually pull liquid through the supply pump (see part #7 in Figure 107)

**Figure 107.**
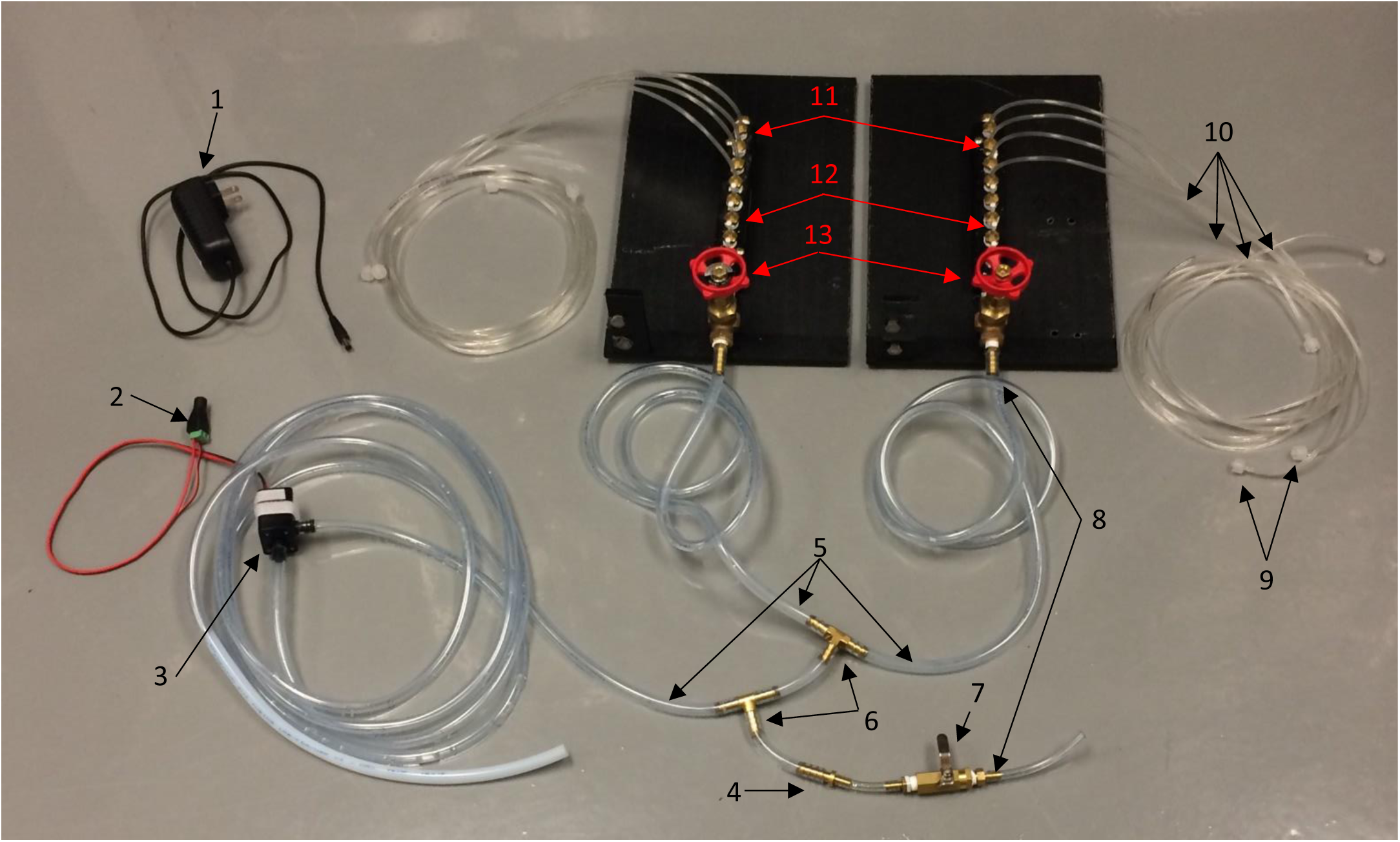
Supply system. Examples of each component are numbered above and identified below. 1. Pump power supply 2. Barrel to screw terminal adapter for pump power supply 3. Supply pump (with white Velcro for mounting) 4. 1/4” barb to 1/8” barb adapter (due to lack of supplies) 5. 1/4” tubing 6. 1/4” barb T 7. Priming valve 8. 3/8” NPT to 1/8” barb adapter 9. Luer lock to 1/16” barb adapter 10. 1/16” tubing 11. 3/8” NPT to 1/16” barb adapter 12. Short pieces of 1/16” tubing with 1/16” plugs 13. Globe valves 14. 3/8” NPT male coupler (not visible, connects valve to manifold)

###### 8. Recommended: Mount manifolds on rack near home cages

– Organizing the tubing on the rack is made easier if the manifold is mounted in some manner on the rack.
– See Figure 109 or our custom rack-adapter mounting panel.

###### 9. Construct main water reservoir

– Drill a 3/8” and 3/4” hole in the top of the water reservoir bucket.
– Fill high enough with potable water that both tubes will easily stay submerged.
– Insert the outlet tube from the return pump and the inlet tube to the supply pump into the two openings.
– Check to make sure the ends of the tubes won’t shift and emerge out of the water

**Figure 109.**
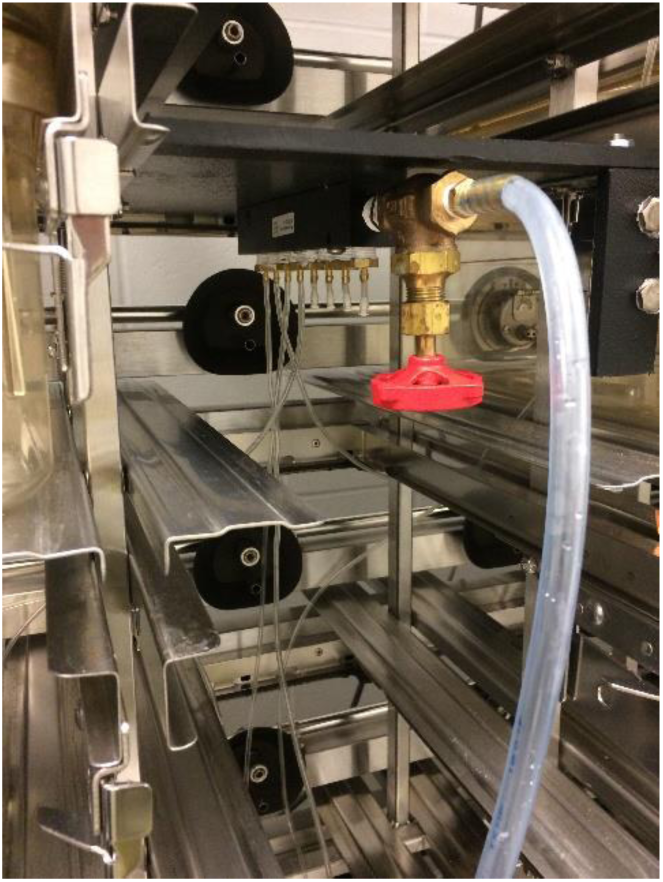
Manifold mounted on the rack

###### 10. Hook up supply and return tubes to each homecage reservoir

– Run the 1/16” supply and return tubes through the rack to each homecage
– Connect the luer lock fittings so the supply tubes connect to the top of each reservoir, and the return tubes connect to the side of each reservoir.
– See Figure 112 for the results of this step

**Figure 110.**
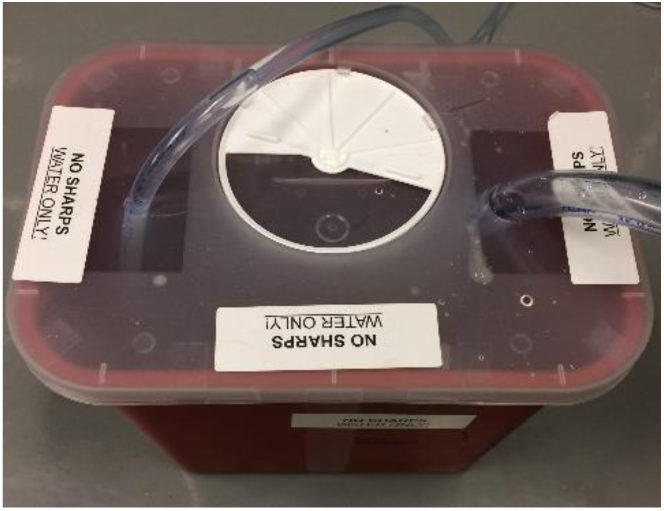
Main reservoir with holes drilled and tubes inserted.

**Figure 111.**
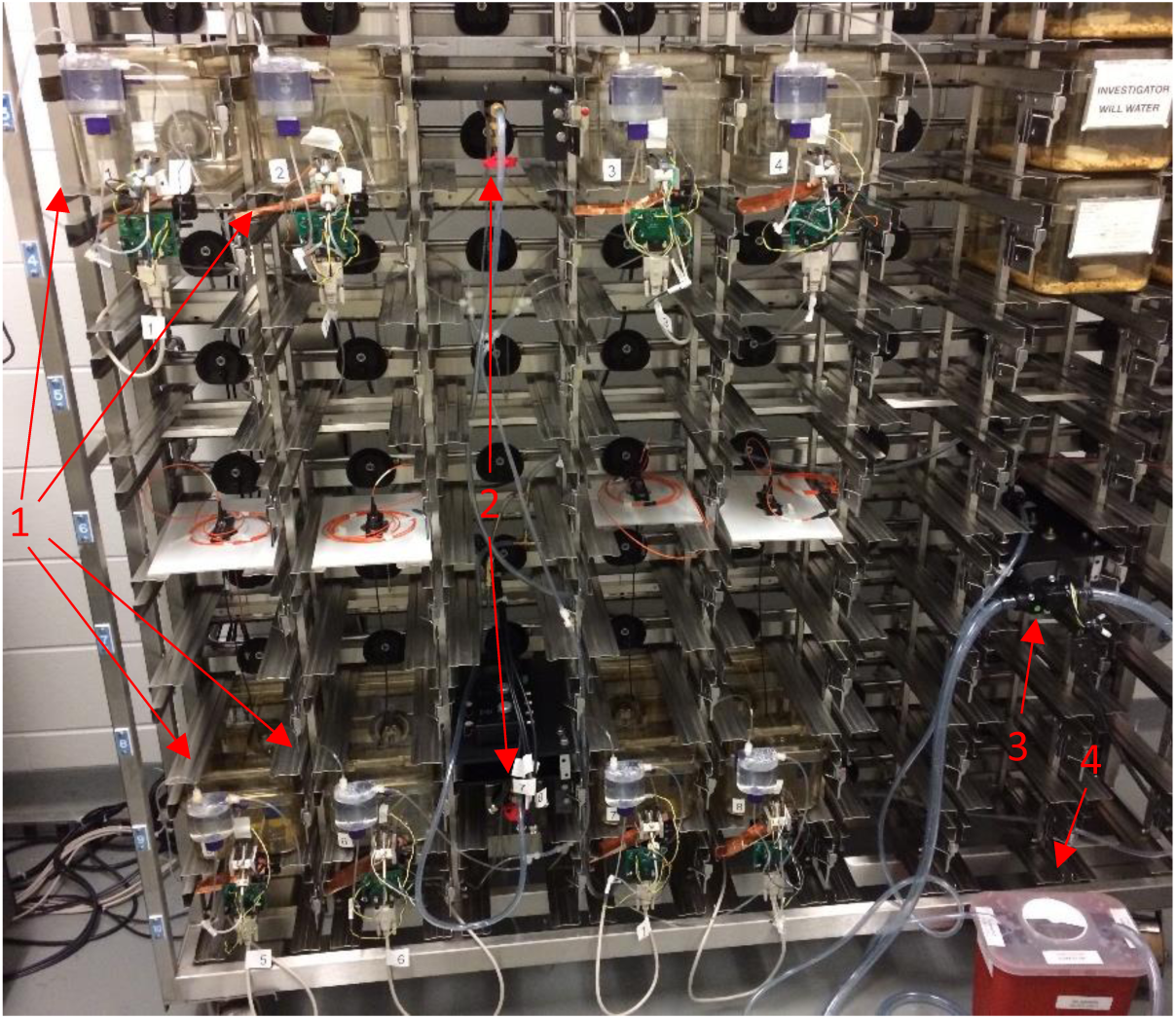
Completed water system installed on rack with 8 homecages 1. Homecages 2. Supply manifolds 3. Pumps (supply and return both mounted) 4. Main reservoir

**Figure 112.**
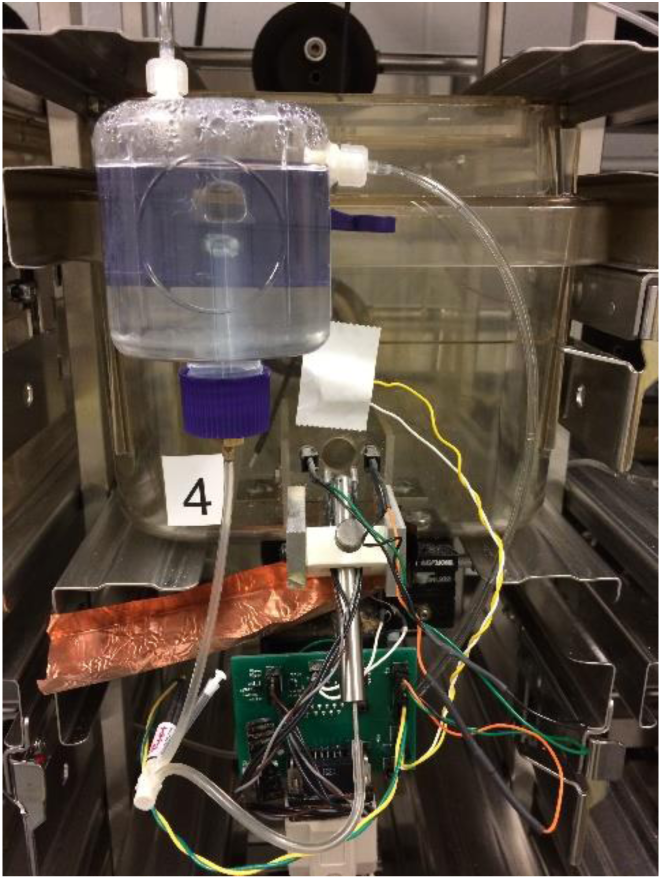
Homecage reservoir connected to supply (top fitting) and return (side fitting) tubes

#### Part XIV: Assembling the whole system

##### Full list of Materials

| # | Part Description | Manufacturer /Supplier | Manufacturer/Supplier Part Number | Datasheet/Drawing/Modification |
| --- | --- | --- | --- | --- |
| 1 | Completed homecages x8 |  | N.A. (Custom Made) | N.A. (Custom Made) |
| 2 | Completed water system |  | N.A. (Custom Made) | N.A. (Custom Made) |
| 3 | Completed DAQ system |  | N.A. (Custom Made) | N.A. (Custom Made) |
| 4 | Completed optogenetics system |  | N.A. (Custom Made) | N.A. (Custom Made) |
| 5 | Control/storage server |  |  |  |
| 6 | DB-15 cables x8 |  |  |  |
| 7 | BNC cables x8 |  |  |  |
| 8 | LabVIEW 2013 |  |  |  |
| 9 | LabVIEW control software |  |  |  |

##### Building Instructions

###### General notes

The entire system can be assembled on a standard mouse cage rack. The 8-homecage arrangement can be repeated for as many units as needed.

###### 1. Install homecages and water system on rack

###### 2. Install optogenetic system on rack

– If possible, install the optogenetic lasers so that a leak from the homecage reservoirs wouldn’t spill on the lasers.

**Figure 113.**
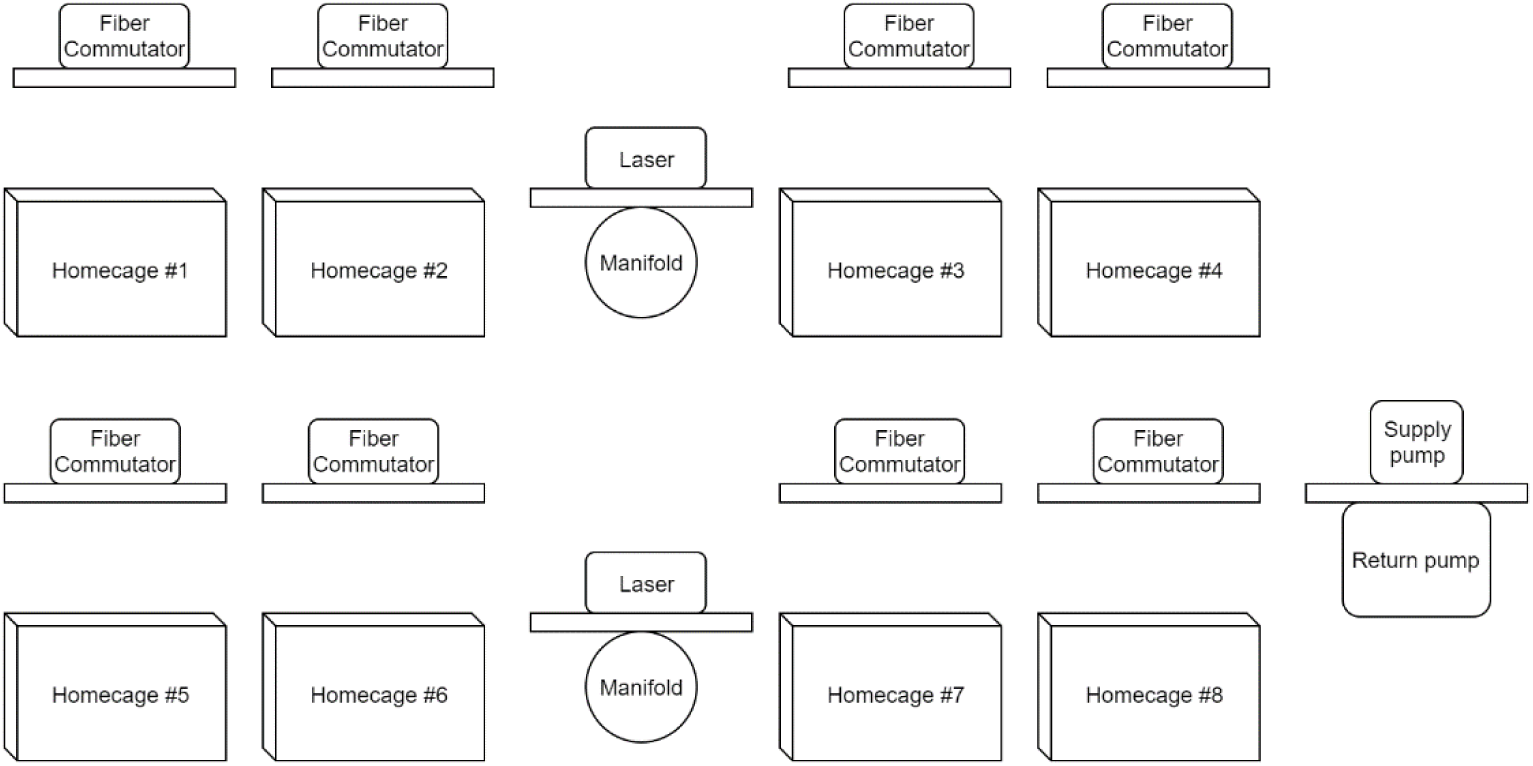
Rack arrangement for 8 homecages.

###### 3. Run ethernet cables from DAQ to network access point

– Connect each DAQ enclosure to a network access point (router or network switch) with two ethernet cables, so each sbRIO can acquire an IP address and communicate over the network.

###### 4. Run DB-15 homecages to DAQ

– Connect each homecage to the DAQ with a DB-15 cable.
– Numbering the cables, homecages, and DAQ outputs to correspond with each other is strongly recommended.

###### 5. Run BNC cables from lasers to DAQ

– Connect each laser to the DAQ with a BNC cable.

###### 6. Connect server to network access point

– Connect the server to the same network as the DAQs
– The server should be able to communicate with each sbRIO via LabVIEW.
– Note that the NI software “NI MAX” is useful for discovering the IP addresses of each sbRIO on the network.

